# Durophagous adaptations in a new crocodyliform from the Santonian of Hungary and a systematic reassessment of European Cretaceous atoposaurids

**DOI:** 10.64898/2026.10.26.755258

**Authors:** Máté Szegszárdi, Attila Ősi, Márton Rabi

## Abstract

Atoposauridae comprises a group of diminutive, semiterrestrial, brevirostrine neosuchian crocodyliforms that represents a characteristic component of European Jurassic–Cretaceous tetrapod faunas. Although frequently evaluated within the context of crown-group Crocodylia origins, their highly contested phylogenetic relationships continue to obscure their precise role in crocodylomorph evolutionary history. Particular ambiguity surrounds the monophyly of the genus *Theriosuchus*—with some hypotheses inferring long-term persistence of the lineage in Mesozoic Europe, whereas others interpreting select Cretaceous species as representatives of separate, even non-atoposaurid clades (e.g. *Sabresuchus* spp). We describe a new, stratigraphically intermediate taxon from the Santonian of Hungary based on isolated cranial elements, consistently placed as the sister taxon of *Sabresuchus (Theriosuchus) sympiestodon* from the Maastrichtian of Romania by our comprehensive set of phylogenetic analyses using three separate datasets. *Sabresuchus tribodon* sp. nov. is either recovered within Atoposauridae (global datasets) or Paralligatoridae (ingroup analysis) and can be distinguished from all other representatives of these groups by a specialized morphology consisting of massive, thick-enameled bulbous posterior maxillary teeth ornamented with apicobasal ridges, an extremely short rostrum with a reduced number of maxillary and dentary teeth, and the significantly upturned dorsal margin of the posterior dentary and surangular. Comparable anatomical adaptations are present in ecomorphologically convergent extinct crocodyliforms whose inferred feeding ecology included durophagy. However, this complete suite of characters has not been previously documented in either atoposaurids or paralligatorids. Our systematic reassessment results in an up-to-date taxonomy of Cretaceous European atoposaurids and tentatively confirms the polyphyly of *Theriosuchus*, though more definitive conclusions require combining individual, differently focused datasets. We reinterpret *Aprosuchus ghirai* from the Maastrichtian of Romania as a junior synonym of *Sabresuchus sympiestodon* and the putative European notosuchian *Ogresuchus furatus* from the Maastrichtian of Spain as an indeterminate atoposaurid.

## Introduction

The crocodyliform group Atoposauridae includes small-bodied, semiaquatic or terrestrial neosuchian taxa predominantly from the Jurassic–Cretaceous of Europe (Pochat-Cottilloux et al., 2024; Tennant et al., 2016). The history of research on atoposaurids dates back to von Meyer (1850, 1851) and the group has been often discussed in the context of the transition to Eusuchia, a clade including crown-group crocodyliforms and its immediate stem (Benton & Clark, 1988; Joffe, 1967; Martin et al. 2020; Pol et al. 2009; Salisbury et al. 2006; Tennant et al. 2016; Turner 2015). Atoposaurids traditionally contain five ‘core’ genera, *Theriosuchus, Alligatorium, Alligatorellus, Montsecosuchus* and *Atoposaurus* (Benton & Clark, 1988; Buscalioni & Sanz, 1988), but some recent phylogenies challenged the monophyly of this group (Tennant et al., 2016) leaving phylogenetic relationships and taxonomy uncertain (Martin et al., 2010, 2014; Pochat-Cottilloux et al., 2024; Schwarz et al., 2017; Tennant et al., 2016; Young et al., 2016). The most completely preserved taxa have been reported from Europe and Southeast Asia (Benton & Clark, 1988; Buscalioni & Sanz, 1988; Csiki-Sava et al., 2015; Martin et al., 2010; Owen, 1878, 1879; Pochat-Cottilloux et al. 2024; Steel, 1973; Schwarz & Salisbury, 2005; Tennant et al., 2016;), whereas more ambiguous records, mostly of isolated teeth and fragmentary cranial remains, come from the Upper Jurassic of the United States (*Theriosuchus morrisonensis* Foster, 2018) and Portugal (Atoposauridae indet.; Puértolas-Pascual, 2025), the Lower Cretaceous of China (*Theriosuchus* sp.; Wu et al., 1996) and Thailand (Atoposauridae indet.; Lauprasert et al., 2011), and the Upper Cretaceous of Cameroon (*Brillanceausuchus babouriensis* Michard et al., 1990).

Tennant et al. (2016) defined the Atoposauridae as all taxa more closely related to *Atoposaurus meyeri* von Meyer, 1850 than to *Crocodylus* Laurenti, 1768 and recognized Atoposauridae as a clade limited to the European Jurassic taxa *Atoposaurus*, *Alligatorium*, and *Alligatorellus*. However, their analyses did not include *Crocodylus* or any other extant taxa, leaving their restricted concept unjustified. In contrast, global phylogenetic analyses omit *Atoposaurus* and *Alligatorellus*, leaving the monophyly of Atoposauridae currently untested (Pochat-Cottilloux et al., 2024; Schwarz et al., 2017; Turner, 2015). Furthermore, the monophyly of the genus *Theriosuchus* remains contentious. Several studies infer an exceptionally long temporal range for this lineage, spanning from the Middle Jurassic to the latest Cretaceous, and supporting the hypothesis of late-surviving insular relict tetrapods in Europe (Csiki-Sava et al., 2015; Martin et al., 2010, 2014; Schwarz & Salisbury, 2005; Szegszárdi et al., 2026; Young et al., 2016). Conversely, other analyses resolve *Theriosuchus* as poly- or paraphyletic (Pochat-Cottilloux et al., 2024; Schwarz et al., 2017; Turner, 2015; Venczel & Codrea, 2019). In particular, the Early Cretaceous *T. ibericus* Brinkman, 1992 and the Late Cretaceous *T. sympiestodon* Martin, Rabi and Csiki, 2010 are recovered as a distinct lineage with closer affinities to Paralligatoridae, unified under the genus *Sabresuchus* (Tennant et al., 2016).

We here describe a new and stratigraphically intermediate *Theriosuchus*-like taxon from the Upper Cretaceous (Santonian) of Hungary with durophagous cranial and dental adaptations, previously unknown among atoposaurids. Including the new taxon into three separate updated taxon-character datasets, we test the monophyly of *Theriosuchus* and provide a taxonomic revision of Cretaceous European atoposaurids.

## Materials and methods

### Materials

The material belonging to the newly described atoposaurid *Sabresuchus tribodon* comes from the Santonian Csehbánya Formation, Iharkút locality in Western Hungary and is currently housed in the Hungarian Natural History Museum’s collection. For comparative anatomical purposes, we personally inspected the material of *Sabresuchus sympiestodon* housed at the Museum of Dacian and Romanian Civilization, Deva (MCDRD 134, 793) and the Laboratory of Paleontology, Faculty of Geology and Geophysics, University of Bucharest (LPB [FGGUB] R. 1781, 1782, 1945), the holotype skull of *Aprosuchus ghirai* at the Babeș-Bolyai University, Cluj-Napoca (UBB V.562/1), the material of *Theriosuchus pusillus* at the Natural History Museum, London (NHMUK 48227-8, 48282, 48282.a, and 48240.a), the holotype maxilla of *Sabresuchus ibericus* at the Institute for Geological Sciences at the Freie Universität Berlin (IPFUB 102/21.2), the holotype of *Ogresuchus furatus* at the Museu de la Conca Dellà (MCD-7149), as well as the material of *Doratodon carcharidens* in the collections of the University of Vienna (IPUW 2349/5, 2349/57) and the Hungarian Natural History Museum (MTM PAL 2024.159.1, 2013.67.1., 2013.64.1., 2014.122.1, 2013.65.1, 2013.66.1.; MTM V 2010.237.1.).

Specimens of the following taxa have been comparatively used based on publications and figures therein: *Varanosuchus sakonnakhonensis* (SM-2021-1-97/101, SM-2023-1-16, SM-2023-1-17; Pochat-Cottilloux et al., 2024), *Knoetschkesuchus langenbergensis* (DFMMh/FV 200, DFMMh/FV 605, DFMMh/FV 261, DFMMh/FV 790.12, DFMMh/FV 279, DFMMh/FV 790.11, DFMMh/FV 325; Schwarz et al., 2017), *Knoetschkesuchus guimarotae* (IPFUB Gui Croc 7308, 7709, 7701, 7403, 8103, 8109, 7507, 8044; Schwarz & Salisbury, 2005), *Theriosuchus grandinaris* (PPC-2; Lauprasert et al. 2011).

### Phylogenetic analyses

The first and second set of analyses utilized taxon-character matrices based on Szegszárdi et al. (2026) (Dataset S1) and Pochat-Cottilloux et al. (2024) (Dataset S2), both of which are in turn based on Turner (2015). In these matrices, *Sabresuchus sympiestodon* and *Aprosuchus ghirai* were completely rescored based on personal observation of the specimens (Materials section). Furthermore, characters 36, 181, 195, 199, 254, and 277 were rescored in *Varanosuchus sakonnakhonensis* based on the figures of Pochat-Cottilloux et al. (2024). In addition, to clarify the relationships of *Sabresuchus tribodon* sp. nov. with other atoposaurids, we added the taxon to the dataset of Tennant et al. (2016) densely sampling tentative atoposaurid taxa (Dataset S3), following the methodology described therein. Phylogenetic analyses were conducted using TNT v. 1.6 (Goloboff & Morales, 2023). Character and state descriptions are available in Supplementary Document S4.

In the analyses on the Szegszárdi et al. (2026) matrix, we followed their method of using the xmult=hits30 command to run multiple replications of sectorial searches, drifting, ratchet and tree fusing until 30 hits to best length are produced. As in Szegszárdi et al. (2026), we set the characters 5, 138, 277 and 281 as inactive and all characters as unordered. In the matrix based on Pochat-Cottilloux et al. (2024), the New Technology Search option was used to run all algorithms (sectorial search, drifting, ratchet, tree fusing) with each algorithm’s number of iterations increased to 100, 10 random seeds and driven search to find the minimum length 10 times, otherwise retaining default settings. A heuristic search using the Traditional Search option was also conducted with 1000 random addition sequences, followed by tree-bisection-reconnection and saving 10 trees per round (Pochat-Cottilloux et al., 2024). As in Pochat-Cottilloux et al. (2024), characters 310, 314, and 315 were set as inactive due to their ontogenetic variability, and all characters were set as unordered.

### Institutional abbreviations

DFMMh/FV, Dinosaurier-Freilichtmuseum Münchehagen/Verein zur Förderung der Niedersächsischen Paläontologie, Rehburg-Loccum, Germany; IPFUB, Institute for Geological Sciences, Freie Universität Berlin, Germany; IPUW, Institute of Paleontology, University of Vienna, Vienna, Austria; LPB (FGGUB), Laboratory of Paleontology, Faculty of Geology and Geophysics, University of Bucharest, Bucharest, Romania; MCD, Museu de la Conca Dellà, Isona, Spain; MCDRD, Museum of Dacian and Romanian Civilization, Deva, Romania; MTM, Hungarian Natural History Museum, Budapest, Hungary; NHMUK Natural History Museum London, London, UK; UBB, Babeș-Bolyai University, Cluj-Napoca, Romania; SM, Sirindhorn Museum, Kalasin, Thailand.

## Results

### Phylogenetic analysis

Our analysis using the taxon-character matrix from Szegszárdi et al. (2026) resulted in 62 trees with a length of 1771 (Fig. 1A). The new taxon from Iharkút is recovered within Atoposauridae as the sister taxon of the clade formed by *Sabresuchus sympiestodon* and ‘*Aprosuchus ghirai’* (here reinterpreted as a junior synonym of *Sabresuchus sympiestodon*), united by a dorsally robust splenial posterior to the symphysis (ch. 161, state 1; *Theriosuchus pusillus*: 0) and a ridge on the ventralmost surface of the angular (ch. 219, state 1; *T. pusillus*: ?). The sister taxon of *Sabresuchus* spp. is *Varanosuchus sakonnakhonensis*, the two sharing the following synapomorphies: the dorsal head of the quadrate articulating with the squamosal, otoccipital and prootic (ch. 47, state 0); the posterolateral end of the quadratojugal slightly overhanging the quadrate (ch. 211, state 1); and the posterior margin of the choanal groove located anteriorly on the pterygoid (ch. 318, state 0). None of these traits are known in the new taxon described here, which shares the following synapomorphies with Atoposauridae: premaxilla with little contribution to internarial bar (ch. 4, state 1); ventrally opened notch at the ventral edge of the premaxilla-maxilla contact (ch. 9, state 1); and cheek teeth constricted at the base of the crown (ch. 162, state 1).

**Figure 1.** Strict consensus trees resulting from parsimony analyses of the taxon-character matrix of **A**) Szegszárdi et al. (2026), **B**) Pochat-Cottilloux et al. (2024) and **C**) Tennant et al. (2016).

The analyses in the Pochat-Cottilloux et al. matrix recovered 689 trees with a length of 1433 (Fig. 1B). Atoposauridae and Paralligatoridae are recovered as sister taxa, sharing the synapomorphies of a narrow premaxilla anterior to the nares (ch. 5, state 0); lack of contact between nasal and lacrimal (ch. 11, state 1); the anterior and posterior portion of the jugal being approximately the same width (ch. 17, state 0); a midline ridge on the frontal and parietal (ch. 22, state 1); a ventrally exposed basisphenoid (ch. 56, state 0); septated external nares (ch. 66, state 0); and the presence of a fork in the hypapophysis (ch. 259, state 1). The new taxon is recovered as part of a monophyletic clade, distant from the type species of *Theriosuchus*, *T. pusillus*, including *Sabresuchus sympiestodon* and ‘*Aprosuchus ghirai’*. This clade is united by the following synapomorphies: articulation of the dorsal head of the quadrate with the squamosal, otoccipital and prootic (ch. 47, state 0; *T. pusillus*: 1); dorsally arched dorsal edge of the surangular (ch. 74, state 1; *T. pusillus*: 0); dorsally robust splenial posterior to the symphysis (ch. 161, state 1; *T. pusillus*: 0); weakly procumbent anterior dentary alveoli (ch. 262, state 1; *T. pusillus*: 0); groove on the parietal-squamosal border (ch. 310, state 1; *T. pusillus*: 1); and posterior margin of choanal groove located anterior on the pterygoid (ch. 318, state 0; *T. pusillus*: 1). Within this clade, the internal relationships are unresolved. *Sabresuchus* spp. (including ‘*Aprosuchus ghirai’*) form the sister clade of an *Alligatorium* + *Theriosuchus* spp. clade.

The analysis using the taxon-character matrix based on Tennant et al. (2016) resulted in 14 trees with a length of 778 with all characters unordered, and 11 trees with a length of 838 when the characters listed in Tennant et al. (2016, Appendix 1) are ordered (Fig. 1C). Each tree consistently recovers a *(Sabresuchus ibericus* (Iharkút taxon + *Sabresuchus sympiestodon))* clade. *Theriosuchus* spp. is recovered as the sister clade of the clade including *Sabresuchus* spp., *Brillanceausuchus babouriensis*, *Shamosuchus djadochtaensis* and *Wannchampsus kirpachi*. As in Tennant et al. (2016), *Knoetschkesuchus guimarotae* is recovered within the *Theriosuchus* spp. clade as the sister taxon of *T. pusillus*, which may be a result of *Knoetschkesuchus langenbergensis* not being included in the matrix. Synapomorphies common to all three species of *Sabresuchus* include a laterally positioned depression on the posterolateral surface and a medial depression on the anterolateral surface of the maxilla (Ch. 49, state 1; *T. pusillus*: 0; Ch. 50, state 1; *T. pusillus*: 0), posterior teeth in a confluent dental groove (Ch. 60, state 0; *T. pusillus*: 1), the ventral edge of the squamosal groove for the upper ear lid being positioned directly beneath the dorsal edge (Ch. 136, state 1; *T. pusillus*: 0), the oblique ridge on the dorsal surface of the squamosal being located posterior to the supratemporal fenestra (Ch. 146, state 0; not preserved in the Iharkút taxon and *T. pusillus*), a shallow and anteriorly tapering dentary symphysis (Ch. 206, state 0; *T. pusillus*: 1), the transition of alveolar morphology from circular to oval in the tooth row (Ch. 218, state 1; *T. pusillus*: 1), narrower roots than crowns of maxillary teeth (Ch. 252, state 0; *T. pusillus*: 1), and the presence of low-crowned and labiolingually compressed maxillary teeth that are mesiodistally wider than apicobasally tall (Ch. 254, state 0; *T. pusillus*: 0). Four synapomorphies are furthermore shared by the Iharkút taxon and *S. sympiestodon*: a mandibular symphysis terminating posteriorly at the level of the 5-6^th^ alveoli (Ch. 204, state 1), a vertically oriented, continuous lateral dentary surface (Ch. 213, state 0), the presence of a diastema between the 7^th^ and 8^th^ dentary teeth (Ch. 225, state 0), and striations on the labial and lingual faces of the maxillary teeth (Ch. 250, state 1). The autapomorphies of *S. ibericus* are a ridged maxillary palatal surface (Ch. 57, state 2), middle maxillary teeth in a confluent dental groove (Ch. 59, state 0), a depressed dorsal surface of the parietal compared to the squamosal (Ch. 115, state 2), a narrower cranial table that still covers more than half of the mediolateral region of the quadrates (Ch. 133, state 1, absent in the Iharkút taxon), a lack of sculpting on the lateral surface of the dentary below the tooth row (Ch. 216, state 0), dentary teeth all located in a confluent dental groove (Ch. 217, state 2), inter-alveolar septa absent within anterior dental arcade of dentary (Ch. 228, state 1), posteroventrally directed hypertrophied 5^th^ tooth (Ch. 259, state 0), and dorsally inset 6^th^ and 7^th^ maxillary teeth. The autapomorphies of the Iharkút taxon are the presence of bulbous tribodont maxillary teeth (ch. 261, state 1), while those of *S. sympiestodon* are flat internal dentary alveolar margins confluent with the occlusal dentary surface (ch. 223, state 1) and a parallel dentary symphysis and dental arcade in dorsoventral view (ch. 229, state 0). The synapomorphies of the clade containing *W. kirpachi, S. djadochtaenis, B. babouriensis* and *Sabresuchus* spp. are a straight lateral contour of the snout (ch. 6, state 0), the lacrimal forming a point contact with the nasal (ch. 72, state 1), the prefrontal and lacrimal forming an elevated rim around the orbits (ch. 94, state 1), a dorsal anteroposterior ridge occupying the entire length of the frontal (ch. 101, state 3), the presence of the parietal-postorbital suture within the supratemporal fossa (ch. 113, state 1), a transversely flattened postorbital bar (ch. 131, state 0), the absence of an obliquely-oriented ridge on the dorsal surface of the squamosal (ch. 145, state 0), restricted posterodorsal exposure of the squamosal (ch. 197, state 1), ‘semi-procoelous’ (sensu Tennant et al., 2016) vertebrae (ch. 265, state 1), procoelous dorsal vertebrae (ch. 271, state 1), and articular anterior processes present on dorsal osteoderms as a discrete convexity on the anterior margin (ch. 310, state 0).

### Systematic palaeontology

Suborder **Crocodyliformes** Hay, 1930

Clade **Mesoeucrocodylia** Whetstone and Whybrow, 1983

Clade **Neosuchia** Benton and Clark, 1988

Family **Atoposauridae** Gervais, 1871

Genus ***Sabresuchus*** Tennant et al. 2016

#### Included species

*Sabresuchus ibericus* (type species), *Sabresuchus sympiestodon*, *Sabresuchus tribodon* sp. nov.

#### Revised diagnosis

Small-sized, short-snouted crocodyliforms united by the following synapomorphies: (a), laterally positioned depression on the posterolateral surface and medial depression on the anterolateral surface of the maxilla; (b), posterior maxillary teeth in a confluent dental groove; (c), ventral edge of squamosal groove for the upper ear lid positioned directly beneath the dorsal edge; (d), shallow, anteriorly tapering dentary symphysis; (e), from anterior to posterior transition of alveolar morphology from circular to oval in the tooth row; (f), narrower roots than crowns of maxillary teeth; (g), presence of low-crowned and labiolingually compressed maxillary teeth mesiodistally wider than apicobasally tall; (h), dentary with sinusoidal dorsolateral edge.

#### Comments

Tennant et al. (2016) furthermore diagnosed *Sabresuchus* on the basis of the following synapomorphies: squamosal posterolateral process (‘lobe’) offset from the dorsally flat skull table; squamosal with an oblique ridge on the dorsal surface(…), posterior to the posterior margin of the supratemporal fenestra (unknown in *S. tribodon* sp. nov.); lateral surface of dentary with concavity for reception of enlarged maxillary tooth; and a hypertrophied maxillary tooth 5, at least 4.0 times the size of adjacent maxillary teeth. It should be noted that the ‘lobe’ on the posterolateral process of the squamosal is sometimes recovered as a synapomorphy of a more inclusive clade uniting Atoposauridae + Paralligatoridae (e.g. Pochat-Cottilloux et al., 2024), while the size of the 5^th^ maxillary tooth in *S. sympiestodon* (consistent with the relative alveoli size in *S. tribodon* sp. nov.) is less than 4 times the size of the adjacent maxillary teeth.

> Species **Sabresuchus tribodon** sp. nov. (Figs 2-6)

**Figure 2.** Rostral elements of *Sabresuchus tribodon* sp. nov. from the Santonian of Iharkút, Hungary. **A-D**: Premaxilla MTM VER 2016.46. in lateral (**A, B**) and medial (**C, D**) view. **E-N**: maxilla MTM V 2010.235.1. Maxilla in lateral view (**E, F**), maxilla in medial view (**G, H**), details of 3rd tooth (**I**), maxilla in ventral (**J, K**), dorsal (**L**) and anterior view (**M, N**). **Abbreviations**: **con j**, contact surface with jugal; **con mx**, contact surface with maxilla; **con n**, contact surface with nasal; **en**, external nares; **fo**, foramina; **no**, notch; **op**, occlusal pit; **ppr**, palatal process; **sof**, suborbital fenestra. Scale bars: A-H, J-N = 10 mm; I = 1 mm. Photographs E-M and illustration I by Nóra Sebők.

> *LSID.* (To be added upon acceptance)

#### Derivation of name

The specific name *tribodon* translates from Greek *τρίβω*, to grind, and *οδον*, tooth in reference to its massive, bulbous posterior crushing teeth, a unique adaptation to a durophagous feeding ecology among Atoposauridae.

#### Type material

MTM V 2010.236.1., right maxilla with in-situ teeth.

#### Type locality

Németbánya II bauxite lens, Iharkút, Bakony Mountains, Hungary; Csehbánya Formation, Santonian, Upper Cretaceous.

#### Referred material and range

MTM VER 2016.46., right premaxilla; MTM V 2010.235.1., fragmentary right maxilla; MTM VER 2026.28, left mandible; MTM VER 2026.27, right mandible; MTM VER 2026.29, fragmentary frontal; MTM VER 2016.4, fragmentary skull roof; MTM PAL 2014.124.1-12., isolated teeth (type locality); Ajka Coal Formation, Santonian, Jókai Mine (Ősi et al., 2016).

#### Diagnosis

A species of *Sabresuchus* based on preserved traits of the diagnosis of the genus (see above). *Sabresuchus tribodon* differs from all described atoposaurid species in the presence of massive, bulbous posterior crushing teeth that are wider than tall, labiolingually compressed and basally constricted on the crown-root boundary with small, pointed apices, no carinae, and a strongly ornamented enamel surface with prominent apicobasal ridges that do not reach the constricted base of the crown mesially and distally, nor in the middle in some cases; and a reduced number of maxillary and dentary teeth (9 maxillary, 13 dentary). Further differs from *Sabresuchus sympiestodon* in possessing a total of nine maxillary teeth (10-11 in *S. sympiestodon*); the suture for the jugal on the maxilla reaching the level of the 7th-8th tooth anteriorly (6th-7th tooth in *S. sympiestodon*); suborbital fenestra reaching the 7-8th maxillary tooth (5-6th in *S. sympiestodon*); and a consequently shorter rostrum. In addition, the adult size of *S. sympiestodon* (MCDRD 793, Martin et al., 2014) is inferred to be about 1.4 times the size of the largest known specimen of *S. tribodon*. An orbitonasal maxillary sulcus is not apparent in *S. tribodon* at the location where it is present in *S. sympiestodon*. Differs from ‘*Aprosuchus ghirai*’ (*Sabresuchus sympiestodon*) in a markedly pronounced dorsal expansion of the dentary and the surangular. Differs from the wannchampsid (sensu Allen et al., 2025) taxa *Wannchampsus kirpachi* Adams, 2014 in lacking true ziphodont teeth and possessing a prominently dorsally expanded surangular and posterior portion of the dentary; from *Thikarisuchus xenodentes* Allen, Wilberg, Turner and Varricchio, 2025 in the proportionally higher anterior half and distinctly tapering posterior half of the maxilla; nine maxillary teeth instead of 11 in *T. xenodentes*; a rounder, more bulbous 8th maxillary tooth; posterior maxillary teeth possessing pointed apices and a constricted root and crown border; apicobasal ridges of crown do not reach the constricted lower crown; a narrower and dorsally shorter frontoparietal ridge lacking the boss present near the frontal-parietal suture in *T. xenodentes*; a proportionally much shorter mandible with an upturned, dorsally expanded posterior portion of the dentary and surangular; a dorsally expanded alveolar margin of the 10th dentary tooth, with a corresponding prominent notch on the maxilla. *S. tribodon* further differs from both *W. kirpachi* and *T. xenodentes* in possessing a shorter, more rounded and more posteriorly directed posterolateral squamosal lobe. Differs from the paralligatorid *Shamosuchus djadochtaensis* (Mook, 1924; Pol et al., 2009; Turner, 2015) in possessing a much more prominent maxillary notch posterior to the 5th maxillary tooth; an enlarged 5th dentary tooth and corresponding notch on the ventral margin of the premaxilla-maxilla suture; expansion of the dorsal dentary margin at the level of the 10^th^ dentary tooth; and a nearly straight ventral margin of the mandible. Differs from the paralligatorid *Scolomastax sahlsteini* Noto, Drumheller, Adams and Turner, 2020 in the number of dentary tooth positions (13 in *Sa. tribodon*, 10-11 in *Sc. sahlsteini*); the 3rd alveolus being smaller than the 4th (lacking the enlarged 3rd alveolus present in *Sc. sahlsteini*); a dorsoventrally narrower dentary and splenial posterior to the symphysis and anterior to the dorsally expanding posterior region; an anteriorly less extended splenial reaching the 6th tooth anteriorly (anteriormost point of the splenial between the 4th and 5th tooth in *Sc. sahlsteini*); and a nearly horizontal ventral margin of the dentary and angular. Differs from the putative paralligatorid *Tarsomordeo winkleri* Adams, 2019 in the expansion of the dorsal dentary margin at the level of the 10^th^ dentary tooth and posterior to the toothrow; and the lower number of dentary tooth positions (13 in *S. tribodon*, 16 in *T. winkleri*).

#### Remarks

The taxonomic integrity of the holotype and the referred material is justified by distinct dental traits (shared between in situ and isolated teeth), skull ornamentation, and skull proportion in the context of the local and regional crocodyliform fauna (Delfino et al., 2008; Martin et al., 2010, 2014; Ősi, 2008; Ősi & Rabi, 2006; Ősi et al., 2012; Rabi & Delfino, 2012; Rabi & Sebők, 2015; Szegszárdi et al. 2026). We furthermore refer isolated teeth previously reported as cf. *Theriosuchus* sp. from the coeval Ajka Coal Formation (heterotypic facies of the Csehbánya Formation) to this taxon.

#### Description

##### Rostrum

###### Premaxilla

The premaxilla of *Sabresuchus tribodon* is represented by VER 2016.46. (Fig. 2A-D). The bone is crescent-shaped, with a slightly convex outer margin. Unlike that of *Theriosuchus pusillus*, but similarly to the condition in *Knoetschkesuchus* spp. and *Ogresuchus furatus*, the premaxilla of *S. tribodon* is taller than wide. Based on the preserved portion of the palatal process, the premaxilla was oriented subvertically in life similarly to *O. furatus* and *Knoetschkesuchus langenbergensis*. The outer ornamentation is characterized by fine, subcircular pitting. Three alveoli are preserved in the specimen, although the anteriormost section of the tooth row is damaged. Posterior to the third preserved alveolus, the ventral margin of the premaxilla is upturned, forming the anterior margin of a notch in the tooth row, presumably at the premaxilla-maxilla contact like in *Knoetschkesuchus guimarotae, Knoetschkesuchus langenbergensis, Theriosuchus pusillus, Theriosuchus grandinaris* and *Ogresuchus furatus*. No in-situ teeth are present. A large, anterodorsally oriented perinarial fossa is present lateral to the external nares. The surface of the perinarial fossa is unsculpted, slightly concave and extends anteroventrally towards the anterior margin of the premaxilla. The suture of the premaxilla with the nasal runs anteroventrally along the downturned, convex dorsal edge of the premaxilla and ends separated from the external nares by a notch like in *O. furatus* and *T. pusillus*. The premaxillary palate is only partially preserved, bearing a row of minute neurovascular foramina medial to the tooth row. Based on the insertion surface present in MTM V 2010.235.1., the premaxilla presumably contacted the maxilla posterolaterally like in *Knoetschkesuchus* spp. and *Theriosuchus pusillus*.

###### Maxilla

The maxilla of *Sabresuchus tribodon* is represented by specimens MTM V 2010.235.1. (Fig. 2E-N) and MTM V 2010.236.1. (Fig. 3, holotype). MTM V 2010.236.1. is preserved almost completely, only missing a fragment of the anterior border and part of the palatal process, reaching an anteroposterior length of c. 44 mm. MTM V 2010.235.1. is proportionately smaller and more incomplete, being broken posterior to the 8^th^ tooth position, as well as missing the dorsal margin of the anterior section and part of the palatal process. Nine tooth positions in total are present in the maxilla, of which the anterior six teeth are located in isolated alveoli, and the posterior three in a septated dental groove with a ventral margin slightly constricted between each two tooth positions. The ventral margin of the maxilla is sinusoidal with a convex anterior and posterior half separated by the concave notch posterior to the 5^th^ tooth. Medial to the notch, an occlusal pit is present like in *S. sympiestodon*. As is typical for most atoposaurids, the 4^th^ tooth position is occupied by a caniniform tooth. Near the caniniform tooth, the lateral surface of the maxilla is convex, and the surface medial to the tooth row is significantly evaginated. Small, subcircular nutrient foramina are present medially to each tooth position like in *Sabresuchus sympiestodon,* but much smaller in size than the occlusal pits of *D. carcharidens*.

**Figure 3.** MTM V 2010.236.1., holotype maxilla of *Sabresuchus tribodon* sp. nov. from the Santonian of Iharkút, Hungary. **A-B**, maxilla in lateral view. **C**, close-up of 7^th^ to 9^th^ maxillary teeth in lateral view. **D**, SEM photograph of 9^th^ tooth in lateral view. **E-F**, maxilla in medial view. **G**, close-up of 7^th^ to 9^th^ teeth in medial view. **H-I**, 7^th^ to 9^th^ teeth in occlusal view. **Abbreviations: aof,** antorbital fenestra; **con j,** contact surface with jugal; **con pmx**, contact surface with premaxilla; **no,** notch**; pproc,** palatal process. Scale bars: A, B, E, F = 10 mm; C, G, H, I = 2 mm; D = 1 mm. Photographs A-H by Nóra Sebők.

The anterior end of the toothrow and the ventral edge of the maxilla are turned upwards, forming the posterior end of the notch on the premaxilla-maxilla border. The outer surface of both specimens is finely sculpted with shallow, minute circular and oval pits. The orbitonasal sulcus present in *Sabresuchus sympiestodon* (Martin et al., 2010, 2014) and some specimens of *T. pusillus* (e.g. B.M.(N.H.) Palaeont. Dept. 48330) is absent from *S. tribodon*. Anterior to the ventral notch posterior to the 5^th^ tooth, the maxilla is dorsoventrally much broader than in the dorsoventrally narrow, tapering posterior section. In dorsal view, the maxilla is mostly straight anteriorly and slightly laterally deflected posteriorly, with the deflection being more apparent in MTM V 2010.235.1. Compared to *Sabresuchus sympiestodon*, the dorsal and ventral halves of the lateral surface anterior to the notch are more distinct, with a thick, slightly convex ventral and a thin, slightly concave dorsal half especially apparent in MTM V 2010.236.1., similarly to the condition in *Theriosuchus pusillus* and *Ogresuchus furatus*. The anterior section of MTM V 2010.235.1. bears an anterodorsal depression which is absent from MTM V 2010.236.1. While the antorbital fenestra itself is not preserved in either of the maxillary specimens, a smooth depression enclosed by an elevated rim is present on the medial surface of MTM V 2010.235.1., which is similar to the surface anteriorly bordering the antorbital fenestra in *Doratodon carcharidens* (Szegszárdi et al. 2026, fig. 2C-D) and *Sabresuchus ibericus* (Brinkmann 1992). This surface is also present in specimens of *Sabresuchus sympiestodon* (LPB (FGGUB) R. 1782, R. 1945). Anteriorly to the contact surface with the jugal, a medially deflected line on the dorsal surface marks the suture with the lacrimal. The suture for the nasal is preserved in MTM 236.1. on the dorsomedial surface of the anterior portion and is deflected slightly downward anteriorly like in *S. sympiestodon*. The suture with the premaxilla is partially preserved on the anterodorsal surface, in addition to a smooth, concave surface anterior to the first tooth position in V 2010.235.1. representing the insertion surface.

Based on the angle of the palatal process, the maxilla was positioned subvertically in the rostrum. Although most of the palate is missing, the palatal process is visibly located more dorsally compared to the toothrow in the anterior half and closer, more ventrally in the posterior half of the maxilla, suggesting that the palate was slightly deflected ventrally. At the level of the notch posterior to the 5^th^ tooth, the palate bears a large, semicircular depression. The suborbital fenestra reaches the level of the 7^th^ tooth anteriorly, its lateral wall running parallel to the lateral edge of the maxilla. In contrast, the suborbital fenestra reaches the 5^th^ to 6^th^ tooth in *Sabresuchus sympiestodon*. The posterior end of both maxillary specimens preserves an anterodorsally tapering, slightly striated contact surface for the anterior portion of the jugal, extending up to the level of the 7^th^ to 8^th^ tooth position, unlike *Sabresuchus sympiestodon* in which it reaches the 5^th^ to 6^th^ tooth position.

##### Skull roof

###### Frontal

The frontal of *Sabresuchus tribodon* is represented by the partial specimen MTM VER 2026.29 (Fig. 4A-D), which is missing its anteriormost section including the contact surface with the nasal. The narrow interorbital section with a sudden mediolateral expansion at a sharp angle posteriorly, differentiates the specimen from all sympatric crocodyliforms. The dorsal interorbital surface is ornamented by subcircular pits, becoming a mediolaterally extended oval shape near the orbital margin. The anterior end of the parasagittal midline ridge characteristic for atoposaurid and paralligatorid taxa is visible on the posterior half of the dorsal surface. The orbital margin is unsculpted and elevated dorsally with only a slight lateral expansion, similarly to *Theriosuchus pusillus* and *Alligatorellus beaumonti*, but contrasting the dorsally lower and laterally more expanded orbital margin of *Sabresuchus sympiestodon* and the sympatric *Doratodon carcharidens*. A row of minute foramina is present ventrally to the elevated rim, on the lateral surface of the frontal. A minute portion of the anteromedial margin of the supratemporal fenestra is visible on the posterior end of the dorsal surface, near the contact with the postorbital and parietal. The parieto-postorbital suture is located more anteriorly compared to the supratemporal fenestra than in *S. sympiestodon*. The ventral surface bears a prominent ridge, confluent with the lateral surface, on each side of the anterior half of the specimen, bordering the orbits medially and the channel for the olfactory lobes laterally. Posteriorly at the level of the lateral expansion of the frontal, these ridges are replaced by a ventrally facing, oval flat surface on each side, possibly forming the contact surface for the laterosphenoid (Iordansky, 1973). The contact surfaces with the postorbitals and parietal are visible on the posterior end of the ventral surface, located laterally and medially respectively.

**Figure 4.** A-D. MTM VER 2026.29, partial frontal of *Sabresuchus tribodon* sp. nov. from the Santonian of Iharkút, Hungary in dorsal (**A**, **C**) and ventral (**B**, **D**) view. **E-F**. partial skull roof MTM VER 2016.4. in dorsal view. **Abbreviations**: **con fr**, contact with frontal; **con lsp**, contact with laterosphenoid; **con p**, contact with parietal; **con po**, contact with postorbital; **dgr**, dorsal groove of squamosal; **dl. proc**, dorsolateral process of squamosal; **fo**, foramina; **lgr**, lateral groove of squamosal; **o**, orbit; **p**, parietal; **pr**, parietal ridge; **sq**, squamosal; **sql**, squamosal lobe; **stf**, supratemporal fenestra. Scale bars = 10 mm.

###### Parietal

The parietal of *Sabresuchus tribodon* is partially preserved in the specimen MTM VER 2016.4 (Fig. 4E-F). Its ornamentation is characterized by small, subcircular pits which are reduced in size near the posteromedial border of the supratemporal fenestra. The medial border of the supratemporal fenestra is smooth and slightly inset from the rest of the skull roof, while the medial wall of the fenestra formed by the parietal tapers posteriorly. The medial wall of the supratemporal fenestra descends at a steeper angle and extends less into the fenestra than that of ‘*Aprosuchus ghirai’*. The parietal bears a sagittal midline crest, a characteristic for atoposaurids and paralligatorids. Similarly to *Sabresuchus sympiestodon*, the pits retain their circular shape in the vicinity of the crest. In contrast, the ornamentation of the sympatric *Doratodon carcharidens* is characterized by more oval, elongated pits medially on the skull roof and near the midline crest. The ridge running anteroposteriorly from the medial wall of the supratemporal fenestra to the posterior end of the parietal-squamosal suture present in both *T. pusillus* and ‘*A. ghirai’* is absent from *S. tribodon*. The posterior margin of the parietal is downturned, posteriorly oriented and smooth, forming part of the occipital surface.

###### Squamosal

The squamosal of *Sabresuchus tribodon* is represented by fragmentary remains assigned to the partial skull roof MTM VER 2016.4 (Fig. 4E-F). The dorsal surface of the squamosal is ornamented by circular and subcircular pits. Unlike *D. carcharidens, ‘Aprosuchus ghirai’* and *Sabresuchus sympiestodon* but similarly to *Alligatorellus beaumonti*, the lateral margin is separated from the main body by a groove, and decorated with sparser, significantly smaller and shallower circular and oval pits. Consistently with the parietal, the lateral border of the supratemporal fenestra formed by the squamosal is slightly depressed and ornamented with pits. The dorsal ridge running anteroposteriorly on the squamosal of *D. carcharidens* is not present in this taxon. The lateral edge of the squamosal possesses a posteriorly widening lateral notch (sensu Schwarz et al., 2017) running anteroposteriorly, separating the dorsal surface from the lobe supporting the outer ear musculature. The posterolateral margin bears an ‘unsculpted’ lobe (sensu Turner, 2015) characteristic for Atoposauridae and Paralligatoridae, although in the case of *S. tribodon* and ‘*Aprosuchus ghirai’*, the lobe bears some slight sculpturing characterized by diminutive, sparse circular pits. The lobe is significantly shorter, thicker and more rounded than the extended, rectangular lobe of *Doratodon carcharidens,* instead resembling that of *Knoetschkesuchus langenbergensis* and *Sabresuchus sympiestodon*. While the lobe of ‘*A. ghirai’* is also short proportionately, the posterior end of the lobe is more rectangular, featuring a straight posterior margin with nearly perpendicular corners. The medial surface of the lobe is smooth and confluent with the medial surface of the main body of the squamosal, while the dorsal surface of the lobe is separated from that of the main body by a groove similarly to the condition in ‘*Aprosuchus ghirai’*. Although mostly smooth, the anterior and posterior rims of the convex dorsal surface preserve one row of small neurovascular foramina each. The ventral surface is smooth and concave, and bears thin, parallel anteroposteriorly running scars on the posterior edge.

##### Mandible

###### Form and proportions

The tooth-bearing region prior to the dorsal expansion of the posterior dentary and the surangular of *Sabresuchus tribodon* is relatively short, comprising about half to two thirds the total length of the mandible (Fig. 5), similarly to the condition in *Iharkutosuchus makadii, Caiman latirostris and Varanosuchus sakonnakhonensis*, although to a lesser extent than in highly specialized notosuchians such as *Simosuchus clarki* or *Libycosuchus brevirostris*. The mandible of ‘*Aprosuchus ghirai’* is characterized by a similar rostrum-to-posterior ratio, although the anterior end of the rostrum is eroded, making the proportions in life uncertain. The surangular and angular are notably expanded dorsally, similarly to the condition in *L. brevirostris*, *Scolomastax sahlsteini* and *Allognathosuchus haupti*, as well as ‘*Aprosuchus ghirai’* and *V. sakonnakhonensis* among atoposaurids. The dentary portion of the mandible appears proportionately shorter in MTM VER 2026.27 due to taphonomic compression lateral to the 11^th^-13^th^ tooth positions, resulting in a shortening of about 4 mm. A mandibular fenestra may have been present based on parts of the posteroventral margin of the dentary and the dorsal margin of the angular, but its presence cannot be inferred for certain due to the preservation of the specimens.

**Figure 5.** Dentary specimens of *Sabresuchus tribodon* from the Santonian Iharkút site, Hungary. A-F: MTM VER 2026.27 mandible in lateral (**A, B**), medial (**C, D**) and occlusal (**E, F**) view. G-L: MTM VER 2016. 92. dentary fragment in anterior (**G, H**), ventrolateral (**I, J**) and dorsomedial (**K, L**) view. M-P: MTM VER 2026.28 mandible in medial (**M, N**) and lateral (**O, P**) view. **Abbreviations**: **an**, angular; **anp**, posterior process of angular; **d**, dentary; **fo**, foramen; **fio**, foramen intermandibularis oralis; **sa**, surangular; **sp**, splenial; **sym**, symphysis. Scale bars = 10 mm.

**Figure 6.** Isolated anterior (**A-E**) and posterior (**F-L**) teeth belonging to *Sabresuchus tribodon* from the Santonian of Iharkút, Hungary, each in lingual, labial and occlusal view. Photographs by Nóra Sebők. Scale bars = 1 mm.

###### Dentary

13 tooth positions are present in the dentary of *Sabresuchus tribodon*, although the dorsal tooth-bearing margin posterior to the symphyseal region in MTM VER 2026.28 and the anterior end of the symphyseal region of MTM VER 2026.27 are poorly preserved. The 5^th^ alveolus is enlarged, preserving a caniniform tooth in MTM VER 2026.28. The dorsal margin of the rostral portion of the dentary is sinusoidal, with a dorsal expansion near the 5^th^ tooth in the symphyseal region corresponding to the notch in the ventral margin premaxilla-maxilla boundary, and another dorsal expansion at the 10th tooth corresponding to the notch on the ventral maxillary margin posterior to the 5^th^ maxillary tooth. Posterior to the last alveolus, the dentary is upturned at a 135-degree angle as the posterior section of the mandible expands dorsally. The symphyseal contact extends posteriorly until the 6^th^ to 7^th^ alveolus in contrast to the 5^th^ in *Sabresuchus ibericus*. The symphysis tapers anteriorly as seen in MTM 2106_128_447, forming a V shape if both mandibles were present. Similarly to *Sabresuchus sympiestodon* but unlike *S. ibericus*, circular nutrient foramina of varying size are present in the symphyseal region medial to the tooth row. The dorsal surface of the symphysis bears a medial semicircular depression in MTM VER 2026.28, which may be an artifact of preservation as it is missing from MTM VER 2026.27. A tooth from the symphyseal region, possibly the 6th in the tooth row, is preserved in MTM VER 2016.92. It is projected labially with the apicobasal axis forming a c. 45° angle with the symphyseal surface. The base of the tooth is surrounded by an evagination of the dorsal surface with a pair of neurovascular foramina located anteromedial to the tooth. Lateral to the 7^th^ tooth position, the lateral surface of the dentary bears a distinct concave evagination, presumably for the upper caniniform. In this region, the lateral sculpting is much less prominent, and a neurovascular foramen is present on the dorsolateral surface, below the alveoli. The outer surface of the dentary is delicately sculpted. The ornamentation is characterized by small, circular pits in the lateral surface of the symphyseal region, and more elongated, oval pits on the ventral surface of the symphyseal region as well as on the posterior section of the dentary, with a thin, unsculpted dorsal margin starting from the evagination. The narrow Meckelian groove is visible in MTM VER 2026.28, with a nearly straight dorsal and ventral margin and thin dorsal and ventral walls formed by the dentary.

###### Splenial

The splenial of *Sabresuchus tribodon* is preserved in MTM VER 2026.27. It forms the posteromedial part of the symphysis and borders the dentary medially posterior to the symphyseal region. The suture of the two bones is directed in an anterodorsal-posteroventral direction, the splenial forming half of the anteroposterior length of the preserved portion of the symphysis dorsally and about a third ventrally. Although much of the symphysis is not preserved in ‘*Aprosuchus ghirai’*, the splenial is less involved in the formation of the symphyseal region than in *S. tribodon*. The splenial is widest mediolaterally in the symphyseal region and tapers posteriorly. The dorsal and medial surfaces are separated by a longitudinal ridge running along the entire preserved length of the splenial, which is continuous with the dorsal surface of the symphyseal region. While a similar ridge is present on the ventral edge of the medial surface, its posterior extent is unknown due to the incomplete preservation of the splenial. Immediately posterior to the symphyseal region, the two ridges border a depression bearing an oval neurovascular foramen (foramen intermandibularis oralis, Noto et al. 2020) on the medial surface. This foramen is present in *Theriosuchus pusillus* but absent in ‘*A. ghirai’*. The ventral outer surface of the splenial is lightly ornamented with large, elongated pits often fused together and bordered by thin, diminutive edges similarly to ‘*A. ghirai’, T. pusillus*, *Theriosuchus grandinaris*, *Knoetschkesuchus guimarotae*, but also with species of *Shamosuchus* (Tennant et al., 2016).

###### Angular

The angular of *Sabresuchus tribodon* is preserved in MTM VER 2026.27 and MTM VER 2026.28. The lateral wall is high and ornamented by pits like in *Knoetschkesuchus guimarotae* but bears a distinct ventral margin and a more inset dorsal portion like in *Knoetschkesuchus langenbergensis.* In contrast to *Knoetschkesuchus* spp. and more similarly to ‘*Aprosuchus ghirai’*, the ventral margin is ornamented with elongated grooves. The ventral surface of the angular is smooth and slightly concave on the posterior end, and anteriorly expands in a dorsomedial direction, wrapping around to form a continuous medial surface with the splenial like in ‘*A. ghirai’*. The bone bears an unsculpted posterior process angled posteroventrally, similarly to ‘*A. ghirai’* and *Knoetschkesuchus* spp. This surface may serve as the attachment surface for a prominent M. pterygoideus ventralis muscle, similarly to the condition in the durophagous neosuchian *Allognathosuchus haupti* (Ősi, 2014).

###### Surangular

The surangular borders the dentary dorsomedially, forming a convex anterodorsal edge posterior to the concave upturned portion of the dentary. This condition is similar to that present in the durophagous neosuchians *Allognathosuchus haupti, Bernissartia fagesii* and *Caiman latirostris*. The surangular contacts the angular dorsally, as well as on the posterior margin of the mandible via a downturned process, similarly to ‘*Aprosuchus ghirai’* and *Theriosuchus pusillus*. The posterior margin of the process is nearly subvertical, at a c. 125° angle with the posteriorly inclined posterodorsal edge of the surangular. The dorsal edge of the surangular is arched dorsally, like in *Acynodon* spp., *Scolomastax sahlsteini, Varanosuchus sakonnakhonensis* and *‘Aprosuchus ghirai’*.

##### Dentition

###### General

In total, at least 3 premaxillary, 9 maxillary and 13 dentary tooth positions are present in *Sabresuchus tribodon,* although the premaxilla is anteriorly incomplete. The dentition is strongly heterodont. In both the maxillary and the dentary dentition, two distinct tooth morphotypes are present in the tooth row (Fig. 6).

Teeth belonging to the anterior morphotype include the 3^rd^ tooth in MTM V 2010.235.1., the 4^th^ tooth in MTM V 128_447 and the 10^th^ tooth in MTM VER 2026.27. The tooth crown is tall and lanceolate, slightly labiolingually flattened, and nearly symmetrical mesiodistally. The apicobasal axis is curved, with the apex slightly deflected lingually. The crown is constricted basally and has an oval cross-section. The teeth bear a mesial and distal carina which lack ornamentation or denticles and are widening and ‘shelf-like’ in medial view. The labial surface of the teeth is convex, while the lingual surface bears one concavity mesiodistally at the level of each carina. On some teeth, a lingual concavity is visible, resulting in a bean- or boomerang-shaped cross-section. The enamel is ornamented by striae which run parallel to the apicobasal axis and often enter the self-like part of the carinae but do not reach the margin of the carinae (pseudoziphodonty). These teeth are distinguished from the coeval *Allodaposuchus* sp. by the presence of well-developed, shelf-like carinae. Small wear facets are visible on some specimens, either restricted to the apex, or in teeth with more well-developed carinae, apically and along the edges of the carinae.

Teeth belonging to the posterior morphotype are massive and bulbous, wider than tall, labiolingually strongly flattened and bear strong apicobasal ridges along the entire height of the crown. The posterior teeth lack carinae and are constricted basally. In some specimens, the striae do not reach the base of the crown and are instead restricted to the apical area with the basal half of the enamel surface being smooth, while in others, the striae reach the base in the middle of the lingual surface and are absent mesially and distally. Although both the labial and lingual surfaces of the enamel are crenulated, several teeth show denser and more prominent striae on the lingual side compared to the labial (e.g. MTM PAL 2014.124.7, 2014.124.11). The apices are located lingual to the medial plane of the tooth and are usually confluent with the rest of the tooth crown, giving it a rounded shape, although in some specimens, the apex is distinct, being bordered by a mesial and distal depression similarly to the posterior teeth of ‘*Aprosuchus ghirai’*. The 12^th^ dentary tooth in MTM VER 2026.27 and a posterior tooth associated with MTM V 128_447 belong to this morphotype. Large, oval wear facets are commonly preserved on the occlusal surface of the posterior teeth. This tooth morphotype is unique among atoposaurids and instead resembles the posterior teeth of derived neosuchian taxa such as *Bernissartia fagesi* and *Allognathosuchus haupti* (Ősi, 2014). Isolated teeth of the above-described morphology are assigned to *S. tribodon* from the coeval Iharkút and Ajka sites (Ősi et al., 2012, 2016). Morphologically similar teeth are known from several Late Cretaceous sites in Central Europe including the Turonian (Gams, Ősi et al., 2019) and Coniacian (Tiefengraben, Ősi et al., 2021) of Austria and the Maastrichtian of Romania (Haţeg Basin, e.g. Botfalvai et al., 2021).

###### Maxillary teeth

The maxillary dentition is incompletely preserved in both specimens. The 3^rd^ (MTM V 2010.235.1., possibly a replacement tooth) and 7^th^ to 9^th^ (MTM V 2010.236.1) teeth are completely preserved in situ, the 2^nd^ (MTM V 2010.236.1.), 4^th^ (both specimens) and 8^th^ (MTM V 2010.235.1.) are partially preserved, broken at the base and missing most or all of the crown, while the rest of the teeth are absent in both specimens. Apical wear facets are visible in all preserved posterior teeth of the posterior morphotype, while the anterior 3^rd^ tooth in MTM V 2010.235.1. appears completely unworn, likely due to its possible status as a replacement tooth. The reduction in tooth size posterior to the 4-5^th^ alveolus is shared with *Sabresuchus sympiestodon* but not ‘*Aprosuchus ghirai’* which may be a result of the latter’s juvenile status. Similarly to *Sabresuchus sympiestodon* but unlike *Theriosuchus pusillus, Theriosuchus grandinaris, Knoetschkesuchus* spp. and *Sabresuchus ibericus*, only one enlarged caniniform is present in each maxilla.

The posterior teeth of both ‘*Aprosuchus ghirai’* and *Sabresuchus sympiestodon* are significantly more compressed labiolingually, taper more strongly mesially and distally, and feature more distinct carinae than those of *Sabresuchus tribodon*. In both Romanian taxa, the apicobasal crenulations on the tooth crown reach the carinae, resulting in strongly denticulate carinae on unworn teeth superficially resembling a true ziphodont condition. The denticles in *A. ghirai* are proportionately larger compared to the size of the crown than those of *S. sympiestodon*. Furthermore, in ‘*A. ghirai’*, the pointed apices of the teeth are more distinct, bordered by concavities mesially and distally.

###### Dentary teeth

The fragmentary specimen VER 2016.92. preserves a single tooth of the first morphotype from the symphyseal region, with a strongly labially deflected apicobasal axis, possibly due to the broken lateral margin of the bone. The tooth crown is lingually bent, bearing apicobasal striae and a small apical wear facet. MTM VER 2026.28 preserves an in-situ 4^th^ caniniform tooth of the anterior morphotype, and an associated posterior tooth of the posterior morphotype, as well as fragmentary remains of the 3^rd^ and 5^th^ teeth. The 4^th^ tooth bears a tall, narrow crown with a labially slightly deflected axis, and the alveolus containing it is significantly larger than those anterior or posterior to it. The crown is bent lingually, with a constricted base, and a convex labial and concave lingual surface. It bears distinct, unserrated carinae and apicobasal striations. The posterior tooth is mesiodistally wider than tall with a worn apex, and associated fragments of a massive root.

MTM VER 2026.27 preserves two in situ teeth. The 10^th^ tooth is lanceolate, labiolingually flattened and mesiodistally symmetrical, belonging to the anterior morphotype. The mesial and distal carinae are well developed and lack denticles. The main body of the tooth bears apicobasal striae. The tooth projects outwards, with a c. 140° angle between the axis and the alveolar plane, which may be due in part to preservation. The base of the tooth is constricted. The 12^th^ tooth is mesiodistally expanded, belonging to the posterior morphotype. The apex of the tooth is heavily worn. The tooth lacks carinae. The base of the tooth is constricted.

While the 7^th^ right dentary tooth in the holotype of *A. ghirai*, as well as some teeth tentatively assigned to *S. sympiestodon* or cf. *Theriosuchus* sp. from the Haţeg Basin preserve denticles on the carina that are not the result of apicobasal striae, making them true ziphodont teeth (Venczel & Codrea, 2019), this condition is not known in any of the Iharkút atoposaurid teeth.

Sabresuchus ibericus Brinkmann, 1989

###### Holotype

IPFUB 102/21.2, right maxilla.

###### Type locality

Lowest horizon of an abandoned lignite mine, La Huerguina Formation (Uña), Upper Barremian (Early Cretaceous); south-west of the eastern edge of Uña, Serrania de Cuenca, eastern Spain (Brinkmann, 1989; Tennant et al., 2016).

###### Referred material

As in Brinkmann (1989).

###### Revised diagnosis

Characters defined as diagnostic by Tennant et al. (2016): grooved ornamentation of maxillary palate; depressed dorsal surface of the parietal compared to the squamosal (unknown in *S. tribodon*); proportionally short dentary symphysis ending posteriorly at the level of the 5^th^ alveolus; all dentary teeth occupy a longitudinal dental groove instead of the anterior teeth occupying separate alveoli; occlusal dentary surface lacks nutrient foramina; hypertrophied caniniform maxillary tooth directed posteroventrally instead of ventrally; fully ziphodont tooth crowns with serrated carinae. *S. ibericus* can be further differentiated from both *S. sympiestodon* and *S. tribodon* by lacking a diastema between the 7^th^ and 8^th^ dentary tooth and lacking crenulations on the lingual and labial surfaces of the teeth.

###### Remarks

Tennant et al. (2016) considered the grooved, rugose decoration of the outer surface of the posterior dentary (S6) diagnostic for this species. However, this condition is also present in *S. tribodon* and ‘*Aprosuchus ghirai’* (*Sabresuchus sympiestodon*).

##### Species **Sabresuchus sympiestodon** Martin, Rabi and Csiki, 2010

2010 *Sabresuchus sympiestodon* Martin, Rabi and Csiki

2019 *Aprosuchus ghirai;* Venczel and Codrea

###### Type material

LPB (FGGUB) R.1782, right maxilla (holotype); LPB (FGGUB) R.1781, partial skull roof (paratype).

###### Type locality

Oltoane Hill, Tuştea (N45°36′25″ E22°50′53″), Haţeg Basin, Transylvania, Romania. Upper part of unnamed Middle Member, Densuş-Ciula Formation (Grigorescu, 1992; Albert et al., 2025), late Maastrichtian.

###### Referred material and range

Material referred to *S. sympiestodon* in Martin et al. (2014): MCDRD 134, dentary, Cioaca Târnovului, Sânpetru section, Sânpetru Formation, early Maastrichtian. MCDRD 793, adult maxilla, lower part of the Sânpetru section, Sânpetru Formation, early Maastrichtian. LPB (FGGUB) R.1945, fragmentary juvenile maxilla, Fântânele microvertebrate bonebed, lower Densus-Ciula Formation, early Maastrichtian. Material previously considered referable to *Aprosuchus ghirai*: UBB V.562/1, partial skull; UBB V.562/2, vertebra from the Pui Gater locality, near Pui village, Haţeg Basin, Transylvania, Romania. ‘Pui beds’, correlated with lower part of Sânpetru Formation, early Maastrichtian (Venczel & Codrea, 2019; Albert et al., 2025).

###### Revised diagnosis

A species of *Sabresuchus* by possessing traits diagnostic for the genus (see above). *Sabresuchus sympiestodon* is unique among Atoposauridae in that the longitudinal ridge of the frontal is anteroposteriorly restricted in extent (Martin et al., 2010, 2014; Tennant et al., 2016). *S. sympiestodon* differs from *Sabresuchus tribodon* in possessing an orbitonasal sulcus along the maxilla similar to the one present in *Theriosuchus pusillus*. Furthermore, *S. sympiestodon* is distinguished from *Sabresuchus tribodon* and *S. ibericus* by the following combination of traits: single lateral maxillary plane, labiolingually strongly flattened posterior teeth and the dentary internal alveolar margins being flat and confluent with the rest of the dentary occlusal surface. See *S. tribodon* and *S. ibericus* for further differences.

###### Remarks

Both Martin et al. (2010, 2014) and Tennant et al. (2016) considered the single, ventrally directed hypertrophied fifth maxillary tooth, and the lack of striae on the labial and lingual surfaces of the anterior maxillary teeth as diagnostic for *S. sympiestodon* along with the diastema between the 7^th^ and 8^th^ dentary alveoli. However, as *Sabresuchus tribodon* also shares these characteristics (see Results: Phylogenetic analyses), they are no longer considered diagnostic for *S. sympiestodon* and are instead synapomorphies of the two Eastern European *Sabresuchus* species.

‘Aprosuchus ghirai’ Venczel and Codrea, 2019

nomen invalidum (junior synonym of *Sabresuchus sympiestodon* [Martin, Rabi & Csiki, 2010])

###### Type material

UBB V.562/1, partial skull.

###### Type locality

Pui Gater locality, near Pui village, Haţeg Basin, Transylvania, Romania. Lower ‘Pui beds’, early Maastrichtian.

###### Referred material and range

None.

###### Comments

Fragmentary atoposaurid material have been reported from several Campanian-Maastrichtian continental vertebrate sites in the Haţeg Basin of Romania (Botfalvai et al., 2021, 2025; Csiki-Sava et al., 2010, 2016; Vasile & Csiki-Sava, 2010; Vasile et al., 2019; Venczel & Codrea, 2019). Most of this material represents isolated teeth with the notable exception of *Sabresuchus sympiestodon* represented by a partial skull and other cranial remains from the lower Maastrichtian of Tuştea and Sânpetru (Martin et al., 2010, 2014), as well as ‘*Aprosuchus ghirai’* Venczel and Codrea, 2019 known from a nearly complete juvenile skull from the stratigraphically slightly lower ‘Pui Swamp’ site (Albert et al., 2025; Botfalvai et al., 2025). A further isolated fragmentary juvenile maxilla attributed to *S. sympiestodon* is known from the Fântânele microvertebrate site roughly coeval with the Pui beds (Botfalvai et al., 2025; Martin et al., 2014). Venczel and Codrea (2019) differentiated ‘*A. ghirai’* from *S. sympiestodon* by a W-shaped nasofrontal suture (straight in *S. sympiestodon*), presence of an antorbital fenestra, a dorsolaterally closed cranioquadrate passage, and the lack of a hypertrophied caniniform tooth. However, while incompletely preserved, an anterorbital fenestra was likely present in *Sabresuchus sympiestodon*. An elongated, anteriorly clearly demarcated depression along the posterior process of the maxilla appears to correspond to the rim of this opening as seen in *Sabresuchus ibericus* with an apparent antorbital fenestra (Brinkman, 1992). The W-shaped nasofrontal suture and lack of a hypertrophied 4^th^ maxillary tooth in ‘*A. ghirai’*, on the other hand, are indicative of the juvenile ontogenetic stage of the holotype (Erickson et al., 2003; Gignac & Erickson, 2015, 2016; Iordansky, 1973) as otherwise suggested by Venczel and Codrea (2019) based on the incomplete fusion of the frontals and parietals, proportionately large orbits, and a short rostrum. Although Venczel and Codrea (2019) proposed that the two roughly coeval species could have filled different ecological niches based on their distinct sizes, ‘*A. ghirai’* falls within the size range of *S. sympiestodon* (Martin et al., 2014) and the two cannot be distinguished on the basis of tooth morphology which is known to correlate with feeding ecology in crocodyliforms (Ősi, 2014; Vasile et al., 2019). Our first-hand examination of both taxa reveals no further apparent morphological differences, and they are here recovered as close relatives (Fig. 1A-B), in contrast to previous work (Allen et al. 2025; Pochat-Cottilloux et al., 2024; Venczel & Codrea, 2019). We therefore consider ‘*Aprosuchus ghirai’ a junior* synonym of the stratigraphically and geographically proximate *Sabresuchus sympiestodon*.

> *Ogresuchus furatus* Sellés et al. 2020
>
> nomen dubium

###### Type material

MCD-7149, partial semi-articulate skeleton preserving parts of the rostrum and associated axial and appendicular elements.

##### Type locality

El Mirador site, Coll de Nargó area, Lleida Province, Catalonia, Spain. ’Lower gray unit’ of Tremp Formation, Upper Cretaceous, lower Maastrichtian. (Sellés et al., 2020)

###### Referred material and range

As in type material.

###### Comments

This taxon was interpreted as a European representative of notosuchian crocodyliforms when described by Sellés et al. (2020), but more recent work reinterpreted it as an atoposaurid (Szegszárdi et al., 2026). The phylogeny of the present study confirms this placement and recovers this taxon in a clade with *Alligatorium* sp. and *Theriosuchu*s spp. Furthermore, the fragmentary nature of the available cranial material renders this taxon undiagnostic in the context of Atoposauridae. Minimal anatomical overlap with stratigraphically congruent *Sabresuchus* species consequently precludes unequivocal taxonomic differentiation. *Ogresuchus furatus* is therefore considered a nomen dubium and is referred to as an indeterminate atoposaurid.

## Discussion

### Taxonomy and durophagous adaptations of *Sabresuchus tribodon*

As each of our analyses using three different datasets find *Theriosuchus pusillus* (type species of the genus) to be more closely related to *Alligatorium* sp. than to *tribodon* sp. nov. or *Sabresuchus* (*Theriosuchus*) *sympiestodon* (Fig. 1), we tentatively assign the new species to the genus *Sabresuchus*. *Sabresuchus tribodon* (Fig. 7) is distinct from *Sabresuchus sympiestodon*, ‘*Aprosuchus ghirai’* (a taxon here recognized as a junior synonym of *S. sympiestodon*; see Systematic Palaeontology), and all other atoposaurids in the presence of massive, bulbous posterior teeth with apicobasal striae, warranting its recognition as a separate taxon. Comparable rear dentition is known in other Cretaceous crocodyliforms from Europe including the bernissartiid *Bernissartia fagesii*, *Koumpiodontosuchus aprosdokiti*, and the hylaeochampsid *Acynodon adriaticus* (Martin et al., 2020; Muscioni et al., 2024; Ősi, 2014; Sweetman et al., 2014). This specialized morphology is commonly inferred to represent adaptation to durophagy (e.g. Brochu, 2004; Ősi, 2014; Young et al., 2014), which in turn may not necessarily predict a specialized niche (Darlim et al., 2023). Capability of crushing hard food items is furthermore supported by the relatively short rostrum of *S. tribodon*, even compared to other brevirostrine forms (e.g. Allen et al., 2025). Evidence for a brevirostrine condition includes the proportions of the maxilla and dentary, the reduced number of teeth posterior to the maxillary notch compared to *S. sympiestodon*, and the posteriorly shifted suborbital fenestra compared to the tooth row. Furthermore, the marked dorsal expansion of the posterior dentary and surangular provides increased attachment for the external adductor muscles (M. adductor mandibulae externus medialis and superficialis) which is common in durophagous taxa (Ősi, 2014). A posteroventrally turned retroarticular process with a large lateral surface could in turn provide an attachment surface for a prominent M. pterygoideus ventralis (Holliday & Witmer, 2007; Ősi, 2014). This is in contrast with the condition in *S. sympiestodon* (as preserved in ‘*A. ghirai’*) lacking specialized bulbous rear teeth and possessing a dorsoventrally shallower posterior dentary and surangular. As such, the high degree of morphological similarity between *S. tribodon* and crocodyliforms such as *Thikarisuchus xenodentes* (Allen et al., 2025), is based on characters highly subject to ecomorphological convergence rather than true shared ancestry (see Systematic Palaeontology). Although some studies infer a terrestrial habitat for atoposaurids and related brevirostrine forms based on cranioanatomical features (e.g. altirostral skull proportions, enlarged palpebrals, forward-facing external nares), depositional environment, and inferred diet based on tooth morphology (Allen et al., 2025; Brinkmann, 1992), others instead propose semiaquatic affinities for the group based on limb bone morphology (Eijkelboom, 2020) and osteoderm ornamentation (Pochat-Cottilloux et al., 2024; contra Allen et al., 2025). Notably, a durophagous feeding ecology often coincides with a semiaquatic or aquatic lifestyle in crocodyliforms (Darlim et al., 2023; Muscioni et al. 2024; Sweetman et al., 2014) and testudines (Evers & Joyce, 2020; Hermanson & Evers, 2026; Parham & Pyenson, 2010). At present, the preferred lifestyle of *Sabresuchus tribodon* cannot be accurately determined until more complete material is recovered, or geochemical analyses are performed.

**Figure 7.** Cranial reconstruction of *Sabresuchus tribodon* sp. nov. from the Santonian of Hungary with the preserved elements highlighted. Illustrations by Márton Zsoldos. Scale bar = 10 mm.

### Systematics of European Cretaceous atoposaurids

Historically, European Cretaceous and Jurassic atoposaurids have been widely referred to the genus *Theriosuchus* Owen, 1878 (Allain et al., 2022; Blanco et al., 2020; Brinkmann, 1992; Buffetaut, 1983; Buscalioni & Sanz, 1988; Karl et al., 2006; Martin et al. 2010, 2014, 2020; Ősi et al., 2019; Puértolas-Pascual et al., 2015; Schwarz & Salisbury, 2005; Schwarz et al., 2017; Schwarz-Wings et al., 2009; Tennant et al., 2016; Venczel & Codrea, 2019; Young et al., 2016). Among these records, the type species, *Theriosuchus pusillus* Owen, 1878 from the Berriasian Purbeck Limestone Group, represents the most exceptionally preserved material. In contrast, other stratigraphically older records remain problematic. Beyond Europe, most but not all phylogenetic analyses support a close systematic relationship between *Theriosuchus grandinaris* from the Early Cretaceous of Thailand and the type species *T. pusillus* (Allen et al., 2025; Lauprasert et al., 2011; Schwarz et al., 2017; Turner, 2015; present study).

The stratigraphically youngest *Theriosuchus* (*Sabresuchus*) *sympiestodon* from the latest Cretaceous of Romania was originally interpreted as a late surviving representative of *Theriosuchus* (Martin et al., 2010; 2014), consistent with other relict elements of the European Late Cretaceous tetrapod fauna (Csiki et al., 2015; Szegszárdi et al., 2026). With a record spanning the Middle Jurassic to the Late Cretaceous, *Theriosuchus* may therefore constitute a particularly persistent evolutionary lineage with the herein described new taxon, recovered as the sister-taxon of *sympiestodon*, partially infilling the sampling gap between the Lower and uppermost Cretaceous (Martin et al., 2010; Young et al., 2016; Fig. 8). However, subsequent comprehensive phylogenetic analyses have challenged the monophyly of the genus due to the exclusion of *Theriosuchus sympiestodon*, *Theriosuchus ibericus, Theriosuchus guimarotae* and *Knoetschkesuchus langenbergensis* from the core clade of *Theriosuchus* (Schwarz et al., 2017; Tennant et al., 2016; Turner, 2015). In particular, the Late Cretaceous *T. sympiestodon* and the Early Cretaceous *T. ibericus* were resolved closer to Paralligatoridae than to *T. pusillus*, prompting their transfer to the newly erected genus *Sabresuchus* (Tennant et al., 2016). In parallel, *T. guimarotae* was reassigned to the genus *Knoetschkesuchus* along with *K. langenbergensis* (Schwarz et al., 2017). Recently, a partial neurocranium from the Late Jurassic of Portugal, assigned to Atoposauridae indet. by Puértolas-Pascual (2025), was found by the author’s phylogenetic analysis to be the sister taxon of *sympiestodon*; however, as the specimen is fragmentary and linked to *sympiestodon* by unreliable synapomorphies (e.g. ‘the bar between the orbit and the supratemporal fossa is narrow, with sculpting restricted to its anterior surface in mature specimens’, despite the skull roof only being known in juvenile specimens of *sympiestodon*), further study is needed to determine their relationship, especially considering the nearly 85 million year gap between the two occurrences (Albert et al., 2025; Puértolas-Pascual, 2025).

**Figure 8.** Spatial and temporal distribution of species assigned to Atoposauridae. **A**. Late Cretaceous paleobiogeographic map showing the distribution of atoposaurid species: 1. *Alligatorellus bavaricus* and *Atoposaurus oberndorferi*, Solnhofen region, Germany; 2. *Alligatorellus beaumonti*, *Alligatorium meyeri* and *Atoposaurus jourdani*, Cerin, France; 3. *Knoetschkesuchus guimarotae*, Guimarota, Portugal; 4. *Knoetschkesuchus langenbergensis*, Langenberg, Germany; 5. *Sabresuchus ibericus*, Uña, Spain; 6. *Sabresuchus sympiestodon* and ‘*Aprosuchus ghirai’*, Haţeg basin, Romania; 7. *Sabresuchus tribodon* sp. nov., Iharkút, Hungary. 8. *Theriosuchus grandinaris* and *Varanosuchus sakonnakhonensis*, Sakon Nakhon, Thailand; 9. *Theriosuchus pusillus*, Dorset, UK. **B**. stratigraphic distribution of putative atoposaurid species. Map modified from Wikimedia Commons (https://commons.wikimedia.org/wiki/File:Afro-Eurasia_location_map_with_borders.svg), timescale modified from the International Chronostratigraphic Chart v2024-12 (Cohen et al., 2025), geographic distribution and temporal range data from Tennant et al. (2016), Pochat-Cottilloux et al. (2024) and the Paleobiogeography Database (Uhen et al., 2023).

The more inclusive *Theriosuchus* concept of Martin et al. (2010, 2014) united the genus by the presence of an orbitonasal groove of the snout in *T*. *(Knoetschkesuchus) guimarotae*, *Theriosuchus pusillus*, and *Theriosuchus* (*Sabresuchus) sympiestodon* (preserved in the maxilla only). Nevertheless, this character is not included in their phylogenetic analyses or that of Tennant et al. (2016). In our analyses of the datasets adapted from Szegszárdi et al. (2026) and Pochat-Cottilloux et al. (2024), this character (311) is uniting an even more inclusive clade since an orbitonasal sulcus is widespread in paralligatorids (Turner, 2015) as well as in other basal neosuchians (Allen et al., 2025; Sereno & Larsson, 2009; Wu et al., 2001). Contrary to Tennant et al. (2016), we confirm the presence of the sulcus in both *S. sympiestodon* and *T. pusillus*, whereas it is not apparent in the maxilla of *S. tribodon* or *S. ibericus*. Martin et al. (2010) furthermore considered the low-crowned, labiolingually compressed, basally constricted, pseudoziphodont posterior teeth synapomorphic of *Theriosuchus* (containing *Theriosuchus pusillus*, *Theriosuchus grandinaris*, *Knoetschkesuchus guimarotae*, *Sabresuchus ibericus* and *Sabresuchus sympiestodon*), but subsequent work demonstrated a widespread presence of this trait among short-snouted basal neosuchians (Adams, 2014; Allen et al., 2025). Nevertheless, Tennant et al. (2016) pointed out that species of *Sabresuchus* share a posterior dental morphotype with *T. pusillus* by exhibiting mesiodistally broader than apicobasally tall crowns with apical margins orientated at <45° from the horizontal. In addition, *S. tribodon* preserves a sculptured splenial (unknown in *S. sympiestodon*), which is notably shared with *T. pusillus*, *T. grandinaris*, *T. (Knoetschkesuchus) guimarotae*, but also with species of *Shamosuchus* (Tennant et al., 2016).

Each of the three analyses of the present study infers close relationship between *Sabresuchus tribodon* sp. nov. and *Sabresuchus sympiestodon*, as well as a polyphyletic *Theriosuchus* (sensu Martin et al., 2010, 2014; Fig. 1). In our analyses of the two datasets derived from Turner (2015; Fig. 1A-B), the type species of *Theriosuchus*, *T. pusillus*, is placed more closely to *Alligatorium* spp. than to *Sabresuchus* spp. due to an anterior choanal edge situated anteriorly between the suborbital fenestra (Ch. 44, state 0; Szegszárdi et al. [2026], Pochat-Cottilloux et al. [2024]), differing from the condition in *S. sympiestodon* (‘*Aprosuchus ghirai*)’, where it is located near the posterior edge of the fenestra (state 1; unknown in *S. tribodon*). Although several synapomorphies uniting *Sabresuchus* spp. are interpreted as absent in *Theriosuchus pusillus* (see Results; Phylogenetic analysis), a targeted reevaluation of these specific characters across both taxa would facilitate a more robust assessment of their phylogenetic relationships. Using the dataset adapted from Tennant et al. (2016; Fig. 1C), *Sabresuchus* spp. is grouped with the paralligatorids *Shamosuchus djadochtaensis*, *Wannchampsus kirpachi* and *Brillanceausuchus babouriensis* rather than *Theriosuchus* spp. by the straight lateral contour of the snout (ch. 6, state 0), a dorsal anteroposterior ridge running along the entire length of the frontal (ch. 101, state 3; restricted to the posterior section of the frontal in *S. sympiestodon*), and the absence of an obliquely-oriented ridge on the dorsal surface of the squamosal (ch. 145, state 0). However, since Tennant et al. (2016) is restricted to atoposaurid and paralligatorid ingroup taxa, a more global taxonomic sampling may alter the inclusiveness of *Theriosuchus* within this dataset. In fact, the highly restricted concept of Atoposauridae in Tennant et al. (2016), as applied to their topologies, is not reproducible from their own phylogenetic definition, which includes the crown-group as an external specifier, yet their analyses/topologies are without any crown-group taxa.

A comprehensive evaluation of atoposaurid, paralligatorid, and *Theriosuchus* inter- and ingroup relationships is currently precluded by limited character and taxon sampling across individual datasets. Indeed, recent work demonstrated that North American paralligatorids are morphologically highly reminiscent of European and Asian taxa presently recovered as atoposaurids (Adams, 2014, 2019; Allen et al., 2025; Noto et al., 2020).

Reconciling these phylogenetic matrices (e.g., Allen et al., 2025; Szegszárdi et al., 2026; Tennant et al., 2016) is critical for more confident systematic and nomenclatural conclusions concerning the status of *Theriosuchus* and *Sabresuchus*. In addition, the relationships of Cretaceous South-East Asian taxa to atoposaurids and paralligatorids warrants re-evaluation since current phylogenies imply complex biogeographic histories and extensive ghost lineages (Lauprasert et al., 2011; Pochat-Cottilloux et al., 2024).

## Conclusions

1. *Sabresuchus tribodon* is a new species of neosuchian crocodyliform from the Santonian of Hungary. It is characterized by bulbous, mesiodistally wider than tall posterior teeth with enamel surface ornamented with apicobasal ridges; an extremely short rostrum with a reduced number of teeth, and an upturned dorsal margin of the dorsally highly expanded dentary and surangular. While comparable adaptations are present among durophagous taxa, this combination of characters is not known in any atoposaurid or paralligatorid taxa.
2. Parsimony analyses of three separate datasets consistently retrieve S. *sympiestodon* from the Maastrichtian of Romania as the sister taxon of *S. tribodon*. Furthermore, we interpret ‘*Aprosuchus ghirai’* from the Maastrichtian of Romania as a junior synonym of *Sabresuchus sympiestodon*, and *Ogresuchus furatus* from the Maastrichtian of Spain as an indeterminate atoposaurid.
3. Our systematic reassessment tentatively confirms the polyphyly of *Theriosuchus* (sensu Martin et al., 2010), although future research on more complete material and a combination of differently focused datasets is needed to reach a more definitive conclusion.
4. Further evaluation of the in- and outgroup relationships of *Theriosuchus*, Atoposauridae and Paralligatoridae is crucial to clarify the taxonomy and phylogeny of *Theriosuchus* and *Sabresuchus*. Furthermore, a re-evaluation of phylogenetic relationships between putative paralligatorid and atoposaurid taxa from Eurasia and North America is necessary for reconstructing the groups’ evolutionary history in a paleobiogeographic context.

## Acknowledgements

We thank János Magyar and Márton Szabó (Hungarian Natural History Museum), Liana Săsăran and Lóránd Silye (Babeș-Bolyai University), Dorin Cărăbeț and Dorina Dan (Museum of Dacian and Romanian Civilization), Zoltán Csiki-Sava (University of Bucharest), Àngel Galobart, Irina Fernández, Laura Fàbrega, and Óscar Castillo-Visa (Institut Català de Paleontologia Miquel Crusafont) for their help in accessing and examining specimens in their care. We are thankful to Gábor Botfalvai and Gergő Konecsni (ELTE Eötvös Loránd University), Albert G. Sellés (Institut Català de Paleontologia Miquel Crusafont), Paul Rummy (Chinese Academy of Sciences), and Yohan Pochat-Cottilloux (University of Warsaw) for sharing their research, and to the Willi Hennig Society for providing free access to TNT. We thank Nóra Sebők for her photographs and research of the specimens described above, and Márton Zsoldos for illustrating the cranial reconstruction. We are thankful to the participants and sponsors of the Iharkút digs; the fieldwork and research of the Iharkút site was supported by the Hungarian Academy of Sciences, the Hungarian Natural History Museum, the National Geographic Society, the Hungarian Scientific Research Fund, the Eötvös Loránd University Departments of Paleontology and Applied and Physical Geology, the Jurassic Foundation, the Hantken Miksa Foundation, the Hungarian Dinosaur Foundation, the Pro Renovanda Cultura Hungariae Foundation, Nemzeti Kutatási és Technológiai Hivatal, and the Hungarian Oil and Gas Company (MOL).

## Author contributions

**Conceptualization** M.R., A.Ő.; **Data Curation** A.Ő.; M.Sz.; **Formal Analysis** M.Sz.; M.R.; **Funding Acquisition** A.Ő., M.Sz.; **Investigation** M.Sz., M.R., A.Ő.; **Methodology** M.R.; **Project Administration** A.Ő.; **Resources** A.Ő.; **Supervision** M.R., A.Ő.; **Validation** M.R.; **Visualization** M.Sz., M.R.; **Writing–Original Draft Preparation** M.Sz., M.R.; **Writing–Review & Editing** M.R., M.Sz., A.Ő.

## Data availability

This published work and the nomenclatural acts it contains, have been registered in ZooBank: http://zoobank.org/References/XXXXXXXXX (to be added upon acceptance)

## Data deposition

Data for this study [including supplementary datasets S1, S2 and S3, and supplementary document S4] are available in the MorphoBank Digital Repository: (to be added upon acceptance).

## Declaration of generative AI use

The authors report generative AI was not used in their research or preparation of this manuscript.

