## Supplementary document S4 for "Durophagous adaptations in a new crocodyliform from the Santonian of Hungary and a systematic reassessment of European Cretaceous atoposaurids"

This document contains character and state descriptions for the phylogenetic matrices used in this study. The matrices were adapted without adding or modifying any characters.

**1. Character and state descriptions for the matrix adapted from Szegszárdi et al. (2026) (Supplementary Dataset S1)**

**Character 1** (Rummy et al., 2022: char. 1, modified from Clark, 1994: char. 1): External surface of dorsal cranial bones: smooth (0), slightly grooved (1), or heavily ornamented with deep pits and grooves (2).

**Character 2** (Rummy et al., 2022: char. 2, modified from Clark, 1994: char. 2): Skull expansion at orbits: gradual (0), or abrupt (1).

**Character 3** (Rummy et al., 2022: char. 3, modified from Clark, 1994: char. 3): Rostrum proportions: narrow oreinirostral (0), broad oreinirostral (1), nearly tubular (2), or platyrostral (3).

**Character 4** (Clark, 1994: char. 4): Premaxilla participation in internarial bar: forming at least the ventral half (0), or with little participation (1).

**Character 5** (Clark, 1994: char. 5): Premaxilla anterior to nares: narrow (0), or broad (1)

**Character 6** (Rummy et al., 2022: char. 6, modified from Clark, 1994: char. 6): + External nares facing anterolaterally or anteriorly (0), dorsally not separated by premaxillary bar from anterior edge of rostrum (1), or dorsally separated by premaxillary bar (2).

**Character 7** (Clark, 1994: char. 7): Palatal parts of premaxillae: do not meet posterior to incisive foramen (0), or meet posteriorly along contact with maxillae (1).

**Character 8** (Clark, 1994: char. 8): Premaxilla-maxilla contact: premaxilla loosely overlies maxilla (i.e. posterodorsal process of the premaxilla overlaps anterodorsal surface of the maxilla) (0), or sutured together along a butt joint (1).

**Character 9** (Rummy et al., 2022: char. 9, modified from Clark, 1994: char. 9): Ventrally opened notch on ventral edge of rostrum at premaxilla-maxilla contact: absent (0), present as a notch (1), or present as a large fenestra (2).

**Character 10** (Rummy et al., 2022: char. 10, modified from Clark, 1994: char. 10): + Posterior palatal branches of maxillae anterior to palatines: do not meet (0), or meet extensively but posterior most parts fail to meet (1), or meet entirely (2).

**Character 11** (Clark, 1994: char. 11): Nasal contacts lacrimal (0), or does not contact (1).

**Character 12** (Clark, 1994: char. 12): Lacrimal contacts nasal along medial edge only (0), or medial and anterior edges (1).

**Character 13** (Clark, 1994: char. 13): Nasal contribution to narial border: yes (0), or no (1).

**Character 14** (Clark, 1994: char. 14): Nasal-premaxilla contact: present (0), or absent (1).

**Character 15** (modified from Clark, 1994: char. 15): Descending process of prefrontal: does not contact palate (0), or contacts palate (1).

**Character 16** (Rummy et al., 2022: char. 16): Postorbital-jugal contact: postorbital anterior to jugal (0), or postorbital medial to jugal (1), or postorbital lateral to jugal (2).

**Character 17** (Clark, 1994: char. 17): Anterior part of the jugal with respect to posterior part: as broad (0), or twice as broad (1).

**Character 18** (Clark, 1994: char. 18): Jugal bar beneath infratemporal fenestra: flattened (0), or rod shaped (1).

**Character 19** (Clark, 1994: char. 19): Quadratojugal dorsal process: narrow, contacting only a small part of postorbital (0), or broad, extensively contacting the postorbital (1).

**Character 20** (Clark, 1994: char. 20): Frontal width between orbits: narrow, as broad as nasals (0), or broad, twice as broad as nasals (1).

**Character 21** (Clark, 1994: char. 21): Frontals: paired (0), or unpaired (1).

**Character 22** (Clark, 1994: char. 22): Dorsal surface of frontal and parietal: flat (0), or with midline ridge (1).

**Character 23** (modified from Clark, 1994: char. 23, by Buckley and Brochu, 1999: char. 81): + Parieto- postorbital suture: absent from dorsal surface of skull roof and supratemporal fossa (0), absent from dorsal surface of skull roof but broadly present within supratemporal fossa (1), or present within supratemporal fossa and on dorsal surface of skull roof (2).

**Character 24** (Clark, 1994: char. 24): Supratemporal roof dorsal surface: complex (0), or dorsally flat ‘‘skull table’’ developed, with postorbital and squamosal with flat shelves extending laterally beyond quadrate contact (1).

**Character 25** (modified from Clark, 1994: char. 25) Postorbital bar: sculpted (if skull sculpted) (0), or unsculpted (1).

**Character 26** (modified from Clark, 1994: char. 26): Postorbital bar: transversely flattened (0), or cylindrical (1).

**Character 27** (Clark, 1994: char. 27): Vascular opening in dorsal surface of postorbital bar: absent (0), or present (1).

**Character 28** (modified from Clark, 1994: char. 28): Postorbital anterolateral process: absent or poorly developed (0), or well developed, long, and acute (1).

**Character 29** (Clark, 1994: char. 29): Dorsal part of the postorbital: with anterior and lateral edges only (0), or with anterolaterally facing edge (1).

**Character 30** (Clark, 1994: char. 30): Dorsal end of the postorbital bar broadens dorsally, continuous with dorsal part of postorbital (0), or dorsal part of the postorbital bar constricted, distinct from the dorsal part of the postorbital (1).

**Character 31** (Clark, 1994: char. 31): Bar between orbit and supratemporal fossa broad and solid, with broadly sculpted dorsal surface if sculpture present (0), or bar narrow, sculpting restricted to anterior surface (1).

**Character 32** (modified from Clark, 1994: char. 32): Parietal: with broad occipital portion (0), or without broad occipital portion (1).

**Character 33** (Clark, 1994: char. 33) Parietal: with broad sculpted region separating fossae (0), or with sagittal crest between supratemporal fossae (1).

**Character 34** (Clark, 1994: char. 34): Postparietal (dermosupraoccipital): a distinct element (0), or not distinct (fused with parietal?) (1).

**Character 35** (Clark, 1994: char. 35): Posterodorsal corner of the squamosal: squared off, lacking extra ‘‘lobe’’ (0), or with unsculptured ‘‘lobe’’ (1).

**Character 36** (Rummy et al., 2022: char. 36, modified from Clark, 1994: char. 36): Posterolateral process of squamosal: poorly developed and projected horizontally at the same level of the skull (0), elongated, thin, and posteriorly directed, not ventrally deflected (1), or elongated, posterolaterally directed, and ventrally deflected (2).

**Character 37** (Clark, 1994: char. 37): + Palatines: do not meet on palate below the narial passage (0), form palatal shelves that do not meet (1), or meet ventrally to the narial passage, forming part of secondary palate (2).

**Character 38** (Clark, 1994: char. 38): Pterygoid: restricted to palate and suspensorium, joints with quadrate and basisphenoid overlapping (0), or pterygoid extends dorsally to contact laterosphenoid and form ventrolateral edge of the trigeminal foramen, strongly sutured to quadrate and latero- sphenoid (1).

**Character 39** (Rummy et al., 2022: char. 39, modified from Clark, 1994: char. 39): Choanal opening: continuous with pterygoid ventral surface except for anterior and anterolateral borders (0), or opens into palate through a deep midline depression (choanal groove) (1).

**Character 40** (Clark, 1994: char. 40): Palatal surface of pterygoids: smooth (0), or sculpted (1).

**Character 41** (Clark, 1994: char. 41): Pterygoids posterior to choanae: separated (0), or fused (1).

**Character 42** (modified from Clark, 1994: char. 42 by Ortega et al., 2000: char. 139): Depression on primary pterygoidean palate posterior to choana: absent or moderate in size, narrower than palatine bar (0), or wider than palatine bar (1).

**Character 43** (Rummy et al., 2022: char 43, modified from Clark, 1994: char. 43): + Primary pterygoidean palate: forms posterior half of the choanal opening (0), or forms posterior, lateral, and part of the anterior margin of the choana (1), or completely enclose choana (2).

**Character 44** (Rummy et al., 2022: char 44, modified from Pol and Norell, 2004a; Clark, 1994: char. 44): + Anterior edge of choanae situated between the suborbital fenestra (or anteriorly) (0), situated near the posterior edge of suborbital fenestra (1), or posterior to the suborbital fenestra (reaching in some cases the edge of pterygoid flange) (2).

**Character 45** (Clark, 1994: char. 45): + Quadrate: without fenestrae (0), with single fenestrae (1), or with three or more fenestrae on dorsal and posteromedial surfaces (2).

**Character 46** (Clark, 1994: char. 46): Posterior edge of quadrate: broad medial to tympanum, gently concave (0), or posterior edge narrow dorsal to otoccipital contact, strongly concave (1).

**Character 47** (Clark, 1994: char. 47): Dorsal, primary head of quadrate articulates with: squamosal, otoccipital, and prootic (0), or with prootic and laterosphenoid (1).

**Character 48** (Clark, 1994: char. 48): Ventrolateral contact of otoccipital with quadrate: very narrow (0), or broad (1).

**Character 49** (Rummy et al., 2022: char 49, modified from Clark, 1994: char. 49): + Quadrate, squamosal, and otoccipital: do not meet to enclose cranioquadrate passage (0), enclose passage near lateral edge of skull (1), or meet lateral to the passage with otoccipital contacting the quadrate lateral to the posterior opening of the passage (2).

**Character 50** (Clark, 1994: char. 50): Pterygoid ramus of quadrate: with flat ventral edge (0), or with deep groove along ventral edge (1).

**Character 51** (Clark, 1994: char. 51): Ventromedial part of quadrate: does not contact otoccipital (0), or contacts otoccipital to enclose carotid artery and form passage for cranial nerves IX–XI (1).

**Character 52** (Clark, 1994: char. 52): Eustachian tubes: not enclosed between basioccipital and basisphenoid (0), or entirely enclosed (1).

**Character 53** (Clark, 1994: char. 53): Basisphenoid rostrum (cultriform process): slender (0), or dorsoventrally expanded (1).

**Character 54** (Clark, 1994: char. 54): Basipterygoid process: prominent, forming movable joint with pterygoid (0), or basipterygoid process small or absent, with basisphenoid joint suturally closed (1).

**Character 55** (modified from Clark, 1994: char. 55 by Ortega et al., 2000: char. 68): Basisphenoid ventral surface: shorter than the basioccipital (0), or wide and similar to, or longer in length than basioccipital (1).

**Character 56** (Clark, 1994: char. 56): Basisphenoid: exposed on ventral surface of braincase (0), or virtually excluded from ventral surface by pterygoid and basioccipital (1).

**Character 57** (Clark, 1994: char. 57): Basioccipital: without well-developed bilateral tuberosities (0), or with large pendulous tubera (1).

**Character 58** (Clark, 1994: char. 58): Otoccipital: without laterally concave descending flange ventral to subcapsular process (0), or with flange (1).

**Character 59** (Clark, 1994: char. 59): Cranial nerves IX–XI: pass through common large foramen vagi in otoccipital (0), or cranial nerve IX passes medial to nerves X and XI in separate passage (1).

**Character 60** (Clark, 1994: char. 60): Otoccipital: without large ventrolateral part ventral to paroccipital process (0), or with large ventrolateral part (1).

**Character 61** (Clark, 1994: char. 61): Crista interfenestralis between fenestrae pseudorotunda and ovalis nearly vertical (0), or horizontal (1).

**Character 62** (Clark, 1994: char. 62): Supraoccipital: forms dorsal edge of the foramen magnum (0), or otoccipitals broadly meet dorsal to the foramen magnum, separating supraoccipital from foramen (1).

**Character 63** (Clark, 1994: char. 63): Mastoid antrum: does not extend into supraoccipital (0), or extends through transverse canal in supraoccipital to connect middle ear regions (1).

**Character 64** (Clark, 1994: char. 64): Posterior surface of supraoccipital: nearly flat (0), or with bilateral posterior prominences (1).

**Character 65** (Rummy et al., 2022: char 65, modified from Clark, 1994: char. 65): + Palpebrals: absent (0), or one small palpebral present in orbit (1), or one large palpebral (2), or two large palpebrals (3).

**Character 66** (Clark, 1994: char. 66): External nares: divided by a septum (0), or confluent (1).

**Character 67** (Clark, 1994: char. 67): + Antorbital fenestra: as large as orbit (0), about half the diameter of the orbit (1), much smaller than the orbit (2), or absent (3).

**Character 68** (modified from Clark, 1994: char. 68 by Ortega et al., 2000: char. 41): Supratemporal fenestrae extension: relatively large, covering most of surface of skull roof (0), or relatively short, fenestrae surrounded by a flat and extended skull roof (1).

**Character 69** (Rummy et al., 2022: char 69, modified from Clark, 1994: char. 69): + Choanal groove: undivided (0), partly septated (1), or completely septated (2).

**Character 70** (Clark, 1994: char. 70): Dentary: extends posteriorly beneath mandibular fenestra (0), or does not extend beneath fenestra (1).

**Character 71** (Rummy et al., 2022: char 71, modified from Clark, 1994: char. 71): Retroarticular process: absent or extremely reduced (0), very short, broad, and robust (1), with an extensive rounded, wide, and flat (or slightly concave) surface projected posteroventrally and facing dorsomedially (2), posteriorly elongated, triangular shaped, and facing dorsally (3), or posteroventrally projecting and paddle shaped (4).

**Character 72** (Clark, 1994: char. 72): Prearticular: present (0), or absent (1).

**Character 73** (Rummy et al., 2022: char 73, modified from Clark, 1994: char. 73): + Articular without medial process (0), with short process not contacting braincase (1), or with process articulating with otoccipital and basisphenoid (2).

**Character 74** (Clark, 1994: char. 74): Dorsal edge of surangular: flat (0), or arched dorsally (1).

**Character 75** (Clark, 1994: char. 75): Mandibular fenestra: present (0), or absent (1).

**Character 76** (Clark, 1994: char. 76): Insertion area for M. pterygoideous posterior: does not extend onto lateral surface of angular (0), or extends onto lateral surface of angular (1).

**Character 77** (Rummy et al., 2022: char 77, modified from Clark, 1994: char. 77): + Splenial involvement in symphysis in ventral view: not involved (0), involved slightly in symphysis (1), or extensively involved (2).

**Character 78** (Clark, 1994: char. 78): Posterior premaxillary teeth: similar in size to anterior teeth (0), or much longer (1).

**Character 79** (Rummy et al., 2022: char 79, modified from Clark, 1994: char. 79): + Maxillary teeth waves: absent, no tooth-size variation (0), one wave of teeth enlarged (1), or enlarged maxillary teeth curved in two waves (festooned) (2).

**Character 80** (Clark, 1994: char. 80): Anterior dentary teeth opposite premaxilla-maxilla contact: no more than twice the length of other dentary teeth (0), or more than twice the length (1).

**Character 81** (Rummy et al., 2022: char 81, modified from Clark, 1994: char. 81): Dentary teeth posterior to tooth opposite premaxilla-maxilla contact: equal in size (0), or enlarged dentary teeth opposite to smaller teeth in maxillary tooth row (1).

**Character 82** (modified from Clark, 1994: char. 82 by Ortega et al., 2000: char. 120): Anterior and posterior scapular edges: symmetrical in lateral view (0), anterior edge more strongly concave than posterior edge (1), or dorsally narrow with straight edges (2).

**Character 83** (modified from Clark, 1994: char. 83 by Ortega et al., 2000: char. 121): Coracoid length: up to two-thirds of the scapular length (0), or subequal in length to scapula (1).

**Character 84** (Clark, 1994: char. 84): Anterior process of ilium: similar in length to posterior process (0), or one-quarter or less of the length of the posterior process (1).

**Character 85** (Clark, 1994: char. 85): Pubis: rodlike without expanded distal end (0), or with expanded distal end (1).

**Character 86** (Clark, 1994: char. 86): Pubis: forms anterior half of ventral edge of acetabulum (0), or pubis contacting the ilium but partially excluded from the acetabulum by the anterior process of the ischium (1), or pubis completely excluded from the acetabulum by the anterior process of the ischium (2).

**Character 87** (Clark, 1994: char. 87): Distal end of femur: with large lateral facet for the fibula (0), or with very small facet (1).

**Character 88** (Clark, 1994: char. 88): Fifth pedal digit: with phalanges (0), or without phalanges (1).

**Character 89** (Clark, 1994: char. 89): Atlas intercentrum: broader than long (0), or as long as broad (1).

**Character 90** (Rummy et al., 2022: char 90, modified from Clark, 1994: char. 90): Cervical neural spines: all anteroposteriorly large (0), only posterior ones rodlike (1), or all spines rodlike (2).

**Character 91** (modified from Clark, 1994: char. 91, by Buscalioni and Sanz, 1988: char. 37, and by Brochu, 1997a: char. 7): + Hypapophyses in cervicodorsal vertebrae: absent (0), present only in cervical vertebrae (1), present in cervical and the first two dorsal vertebrae (2), present up to the third dorsal vertebra (3), or up to the fourth dorsal vertebrae (4).

**Character 92** (Clark, 1994: char. 92): Cervical vertebrae: amphicoelous or amphyplatian (0), or procoelous (1).

**Character 93** (Clark, 1994: char. 93): Trunk vertebrae: amphicoelous or amphyplatian (0), or procoelous (1).

**Character 94** (Rummy et al., 2022: char 94, modified from Clark, 1994: char. 94): First caudal vertebrae: amphicoelous or amphy-platian (0), biconvex (1), or opisthocoelous (2), or procoelous (3).

**Character 95** (Clark, 1994: char. 95): Dorsal osteoderms, shape: rounded or ovate (0), or rectangular, broader than long (1), or square (2), or rectangular, longer than broad (3).

**Character 96** (Rummy et al., 2022: char 96, modified from Clark, 1994: char. 96; Brochu, 1997a: char. 40): Dorsal osteoderms: without articular anterior process (0), with a discrete convexity on anterior margin (1), or with a well-developed process located anterolaterally in dorsal parasagittal osteoderms (2).

**Character 97** (modified from Clark, 1994: char. 97 by Ortega et al., 2000: chars. 107 and 108 and Rummy et al., 2022): Rows of dorsal primary osteoderms (sensu Frey, 1988): two parallel rows (0), more than two (1), or more than four (2).

**Character 98** (Clark, 1994: char. 98): Osteoderms: some or all imbricated (0), or sutured to one another (1).

**Character 99** (Clark, 1994: char. 99): Tail osteoderms: dorsal only (0), or completely surrounded by osteoderms (1).

**Character 100** (Clark, 1994: char. 100): Trunk osteoderms: absent from ventral part of the trunk (0), or present (1).

**Character 101** (Clark, 1994: char. 101): Osteoderms: with longitudinal keels on dorsal surfaces (0), or without longitudinal keels (1).

**Character 102** (Wu and Sues, 1996: char. 14): Jugal: participating in margin of antorbital fossa (0), or separated from it (1).

**Character 103** (Rummy et al., 2022: char 103, modified from Wu and Sues, 1996: char. 17): Mandibular symphysis in lateral view: shallow and tapering anteriorly (0), deep and tapering anteriorly (1), deep and anteriorly convex (2), or shallow and anteriorly convex (3).

**Character 104** (Rummy et al., 2022: char 104, modified from Wu and Sues, 1996: char. 23): + Articular facet for quadrate condyle: equal in length to the quadrate condyles (0), slightly longer (1), or close to three times the length of the quadrate condyles (2).

**Character 105** (Rummy et al., 2022: char 105, modified from Wu and Sues, 1996: char. 24; Wu et al., 1997: char. 124): + Jaw joint: placed at level with basioccipital condyle (0), below basioccipital condyle about above level of lower tooth row (1), or below level of tooth row (2).

**Character 106** (Rummy et al., 2022: char 106, modified from Wu and Sues, 1996: char. 27 and Ortega et al., 2000: char. 133): + Premaxillary tooth number: six (0); five (1); four (2); three (3); two (4); one (5).

**Character 107** (Rummy et al., 2022: char 107, modified from Wu and Sues, 1996: char. 29): Unsculptured region along alveolar margin on lateral surface of maxilla: absent (0), or present (1).

**Character 108** (Wu and Sues, 1996: char. 30): + Maxilla: with eight or more teeth (0), seven (1), six (2), five (3), or four teeth (1).

**Character 109** (Wu and Sues, 1996: char. 33): Coracoid, posteromedial or ventromedial process: absent (0); elongate posteromedial process present (1); distally expanded ventromedial process present (2).

**Character 110** (Wu and Sues, 1996: char. 40): Radiale and ulnare: short and massive (0), or elongate (1).

**Character 111** (Rummy et al., 2022: char 111, modified from Gomani, 1997: char. 4): Prefrontals anterior to orbits: elongated, oriented parallel to anteroposterior axis of the skull (0), or short and broad, oriented posteromedially-anterolaterally (1).

**Character 112** (Rummy et al., 2022: char 112, modified from Gomani, 1997: char. 32): Basioccipital and ventral part of otoccipital: facing posteriorly (0), or posteroventrally (1).

**Character 113** (Buscalioni and Sanz, 1988: char. 35): Vertebral centra: cylindrical (0), or spool shaped (1).

**Character 114** (Rummy et al., 2022: char 114, modified from Buscalioni and Sanz, 1988: char. 39): Transverse process of posterior dorsal vertebrae dorsoventrally low and laminar (0), or dorsoventrally high (1).

**Character 115** (Buscalioni and Sanz, 1988: char. 44): Number of sacral vertebrae: two (0), or more than two (1).

**Character 116** (Buscalioni and Sanz, 1988: char. 49): Supra-acetabular crest: present (0), or absent (1).

**Character 117** (Buscalioni and Sanz, 1988: char. 54): Proximal end of radiale expanded symmetrically, similarly to the distal end (0), or more expanded proximolaterally than proximomedially (1).

**Character 118** (Ortega et al., 1996: char. 5): Lateral surface of the anterior region of surangular and posterior region of dentary: without a longitudinal depression (0), or with a longitudinal depression (1).

**Character 119** (Ortega et al., 1996: char. 9): Ventral exposure of splenials: absent (0), or present (1).

**Character 120** (Rummy et al., 2022: char 120, modified from Andrade and Bertini, 2008a: char. 120; Ortega et al., 1996: char. 11; Ortega et al., 2000: char. 100): Tooth margin carinae: without carinae or with smooth or crenulated carinae (0); with homogeneous denticulate carinae (denticles are small and symmetrical in form as in ziphodont teeth (1); with heterogeneous carinae possessing rounded tubercle-like denticles, developed preferentially along posterior margin (2).

**Character 121** (Rummy et al., 2022: char 121, modified from Pol, 1999a: char. 133; Ortega et al., 2000: char. 145): Lateral surface of anterior process of jugal: flat or convex (0), or with broad shelf below the orbit with triangular depression underneath it (1).

**Character 122** (Pol, 1999a: char. 134): Jugal: does not exceed the anterior margin of orbit (0), or exceeds margin (1).

**Character 123** (Pol, 1999a: char. 135): Notch in premaxilla on lateral edge of external nares: absent (0), or present on the dorsal half of the external nares lateral margin (1).

**Character 124** (Pol, 1999a: char. 136): Dorsal border of external nares: formed mostly by the nasals (0), or by both the nasals and premaxilla (1).

**Character 125** (Pol, 1999a: char. 138): Posterodorsal process of premaxilla: absent (0), or present extending posteriorly wedging between maxilla and nasals (1).

**Character 126** (Pol, 1999a: char. 139 and Ortega et al., 2000: char. 9): + Premaxilla- maxilla suture in palatal view, medial to alveolar region, orientation of suture: anteromedially directed (0); sinusoidal, posteromedially directed on its lateral half and anteromedially directed along its medial region (1); posteromedially directed (2); straight (3); posteromedially curved (U shaped) (4).

**Character 127** (Pol, 1999a: char. 140): Nasal lateral border posterior to external nares: laterally concave (0), or straight (1).

**Character 128** (Pol, 1999a: char. 141): Nasal lateral edges: nearly parallel (0), oblique to each other converging anteriorly (1), or oblique to each other diverging anteriorly (2).

**Character 129** (Pol, 1999a: char. 143): Palatine anteromedial margin: exceeding the anterior mar- gin of the palatal fenestrae extending anteriorly between the maxillae (0), or not exceeding the anterior margin of palatal fenestrae (1).

**Character 130** (Pol, 1999a: char. 144): Dorsoventral height of jugal antorbital region respect to infra- orbital region: equal or lower (0), or antorbital region more expanded than infraorbital region of jugal (1).

**Character 131** (Pol, 1999a: char. 145): Maxilla- lacrimal contact: partly included in antorbital fossa (0), or completely included (1).

**Character 132** (Pol, 1999a: char. 146): Lateral eustachian tube openings: located posteriorly to the medial opening (0), or aligned anteroposteriorly and dorsoventrally (1).

**Character 133** (Pol, 1999a: char. 147): Anterior process of ectopterygoid: developed (0), or reduced to absent (1).

**Character 134** (Pol, 1999a: char. 148): Posterior process of ectopterygoid: developed (0), or reduced to absent (1).

**Character 135** (Pol, 1999a: char. 149; Ortega et al., 2000: char. 13): Small foramen located in the premaxillo-maxillary suture in lateral surface (not for big mandibular teeth): absent (0), or present (1).

**Character 136** (Pol, 1999a: char. 150): Jugal posterior process, extent of process: exceeding posteriorly the infratemporal fenestrae (0); does not exceed infratemporal fenestrae (1).

**Character 137** (Pol, 1999a: char. 151): Compressed crown of maxillary teeth: oriented parallel to the longitudinal axis of skull (0), or obliquely disposed (1).

**Character 138** (Pol, 1999a: char. 152): Large and aligned neurovascular foramina on lateral maxillary surface: absent (0), or present (1).

**Character 139** (Rummy et al., 2022: char 139, modified from Pol, 1999a: char. 153): External surface of maxilla and premaxilla, general shape: with a single plane facing laterally (0); with ventral region facing laterally and dorsal region facing dorsolaterally (1).

**Character 140** (Rummy et al., 2022: char 140, modified from Pol, 1999a: char. 154; Ortega et al., 2000: char. 104; Andrade and Bertini, 2008a: char. 135): +Maxillary teeth, lateral compression: absent (0); present, compression assymetrically occuring only along distal margin giving teeth a teardrop shape (1); present, lateral compression symmetrically developed (2).

**Character 141** (Pol, 1999a: char. 155): Posteroventral corner of quadratojugal: reaches the quadrate condyles (0); does not reach the quadrate condyles (1).

**Character 142** (Rummy et al., 2022: char 142, modified from Pol, 1999a: char. 156): + Base of postorbital process of jugal: directed posterodorsally (0), or dorsally (1), or anterodorsally (2).

**Character 143** (Pol, 1999a: char. 157): + Postorbital process of jugal, location on jugal: anteriorly placed (0); in the middle (1); posteriorly places (2).

**Character 144** (Pol, 1999a: char. 158; Ortega et al., 2000: char. 36): Postorbital-ectopterygoid contact: present (0), or absent (1).

**Character 145** (Pol, 1999a: char. 161): Quadratojugal, ornamentation: absent (0); ornamented on the base (1).

**Character 146** (Pol, 1999a: char. 162): Prefrontal- maxillary contact in the inner anteromedial region of orbit absent (0); present (1).

**Character 147** (Pol, 1999a: char. 163): Basisphenoid, exposure on braincase: without lateral exposure (0); with lateral exposure (1).

**Character 148** (Pol, 1999a: char. 165): Quadrate process of pterygoids: well developed (0), or poorly developed (1).

**Character 149** (Rummy et al., 2022: char 149, modified from Pol, 1999a: char. 166 and Ortega et al., 2000: char. 44): + Quadrate major axis, direction of orientation: posteroventrally (0); ventrally (1); anteroventrally (2).

**Character 150** (Pol, 1999a: char. 167): Quadrate distal end: with only one plane facing posteriorly (0), or with two distinct faces in posterior view, a posterior one and a medial one bearing the foramen aereum (1).

**Character 151** (Pol, 1999a: char. 168): Anteroposterior development of neural spine in axis: well developed covering all the neural arch length (0); poorly developed, located over the posterior half of the neural arch (1).

**Character 152** (Pol, 1999a: char. 169): Prezygapophyses of axis, development relative to neural arch: not exceeding edge of neural arch (0); exceeding the anterior margin of neural arch (1).

**Character 153** (Pol, 1999a: char. 170): Postzygapophyses of axis: well developed, curved laterally (0); poorly developed (1).

**Character 154** (Rummy et al., 2022: char 154, modified from Pol, 1999b: char. 212): Shape of dentary symphysis in ventral view: tapering anteriorly forming an angle (0), U-shaped, smoothly curving anteriorly (1); lateral edges longitudinally oriented, convex anterolateral corner, and extensive transversally oriented anterior edge (2).

**Character 155** (Pol, 1999b: char. 213): Unsculpted region in the dentary below the tooth row: absent (0); present (1).

**Character 156** (Buckley and Brochu, 1999: char. 102): Surangular, contribution to the glenoid fossa: forms only the lateral wall of glenoid (0); forms approximately one-third of the glenoid (1).

**Character 157** (Rummy et al., 2022: char 157, modified from Buckley and Brochu, 1999: char. 102): Femur, anterior margin: linear (0); bears flange for coccygeofemoralis musculature (1).

**Character 158** (Rummy et al., 2022: char 158, modified from Buckley and Brochu, 1999: char. 105): Dentary, lateral surface: smooth lateral to seventh alveolous (0); with lateral concavity for the reception of the enlarged maxillary tooth (1).

**Character 159** (Rummy et al., 2022: char 159, modified from Ortega et al., 1996: char. 1 and Buckley and Brochu, 1999: char. 107): Dorsal edge of dentary: slightly concave or straight and subparallel to the longitudinal axis of skull (0); straight with an abrupt dorsal expansion, being straight posteriorly (1); with a single dorsal expansion and concave posterior to this (2); sinusoidal, with two concave waves (3).

**Character 160** (Rummy et al., 2022: char 160, modified from Ortega et al., 1996: char. 2 and Buckley and Brochu, 1999: char. 108): Dentary compression and lateroventral surface anterior to mandibular fenestra: compressed and vertical (0); not compressed and convex (1).

**Character 161** (Ortega et al., 1996: char. 7 and Buckley and Brochu, 1999: char. 110): Splenial posterior to symphysis: thin (0); robust dorsally (1).

**Character 162** (Ortega et al., 1996: char. 13; Buckley et al., 2000: char. 117): Cheek teeth: not constricted at base of crown (0); constricted (1).

**Character 163** (Ortega et al., 2000: char. 10): Ventral edge of premaxilla, location relative to maxilla: at the same height as the ventral edge of maxilla (0); located deeper, with the dorsal contour of anterior part of dentary strongly concave (1).

**Character 164** (Rummy et al., 2022: char 164, modified from Ortega et al., 2000: char. 19): Maxillary dental implantation: teeth in isolated alveoli (0); located on a dental groove (1).

**Character 165** (Ortega et al., 2000: char. 24): Caudal tip of nasals: converge at sagittal plane (0); caudally separated by anterior sagittal projection of frontals (1).

**Character 166** (Ortega et al., 2000: char. 33): Relative length between squamosal and postorbital: squamosal is longer (0); postorbital is longer (1).

**Character 167** (Rummy et al., 2022: char 167, modified from Ortega et al., 2000: character 34): + Jugal portion of postorbital bar, relative to lateral surface of jugal: flush with lateral surface (0); anteriorly continuous but posteriorly inset (1); medially displaced and a ridge separate postorbital bar from lateral surface of jugal (2).

**Character 168** (Ortega et al., 2000: char. 42): Outer surface of squamosal laterodorsally oriented: extensive (0); reduced and sculpted (1); reduced and unsculpted (2).

**Character 169** (Ortega et al., 2000: char. 47): Quadratojugal spine at caudal margin of infratemporal fenestra: absent (0), or present (1).

**Character 170** (Rummy et al., 2022: char 170, modified from Ortega et al., 2000: char. 53): Quadrate condyles: poorly developed intercondylar groove (0); medial condyle expands ventrally, being separate from the lateral condyle by a deep intercondylar groove (1).

**Character 171** (Ortega et al., 2000: char. 62): Exposure of supraoccipital in skull roof: absent (0); present (1).

**Character 172** (Ortega et al., 2000: char. 70): Nasal participation in antorbital fenestra: yes (0); no (1).

**Character 173** (Ortega et al., 2000: char. 75): Anterior opening of temporo-orbital canal, in dorsal: exposed (0); hidden in dorsal view and overlapped by squamosal rim of supratemporal fossa (1).

**Character 174** (Ortega et al., 2000: char. 90): Foramen intramandibularis oralis: small or absent (0); big and slotlike (1).

**Character 175** (Rummy et al., 2022: char 175, modified from Ortega et al., 2000: char 98): Coronoid size: short and located below the dorsal edge of the mandibular ramus (0); anteriorly extended with posterior region elevated at the dorsal margin of the mandibular ramus (1).

**Character 176** (Ortega et al., 2000: char. 101): Width of root of teeth respect to crown: narrower or equal (0); wider (1).

**Character 177** (Ortega et al., 2000: char. 109): Gap in cervico-thoracic dorsal armor: absent (0); present (1).

**Character 178** (Ortega et al., 2000: char. 130): Lateral contour of snout in dorsal view: straight (0); sinusoidal (1).

**Character 179** (Ortega et al., 2000: char. 138): Pterygoidean flanges: laminar and expanded (0); barlike and elongated (1); barlike and poorly developed (2).

**Character 180** (Ortega et al., 2000: char. 146): Ectopterygoid medial process: single (0); forked (1).

**Character 181** (Rummy et al., 2022: char 181, modified from Ortega et al., 2000: char. 157): Skull roof, shape in dorsal view: rectangular (0); trapezoidal (1).

**Character 182** (Ortega et al., 2000: char. 30): + Prefrontal pillars when integrated in palate: pillars transversely expanded (0); transversely expanded in their dorsal part and columnar ventrally (1); longitudinally expanded in their dorsal part and columnar ventrally (2).

**Character 183** (Ortega et al., 2000: char. 21): Ventral edge of maxilla in lateral view: straight or convex (0); sinusoidal (1).

**Character 184** (Rummy et al., 2022: char 184, modified from Ortega et al., 2000: char. 156): Position of first enlarged maxillary teeth: second or third alveoli (0); fourth or fifth (1).

**Character 185** (Pol and Apesteguia, 2005: char. 180): Splenial-dentary suture at symphysis on ventral surface: v-shaped (0); transversal (1).

**Character 186** (Pol and Apesteguia, 2005: char. 181): Posterior peg at symphysis: absent (0), or present (1).

**Character 187** (Pol and Apesteguia, 2005: char. 182): Posterior ridge on glenoid fossa of articular: present (0), or absent (1)

**Character 188** (Rummy et al., 2022: char 188, modified from Pol et al., 2009; previously from Gomani, 1997: char. 46; Buckley et al., 2000: char. 113; Andrade and Bertini, 2008a: char 149): Cusps of teeth, number and conformation: one unique cusp (0); one main cusp with smaller cusps arranged in one row (1); one main cusp with smaller cusps arranged in more than one row (2); several cusps of equal size arranged in more than one row (3); multiple small cusps along edges of occlusal surface (4).

**Character 189** (Pol and Apesteguia, 2005: char. 184): Dorsal surface of mandibular symphysis: flat or slightly concave (0); strongly concave and narrow, trough shaped (1)

**Character 190** (Pol and Apesteguia, 2005: char. 185): Medial surface of splenials posterior to symphysis: flat or slightly convex (0); markedly concave (1).

**Character 191** (Pol and Apesteguía, 2005: char. 186): Choanal septum shape: narrow vertical bony sheet (0); T-shaped bar expanded ventrally (1); massive and blocky (2).

**Character 192** (Pol and Norell, 2004a: char. 164): Cross section of distal end of quadrate: mediolaterally wide and anteroposteriorly thin (0); subquadrangular (1).

**Character 193** (Pol and Apesteguía, 2005: char. 188): Lateral surface of dentaries below alveolar margin, at mid to posterior region of tooth row: vertically oriented, continuous with rest of lateral surface of the dentaries (0); flat surface exposed laterodorsally, divided by a ridge from rest of the lateral surface of the dentaries (1).

**Character 194** (Pol and Norell, 2004a: char. 165): Palatine-pterygoid contact on palate: palatines overlies pterygoids (0); palatines firmly sutured to pterygoids (1).

**Character 195** (Pol et al., 2004: char. 164): Ectopterygoid main axis oriented: laterally or slightly anterolaterally (0); anteriorly, subparallel to the skull longitudinal axis (1).

**Character 196** (Wu et al., 1997: char. 103): Squamosal descending process: absent (0), or present (1).

**Character 197** (Rummy et al., 2022: char 197, modified from Wu et al., 1997: char. 105): + Development of distal quadrate body ventral to otoccipital-quadrate contact: distinct (0), incipiently distinct (1), or indistinct (2).

**Character 198** (Wu et al., 1997: char. 106): Pterygoid flanges: thin and laminar (0); dorsoventrally thick, with pneumatic spaces (1).

**Character 199** (Wu et al., 1997: char. 108): Postorbital participation in infratemporal fenestra: almost or entirely excluded (0); bordering infratemporal fenestra (1).

**Character 200** (Wu et al., 1997: char. 109): Palatines: form margin of suborbital fenestra (0), or excluded from margin of suborbital fenestra (1).

**Character 201** (Wu et al., 1997: char. 110): Angular posterior to mandibular fenestra: widely exposed on lateral surface of mandible (0); shifted to the ventral surface of mandible (1).

**Character 202** (Wu et al., 1997: char. 112): Posteroventral edge of mandibular ramus, shape: straight or convex (0); markedly deflected (1).

**Character 203** (Rummy et al., 2022: char 203, modified from Wu et al., 1997: char. 119): Quadrate ramus of pterygoid, width in ventral view: narrow (0); broad (1).

**Character 204** (Wu et al., 1997: char. 121): Pterygoids, contact on palate: not in contact anterior to basisphenoid on palate (0); pterygoids in contact (1).

**Character 205** (Rummy et al., 2022: char 205, modified from Wu et al., 1997: char. 122): Olecranon: well developed (0); absent (1).

**Character 206** (Wu et al., 1997: char. 123): Cranial table width with respect to ventral portion of skull: as wide as ventral portion (0); narrower than ventral portion of skull (1).

**Character 207** (Wu et al., 1997: char. 127): Depression on posterolateral surface of maxilla: absent (0); present (1).

**Character 208** (Rummy et al., 2022: char 208, modified from Wu et al., 1997: char. 128 by Pol et al., 2009): Paired anterior palatal fenestra: absent (0); present (1).

**Character 209**(Pol and Norell, 2004a: char. 179): Paired ridges located medially on ventral surface of basisphenoid: absent (0); present (1).

**Character 210** (Pol et al., 2004: char. 179): Ventral margin of infratemporal bar of jugal: straight (0); dorsally arched (1).

**Character 211** (Pol and Norell, 2004a: char. 180): Posterolateral end of quadratojugal, shape and relationship with quadrate: acute or rounded, tightly overlapping the quadrate (0); with sinusoidal ventral edge and wide and rounded posterior edge slightly overhanging the lateral surface of the quadrate (1).

**Character 212** (Pol and Norell, 2004a : char. 181): Quadrate body distal to otoccipital- quadrate, orientation of contact in posterior view: ventrally oriented (0); ventrolaterally oriented (1).

**Character 213** (Gasparini et al., 1993: char. 3): Wedge-like process of the maxilla in lateral surface of premaxilla-maxilla: absent (0); present (1).

**Character 214** (Pol and Norell, 2004b: char. 181): Palpebrals: separated from the lateral edge of the frontal (0); extensively sutured to each other and to the lateral margin of the frontals (1).

**Character 215** (Pol and Norell, 2004b: char. 182): External surface of ascending process of jugal: exposed laterally (0); exposed posterolaterally (1).

**Character 216** (modified from Pol and Norell, 2004b: char. 183 by Pol et al., 2009 and Turner and Sertich, 2010): Longitudinal ridge on lateral surface of jugal below infratemporal fenestra: absent (0); present, running entire length of posterior process of jugal (1); present, running entire length of jugal (2).

**Character 217** (Pol and Norell, 2004b: char. 184): Dorsal surface of posterolateral region of squamosal: without ridges (0); with three curved ridges oriented longitudinally (1).

**Character 218** (Pol and Norell, 2004b: char. 185): Ridge along dorsal section of quadrate-quadratojugal contact: absent (0); present (1).

**Character 219** (modified from Pol and Norell, 2004b: char. 186 by Pol et al., 2009): Sharp ridge on the surface of the angular: absent (0); present on the ventral-most margin (1); present along the lateral surface (2).

**Character 220** (Pol and Norell, 2004b: char. 187): Longitudinal ridge along the dorsolateral surface of surangular: absent (0); present (1).

**Character 221** (Pol and Norell, 2004b: char. 188): Dorsal surface of osteoderms ornamented with anterolaterally and anteromedially directed ridges (fleur de lys pattern of Osmo ́lska et al., 1997): absent (0), or present (1).

**Character 222** (Pol and Norell, 2004b: char. 189): Cervical region surrounded by lateral and ventral osteoderms sutured to the dorsal elements: absent (0), or present (1).

**Character 223** (Pol and Norell, 2004b: char. 190): Appendicular osteoderms: absent (0), or present (1).

**Character 224** (Ortega et al., 2000: character 72): Supratemporal fenestra: present (0), or absent (1).

**Character 225** (Rummy et al., 2022: char 225, modified from Pol and Apesteguia, 2005: char. 220): Flat ventral surface of internal nares septum: parallel sided (0), or tapering anteriorly (1), or expanding anteriorly (2).

**Character 226** (Rummy et al., 2022: char 226, modified from Pol and Apesteguía, 2005: char. 221): + Perinarial fossa: restricted extension (0); extensive, with distinctly concave surface facing anteriorly (1); large concave surface facing anteriorly, projecting anteroventrally from external nares and opening toward alveolar margin (2); or extremely large and well-developed, occupying nearly entire surface of premaxilla ventral to external naris (3).

**Character 227** (Rummy et al., 2022: char 227, modified from Sereno et al., 2001: char. 67): Premaxillary palate, circular paramedian depressions: absent (0); present, located anteriorly on the premaxilla (1); present, located at the premaxilla-maxilla suture (2).

**Character 228** (Pol and Gasparini, 2009: char. 223): Posterolateral region of nasals: flat surface facing dorsally (0); lateral region deflected ventrally, forming part of the lateral surface of the snout (1).

**Character 229** (Zaher et al., 2006: char. 193): Lacrimal, posterior extent and relationship with jugal: extends ventroposteriorly, widely contacting the jugal (0); tapers ventroposteriorly, does not contact or contacts the jugal only slightly (1).

**Character 230** (Zaher et al., 2006: char. 194): Jugal, large foramen on the lateral surface near the anterior margin: absent (0); present (1).

**Character 231** (Rummy et al., 2022: char 231, modified from Zaher et al., 2006: char. 195): Procumbent premaxillary: alveoli absent (0); present (1).

**Character 232** (modified from Martinelli, 2003: char. 36 by Zaher et al., 2006: char. 196 and Turner, 2004: char. 119): Palatines, orientation: run parasagittally along midline (0); diverge laterally becoming rodlike caudally forming palatine bars (1).

**Character 233** (Zaher et al., 2006: char. 197): Ectopterygoid, participation in the palatine bar: absent (0); present (1).

**Character 234** (Pol and Norell, 2004a: char. 183): Choanal opening: opened posteriorly and continuous with pterygoid surface (0); closed posteriorly by an elevated wall formed by the pterygoids (1).

**Character 235** (Zaher et al., 2006: char. 198): Ectopterygoid, extent of medial projection on the ventral surface of pterygoid flanges: barely extended (0); widely extended covering approximately the lateral half of the ventral surface of the pterygoid flanges (1).

**Character 236** (Gasparini et al., 2006: char. 236): Evaginated maxillary alveolar edges: absent (0); present as a continuous sheet (1); present as discrete evaginations at each alveoli (2).

**Character 237** (Gasparini et al., 2006: char. 237): Premaxilla, foramen in perinarial depression: absent (0); present (1).

**Character 238** (Sereno et al., 2001: char. 27): Frontal, anterior ramus with respect to the tip of the prefrontal: ending posteriorly (0); ending anteriorly (1).

**Character 239** (Rummy et al., 2022: char 239, modified from Sereno et al., 2001: char. 68): Premaxilla, anterior alveolar margin orientation: vertical (0); inturned (1).

**Character 240** (Rummy et al., 2022: char 240, modified from Sereno et al., 2001: char. 69): Premaxillary tooth row orientation: arched posteriorly from midline (0); angled posterolaterally, at 120-degree angle (1); transverse (2).

**Character 241** (Sereno et al., 2001: char. 70): Last premaxillary tooth position relative to tooth row: anterior (0); anterolateral (1).

**Character 242** (Pol and Gasparini, 2009: char. 242): Posterior teeth with rings of undulating enamel: absent (0); present (1).

**Character 243** (modified from Brochu, 1999: char. 108 by Gasparini et al., 2006: char. 243): Maxilla-palatine suture, shape of palatines: palatine anteriorly rounded (0); palatine anteriorly pointed (1); palatine invaginated (2).

**Character 244** (Gasparini et al., 2006: char. 244): Postorbital bar, lateral surface formed by: postorbital and jugal (0); only by postorbital (1).

**Character 245** (Gasparini et al., 2006: char. 245): Surangular groove, enlarged foramen at anterior end: absent (0); present (1).

**Character 246** (Pol and Gasparini, 2009: char. 246): Shape of antorbital fossa: subcircular or subtriangular (0); elongated, low, and oriented obliquely (1).

**Character 247** (Pol and Gasparini, 2009: char. 247): Prefrontal lateral development: reduced (0); enlarged, extending laterally over the orbit (1).

**Character 248** (Pol and Gasparini, 2009: char. 248): Foramen for the internal carotid artery: reduced, similar in size to the openings for cranial nerves IX–XI (0); extremely enlarged (1).

**Character 249** (Pol and Gasparini, 2009: char. 249): Squamosal posterolateral region, lateral to paroccipital process: narrow (0); bearing a sub- rounded subcircular flat surface (1).

**Character 250** (Gasparini et al., 2006: char. 250): Posteromedial branch of squamosal, orientation: transversely oriented (0); posterolaterally oriented (1).

**Character 251** (Pol and Gasparini, 2009: char. 251): Dorsal margin of squamosal occipital flange: straight (0); dorsally concave (1).

**Character 252** (Pol and Gasparini, 2009: char. 252): Sculpture in external surface of rostrum: absent (0); present (1).

**Character 253** (Gasparini et al., 2006: char. 253): Longitudinal depressions on palatal surface of maxillae and palatines: absent (0); present (1).

**Character 254** (Pol and Gasparini, 2009: char. 254): Angle between medial and anterior margins of supratemporal fossa: approximately 90° (0); approximately 45° (1).

**Character 255** (Gasparini et al., 2006: char. 255): Sacral vertebrae, direction of transverse processes: laterally (0); markedly deflected ventrally (1).

**Character 256** (Pol and Gasparini, 2009: char. 256): Prefrontal and lacrimal around orbits: forming flat rims (0); evaginated, forming elevated rims (1).

**Character 257** (Pol and Gasparini, 2009: char. 257): Nasal bones: paired (0); partially or completely fused (1).

**Character 258** (Brochu, 1997a: char. 3): Axial neural spines, width of posterior half: wide (0); narrow (1).

**Character 259** (Brochu, 1997a: char. 19): Axial hyapophysis, deep fork: present (0); absent (1).

**Character 260** (Brochu, 1997a: char. 27): Ulna, width of olecranon process: narrow and subangular (0); wide and rounded (1).

**Character 261** (Brochu, 1997a: char. 29): M. teres major and M. dorsalis scapulae insert separately on humerus: scars can be distinguished dorsal to deltopectoral crest (0); insert with common tendon, single insertion scar (1).

**Character 262** (Rummy et al., 2022: char 262, modified from Brochu, 1997a: char. 53): Dentary, projection of anterior alveoli: anterodorsally (0); weakly procumbent (1); strongly procumbent (2).

**Character 263** (Brochu, 1997a: char. 84): Squamosal, dorsal and ventral rims of squamosal groove for external ear valve musculature: parallel (0); or squamosal groove flares anteriorly (1).

**Character 264** (Brochu, 1997a: char. 91): Ectopterygoid, contact with maxilla near toothrow: ectopterygoid abuts maxillary toothrow (0); maxilla broadly separates ectopterygoid from maxillary toothrow (1).

**Character 265** (Brochu, 1997a: char. 92): Shallow fossa at anteromedial corner of supratemporal fenestra: present (0); absent, anteromedial corner of supratemporal fenestra smooth (1).

**Character 266** (Rummy et al., 2022: char 266, modified from Brochu, 1997: char. 103): Lateral margins of frontal: flush with skull surface (0); elevated, forming ridged orbital margins (1).

**Character 267** (Brochu, 1997a: char. 130): Laterosphenoid, orientation of capitate process: laterally oriented (0); anteroposteriorly oriented toward midline (1).

**Character 268** (modified from Brochu, 1997a: char. 141 by Pol et al., 2009): Exoccipital, development of boss and paroccipital process: boss prominent on paroccipital process, process lateral to cranioquadrate opening short (0); boss small or absent on paroccipital process, process lateral to cranioquadrate opening long (1).

**Character 269** (modified from Norell, 1988: char. 32 by Brochu, 1997a: char. 149): Ectopterygoid, extent along lateral pterygoid flange, at maturity: extends to posterior tip of lateral pterygoid flange (0); does not extend to posterior tip of lateral pterygoid (1).

**Character 270** (Rummy et al., 2022: char 270, modified from Brochu, 1997a: char. 153): Incisive foramen, location relative to premaxillary toothrow: foramen situated far from premaxillary toothrow, at the level of the second or third alveolus (0); abuts premaxillary toothrow (1); projects between first premaxillary teeth (2).

**Character 271** (modified from Turner, 2006: character 126 by Pol et al., 2009 by Pritchard et al., 2013): Ventral surface of choanal septum: smooth to slightly depressed (0); marked by a acute groove (1); vomeral septum divided by into bilateral laminae (2).

**Character 272** (modified from Turner, 2006 char. 128 by Pol et al., 2009): Proximalmost portion of fibular head: straight sided to weakly developed posteriorly (0); very sharply projecting posteriorly, forming distinct extension (1).

**Character 273** (Rummy et al., 2022: char 273, modified from Turner, 2006: char. 129): Cervical rib shaft, posterior process, posterodorsally projecting spine at the junction with the tubercular process: absent (0); present (1); State 2 (2).

**Character 274** (Rummy et al., 2022: char 274, modified from Pol et al., 2009: char. 274): Longitudinal keels on dorsal surface of osteoderms: restricted to the posterior edge of osteoderm (0); not restricted to the posterior edge (1).

**Character 275** (Pol et al., 2009: char. 275): Jugal, anteriorly on lateral surface below orbits: lacks a depression (0); possesses a depression (1).

**Character 276** (modified from Pol et al., 2009: char. 276 by Pritchard et al., 2013): Transverse ridge crossing the frontal anteromedial to the orbits: absent (0); present as a ridge (1); present as prominent anteriorly curved shelf (transverse interorbital crest sensu Andrade and Hornung 2011) (2); present as anteroposteriorly oriented crest on frontal (3).

**Character 277** (Pol et al., 2009: char. 277): Shallow hemispherical depression on the lacrimal and/or prefrontal anterior to the orbital margin (not articulation facet for palpebral): absent (0); present (1).

**Character 278** (Pol et al., 2009: char. 278): Anterior half of interfenestral bar between suborbital fenestrae: lateral margins are parallel to subparallel (0); flared anteriorly (1).

**Character 279** (Pol et al., 2009: char. 279): Posterior half of interfenestral bar between suborbital fenestrae: lateral margins are parallel to subparallel (0); flared posteriorly (1).

**Character 280** (Pol et al., 2009: char. 280): Angular, shape of posteroventral margin: straight or gently arched dorsally (0); strongly arched dorsally (1).

**Character 281** (Pol et al., 2009: char. 281): Squamosal, lateral margin of dorsal surface: squared off with continuous ear valve groove (0); bears a prominent depressed area just anterior to the posterior lobe of the squamosal, groove for ear valve discontinuous (1).

**Character 282** (Pol et al., 2009: char. 282): Fibula, shaft distal to iliofibularis trochanter: straight (0); bowed posteriorly (1).

**Character 283** (Buckley and Brochu, 1999: char. 106): Scapular blade width: no more than twice the length of the scapulocoracoid articulation (0); scapular blade very broad and greater than twice the length of the scapulocoracoid articulation (1).

**Character 284** (modified from Buckley et al., 2000: char. 115 by Pritchard et al., 2013): Vomer, exposure on palate: vomer contributes flattened plate to secondary palate (0); vomer forms no part of secondary palate (1).

**Character 285** (Turner and Buckley, 2008: char. 285): Supraoccipital, when present on dorsal skull roof: with narrow exposure, parietal forms portion of occipital surface (0); with broad exposure, parietal does not form portion of occipital surface (1).

**Character 286** (Turner and Buckley, 2008: char. 286): Jugal, anterior and posterior processes: inline dorsoventrally (0); anterior and posterior processes at a sharp angle to one another, both processes slope ventrally to form a strongly arched jugal (1).

**Character 287** (Turner and Buckley, 2008: char. 287): Lateral expansion of posterodorsal edge of surangular anterior to glenoid fossa: absent (0); present (1).

**Character 288** (Turner and Buckley, 2008: char. 288): In lateral view, anterior process of the squamosal extending to the orbital margin, overlapping the postorbital: absent (0); present (1).

**Character 289** (Turner and Buckley, 2008: char. 289): In lateral view, surangular and dentary suture: simple with little or no interdigitating (0); suture complex with interlocking prongs from both surangular and dentary, three posterior prongs from the dentary and two form the surangular (1).

**Character 290** (Turner and Buckley, 2008: char. 290): Prominent depression on the palate near alveolar margin at the level of the 6th or 7th alveoulus: absent (0); present (1).

**Character 291** (Andrade and Bertini, 2008a: char. 103): Pterygoid, ventral surface of pterygoid flanges, parachoanal fossae: absent (0); present (1).

**Character 292** (Turner and Sertich, 2010: char. 292): Pterygoid, in ventral view, participation in the suborbital fenestra: pterygoid forms margin of suborbital fenestra (0); excluded from suborbital fenestra (1).

**Character 293** (Turner and Sertich, 2010: char. 293): Maxilla, lateral surface along alveolar margin, conformation of the neurovascular foramina: foramina absent or form a single continuous row (0); gap in foramina between an anterior series and a posterior series (1).

**Character 294** (Turner and Sertich, 2010: char. 294): Surface of tooth enamel: smooth or slightly crenulated (0); with ridges at base of crown (1).

**Character 295** (Turner and Sertich, 2010: char. 295): Posterior (molariform) teeth, wear facets: absent (0); present (1).

**Character 296** (Turner and Sertich, 2010: char. 296): Tooth (with transitional morphology) present at premax-max contact absent (0); present (1).

**Character 297** (Turner and Sertich, 2010: char. 297): Basiocciptal, midline crest on basioccipital plate below occipital condyle absent (0); present (1).

**Character 298** (Turner and Sertich, 2010: char. 298): Dorsal osteoderms, accessory ranges of osteoderms (sensu Frey, 1988) absent (0); present (1).

**Character 299** (Andrade and Bertini, 2008a: char. 131): Maxillary tooth, size relative to maxillary palatal surface in palatal view proportionally small teeth, occupying only marginal portion of ventral surface of maxilla (0); proportionally well-developed teeth, occupying large area of maxillary palatal surface (1).

**Character 300** (Jouve, 2009: character 75; Jouve, 2004: character 68): Ventral lamina of jugal extends far anterior to the ectopterygoid (0); ends at the level of the ectopterygoid (1).

**Character 301** (Sereno and Larsson, 2009: character 199; adapted from Norell, 1988: character 42 and Brochu, 1997: character 51): Surangular extension toward posterior end of retroarticular process along entire length (0); pinched off anterior to posterior tip (1).

**Character 302** (Turner, 2015: char. 302) Muscle attachment scars on ventral surface of quadrate ramus: form modest crests (0); prominent knobs (1).

**Character 303** (Turner, 2015: char. 303) Pterygoid flange shape: mediolaterally broad, reaching laterally beyond medial margin of quadrate condyles (0); relatively narrow, does not reach laterally to medial margin of quadrate condyles (1).

**Character 304** (Turner, 2015: char. 304) In ventral view, posterior process of maxilla relative to ITF excluded from ITF (0); forms part of ITF (1).

**Character 305** (Turner, 2015: char. 305) Highly modified ectopterygoid, mediolaterally broad and flattened with greatly expanded: absent (0); present, robust anterior process larger than posterior process (1); present, anterior and posterior process roughly equal in size (2).

**Character 306** (Turner, 2015: char. 306) In ventral view, palate medial to toothrow: forms a single continuous surface (0); ridge running on the palate medial to toothrow formed by maxilla and ectopterygoid (1).

**Character 307** (Turner, 2015: char. 307) Maxillary tooth row, penultimate and ultimate maxillary teeth enlarged and highly modified crushing tooth: absent (0); present (1).

**Character 308** (Turner, 2015: char. 308) Prefrontals: do not meet at midline (0); meet at midline (1).

**Character 309** (Turner, 2015: char. 309) Pear shaped external naris: absent (0); present (1).

**Character 310** (Turner, 2015: char. 310) Skull, dorsal surface at parietal-squamosal contact surface: continuous across suture (0); suture marked by groove or sulcus (1).

**Character 311** (Turner, 2015: char. 311) Maxilla, lateral surface, continuous groove or sulcus extending from orbital margin to near orbital margin towards narial opening: absent (0); present (1).

**Character 312** (Turner, 2015: char. 312) Maxilla, posteromedial process curving posteriorly onto palatine- formed nasopharyngeal passage: absent (0); present (1).

**Character 313** (Turner, 2015: char. 313) Squamosal, posterior half, dorsal and ventral rims of groove for external ear valve musculature: thin or parallel sided (0); flared posteriorly (1).

**Character 314** (Turner, 2015: char. 314) Lacrimal, in dorsal view, anterior extent on rostrum relative to prefrontal: prefrontal extends farther anteriorly (0); lacrimal extends farther anteriorly (1); lacrimal and prefrontal subequal in anterior extent (2).

**Character 315** (Turner, 2015: char. 315) Lacrimal, in dorsal view, mediolateral width relative to prefrontal: equal to or less than width of prefrontal (0); wider than prefrontal (1).

**Character 316** (Turner, 2015: char. 316) Premaxillae, degree of contact posterior to the incisive foramen: extensive contact (0); narrow contact (1).

**Character 317** (Turner, 2015: char. 317) Posterior margin of the palatines where they form the floor of the nasopharyngeal passage, shape: V- or U-shaped (0); straight (1).

**Character 318** (Turner, 2015: char. 318) Posterior margin of the choanal groove, location: anteriorly on the pterygoids (0); posteriorly on the pterygoids near the posterior margin of pterygoids (1).

**Character 319** (Turner, 2015: char. 319) Pterygoid-palatine contact, ventral aspect of palate, shape of the suture: transverse, or nearly so (0); prong of pterygoid projects anteriorly (1).

**Character 320** (Turner, 2015: char. 320) On palate, foramen located on premaxilla/maxilla suture near the alveolar border: absent (0); present (1).

**Character 321** (Turner, 2015: char. 321) Ectopterygoid/pterygoid contact in ventral view: complex, anterior part of ectopterygoid forming suture whereas posterior part of ectopterygoid overlaps the pterygoid (0); sutured along entire contact, no part of ectopterygoid overlapping pterygoid (1).

**Character 322** (Kuzmin et al., 2019) Length of the dentary symphysis, in dorsal view: shortened, not extending posteriorly beyond the level of the fifth dentary alveolus (0); elongated, extending posteriorly beyond the level of the fifth dentary alveolus (1).

**Character 323** (Kuzmin et al., 2019) Relative degree of the prefrontal-frontal participation to the medial orbital margin (in adults): equal, or the frontal forms most of the medial orbital margin (0); the prefrontal forms most of the medial orbital margin, with the frontal forming only small posteromedial corner (1).

**Character 324** (Kuzmin et al., 2019) Width of the premaxillary rostrum at the level of the reception of enlarged dentary teeth, in dorsal view (the character is scored only for taxa, which have dorsally or slightly anterodorsally oriented naris in the sagittal plane): narrow, width of the premaxillary rostrum is nearly equal to the width of the naris (0); wide, width of the premaxillary rostrum is nearly twice the width of the naris (1).

**2. Character and state descriptions for the matrix adapted from Pochat-Cottilloux et al. (2024) (Supplementary Dataset S2)**

**Character 1 (Turner, 2015: char. 1, modified from Clark, 1994: char. 1)**: External surface of dorsal cranial bones: smooth (0), slightly grooved (1), heavily ornamented with deep pits and grooves (2)

**Character 2 (Turner, 2015: char. 2, modified from Clark, 1994: char. 2)**: Skull expansion at orbits: gradual (0), abrupt (1).

**Character 3 (Turner, 2015: char. 3, modified from Clark, 1994: char. 3)**: Rostrum proportions: narrow oreinirostral (0), broad oreinirostral (1), nearly tubular (2), platyrostral (3).

**Character 4 (Clark, 1994: char. 4)**: Premaxilla participation in internarial bar: forming at least ventral half (0), little participation (1).

**Character 5 (Clark, 1994: char. 5)**: Premaxilla anterior to nares: narrow (0), broad (1).

**Character 6 (Turner, 2015: char. 6, modified from Clark, 1994: char. 6)**: External nares facing anterolaterally or anteriorly (0); dorsally, not separated by premaxillary bar from anterior edge of rostrum (1); dorsally, separated by premaxillary bar (2)

**Character 7 (Clark, 1994: char. 7)**: Palatal parts of premaxillae: do not meet posterior to incisive foramen (0), meet posteriorly along contact with maxillae (1).

**Character 8 (Clark, 1994: char. 8)**: Premaxilla-maxilla contact: premaxilla loosely overlies maxilla (i.e., posterodorsal process of premaxilla overlaps anterodorsal surface of maxilla) (0), or sutured together along butt joint (1).

**Character 9 (Turner, 2015: char. 9, modified from Clark, 1994: char. 9)**: Ventrally opened notch on ventral edge of rostrum at premaxilla-maxilla contact: absent (0), present as notch (1), present as large fenestra (2)

**Character 10 (Pol et al., 2009: char. 10, modified from Clark, 1994: char. 10)**: Posterior palatal branches of maxillae anterior to palatines: do not meet (0), meet extensively but posteriormost parts fail to meet (1), meet entirely (2)

**Character 11 (Clark, 1994: char. 11)**: Nasal contacts lacrimal (0), does not contact lacrimal (1).

**Character 12 (Clark, 1994: char. 12)**: Lacrimal contacts nasal along medial edge only (0), or along medial and anterior edges (1).

**Character 13 (Clark, 1994: char. 13)**: Nasal contribution to narial border: present (0), absent (1).

**Character 14 (Clark, 1994: char. 14)**: Nasal-premaxilla contact: present (0), absent (1).

**Character 15 (Turner, 2015: char. 15, modified from Clark, 1994: char. 15)**: Descending process of prefrontal: does not contact palate (0), contacts palate (1).

**Character 16 (Clark, 1994: char. 16)**: Postorbital-jugal contact: postorbital anterior to jugal (0), postorbital medial to jugal (1), or postorbital lateral to jugal (2).

**Character 17 (Clark, 1994: char. 17)**: Anterior part of jugal with respect to posterior part: as broad (0), twice as broad (1).

**Character 18 (Clark, 1994: char. 18)**: Jugal bar beneath infratemporal fenestra: dorsolaterally flattened (0), rod-shaped (1).

**Character 19 (Clark, 1994: char. 19)**: Quadratojugal dorsal process: narrow, contacting only small part of postorbital (0); broad, extensively contacting postorbital (1).

**Character 20 (Clark, 1994: char. 20)**: Frontal width between orbits: narrow, as broad as nasals (0); broad, twice as broad as nasals (1).

**Character 21 (Clark, 1994: char. 21)**: Frontals in mature specimens: paired (0), unpaired (1).

**Character 22 (Clark, 1994: char. 22)**: Dorsal surface of frontal and parietal: flat (0), with midline ridge (1).

**Character 23 (Buckley & Brochu, 1999: char. 81, modified from Clark, 1994: char. 23)**: Parieto-postorbital suture: absent from dorsal surface of skull roof and supratemporal fossa (0), absent from dorsal surface of skull roof but broadly present within supratemporal fossa (1), present within supratemporal fossa and on dorsal surface of skull roof (2).

**Character 24 (Clark, 1994: char. 24)**: Dorsal surface of supratemporal roof: complex (0); dorsally flat ‘skull table’ developed, with postorbital and squamosal bearing flat shelves extending laterally beyond quadrate contact (1).

**Character 25 (Turner, 2015: char. 25, modified from Clark, 1994: char. 25)**: Postorbital bar: sculpted (if skull sculpted) (0), unsculpted (1).

**Character 26 (Turner, 2015: char. 26, modified from Clark, 1994: char. 26)**: Postorbital bar: transversely flattened (0), cylindrical (1).

**Character 27 (Clark, 1994: char. 27)**: Vascular opening on dorsal surface of postorbital bar: absent (0), present (1).

**Character 28 (Turner, 2015: char. 28, modified from Clark, 1994: char. 28)**: Postorbital anterolateral process: absent or poorly developed (0); well developed, long, and acute (1).

**Character 29 (Clark, 1994: char. 29)**: Dorsal part of postorbital: with anterior and lateral edges only (0), with anterolaterally facing edge (1).

**Character 30 (Clark, 1994: char. 30)**: Dorsal end of postorbital bar broadens dorsally, continuous with dorsal part of postorbital (0); dorsal part of postorbital bar constricted, distinct from dorsal part of postorbital (1).

**Character 31 (Clark, 1994: char. 31)**: Bar between orbit and supratemporal fossa in mature specimens: broad and solid, with broadly sculpted dorsal surface if sculpture present (0); bar narrow, sculpting restricted to anterior surface (1).

**Character 32 (Turner, 2015: char. 32, modified from Clark, 1994: char. 32)**: Parietal: with broad occipital portion (0), without broad occipital portion (1).

**Character 33 (Clark, 1994: char. 33)**: Parietal: with broad sculpted region separating supratemporal fossae (0), with sagittal crest between supratemporal fossae (1).

**Character 34 (Clark, 1994: char. 34)**: Postparietal (dermosupraoccipital): a distinct element (0), not distinct (fused with parietal?) (1).

**Character 35 (Clark, 1994: char. 35)**: Posterodorsal corner of squamosal: squared off, lacking extra ‘lobe’ (0); with unsculptured ‘lobe’ (1).

**Character 36 (Turner, 2015: char. 36, modified from Clark, 1994: char. 36)**: Posterolateral process of squamosal: poorly developed and projecting horizontally at same level of skull (0); elongate, thin, and posteriorly directed, not ventrally deflected (1); elongate, posterolaterally directed, and ventrally deflected (2).

**Character 37 (Clark, 1994: char. 37)**: Palatines: do not meet on palate below narial passage (0); form palatal shelves that do not meet (1); meet ventral to narial passage, forming part of secondary palate (2).

**Character 38 (Clark, 1994: char. 38)**: Pterygoid: restricted to palate and suspensorium, joints with quadrate and basisphenoid overlapping (0); extends dorsally to contact laterosphenoid and form ventrolateral edge of trigeminal foramen, strongly sutured to quadrate and laterosphenoid (1).

**Character 39 (Turner, 2015: char. 39, modified from Clark, 1994: char. 39)**: Choanal opening: continuous with pterygoid ventral surface except for anterior and anterolateral borders (0), opens into palate through deep midline depression (choanal groove) (1).

**Character 40 (Clark, 1994: char. 40)**: Palatal surface of pterygoids: smooth (0), sculpted (1).

**Character 41 (Clark, 1994: char. 41)**: Pterygoids posterior to choanae: separated (0), fused (1).

**Character 42 (Ortega et al., 2000: char. 139, modified from Clark, 1994: char. 38)**: Depression on primary pterygoidean palate posterior to choana: absent or moderate in size, being narrower than palatine bar (0), wider than palatine bar (1).

**Character 43 (Clark, 1994: char. 43)**: Primary pterygoidean palate, role in forming choanal opening: does not enclose choana (0), completely encloses choana (1).

**Character 44 (Pol et al., 2009: char. 44, modified from Clark, 1994: char. 44)**: Anterior edge of choanae situated between suborbital fenestra (or anteriorly) (0), situated near the posterior edge of suborbital fenestra (1), situated near the posterior edge of pterygoid flange (2)

**Character 45 (Clark, 1994: char. 45)**: Quadrate: without fenestrae (0), with single fenestra (1), with three or more fenestrae on dorsal and posteromedial surfaces (2)

**Character 46 (Clark, 1994: char. 46)**: Posterior edge of quadrate: broad medial to tympanum, gently concave (0); or posterior edge narrow dorsal to otoccipital contact, strongly concave (1).

**Character 47 (Clark, 1994: char. 47)**: Dorsal, primary head of quadrate articulates with: squamosal, otoccipital, and prootic (0); prootic and laterosphenoid (1).

**Character 48 (Clark, 1994: char. 48)**: Ventrolateral contact of otoccipital with quadrate: very narrow (0), broad (1).

**Character 49 (Clark, 1994: char. 49)**: Quadrate, squamosal, and otoccipital: do not meet to enclose cranioquadrate passage (0), enclose passage near lateral edge of skull (1), meet broadly lateral to passage (2).

**Character 50 (Clark, 1994: char. 50)**: Pterygoid ramus of quadrate: with flat ventral edge (0), with deep groove along ventral edge (1).

**Character 51 (Clark, 1994: char. 51)**: Ventromedial part of quadrate: does not contact otoccipital (0), contacts otoccipital to enclose carotid artery and form passage for cranial nerves IX–XI (1).

**Character 52 (Clark, 1994: char. 52)**: Eustachian tubes: not enclosed between basioccipital and basisphenoid (0); entirely enclosed (1).

**Character 53 (Clark, 1994: char. 53)**: Basisphenoid rostrum (cultriform process): slender (0); dorsoventrally expanded (1).

**Character 54 (Clark, 1994: char. 54)**: Basipterygoid process: prominent, forming movable joint with pterygoid (0); small or absent, with basisphenoid joint suturally closed (1).

**Character 55 (Ortega et al., 2000: char. 68, modified from Clark, 1994: char. 55)**: Basisphenoid ventral surface: shorter than basioccipital (0); wide and similar to, or longer, in length than basioccipital (1).

**Character 56 (Clark, 1994: char. 56)**: Basisphenoid: exposed on ventral surface of braincase (0), virtually excluded from ventral surface by pterygoid and basioccipital (1).

**Character 57 (Clark, 1994: char. 57)**: Basioccipital: without well-developed bilateral tuberosities (0), with large pendulous tubera (1).

**Character 58 (Clark, 1994: char. 58)**: Otoccipital: without laterally concave descending flange ventral to subcapsular process (0); with flange (1).

**Character 59 (Clark, 1994: char. 59)**: Cranial nerves IX–XI: all pass through common large foramen vagi in otoccipital (0); cranial nerve IX passes medial to nerves X and XI in separate passage (1).

**Character 60 (Clark, 1994: char. 60)**: Otoccipital: without large ventrolateral part ventral to paroccipital process (0); with large ventrolateral part (1).

**Character 61 (Clark, 1994: char. 61)**: Crista interfenestralis between fenestrae pseudorotunda and ovalis: nearly vertical (0); horizontal (1).

**Character 62 (Clark, 1994: char. 62)**: Supraoccipital: forms dorsal edge of foramen magnum (0); otoccipitals broadly meet dorsal to foramen magnum, separating supraoccipital from foramen magnum (1).

**Character 63 (Clark, 1994: char. 63)**: Mastoid antrum: does not extend into supraoccipital (0); extends through transverse canal in supraoccipital to connect middle ear regions (1).

**Character 64 (Clark, 1994: char. 64)**: Posterior surface of supraoccipital: nearly flat (0); with bilateral posterior prominences (1).

**Character 65 (Turner, 2015: char. 65, modified from Clark, 1994: char. 65)**: Palpebrals: absent (0); one small palpebral present in orbit (1); one large palpebral (2); two large palpebrals (3)

**Character 66 (Clark, 1994: char. 66)**: External nares: divided by septum (0); confluent (1).

**Character 67 (Clark, 1994: char. 67)**: Antorbital fenestra: as large as orbit (0); about half the diameter of orbit (1); much smaller than the orbit (2); absent (3)

**Character 68 (Ortega et al., 2000: char. 41, modified from Clark, 1994: char. 68)**: Supratemporal fenestrae extension: relatively large, covering most of surface of skull roof (0); relatively short, fenestrae surrounded by flat and extended skull roof (1).

**Character 69 (Turner, 2015: char. 69, modified from Clark, 1994: char. 69)**: Choanal groove: undivided (0); partially septated (1); completely septated (2)

**Character 70 (Clark, 1994: char. 70)**: Dentary: extends posteriorly beneath mandibular fenestra (0); does not extend beneath fenestra (1).

**Character 71 (Turner, 2015: char. 71, modified from Clark, 1994: char. 71)**: Retroarticular process: absent or extremely reduced (0); very short, broad, and robust (1); with extensive, rounded, wide, and flat (or slightly concave) surface projecting posteroventrally and facing dorsomedially (2); posteriorly elongate, triangular, and facing dorsally (3); posteroventrally projecting and paddle-shaped (4).

**Character 72 (Clark, 1994: char. 72)**: Prearticular: present (0); absent (1).

**Character 73 (Turner, 2015: char. 73, modified from Clark, 1994: char. 73)**: Articular: without medial process (0); with short process not contacting braincase (1); with process articulating with otoccipital and basisphenoid (2)

**Character 74 (Clark, 1994: char. 74)**: Dorsal edge of surangular: flat (0); arched dorsally (1).

**Character 75 (Clark, 1994: char. 75)**: Mandibular fenestra: present (0); absent (1).

**Character 76 (Clark, 1994: char. 76)**: Insertion area for M. pterygoideous posterior: does not extend onto lateral surface of angular (0); extends onto lateral surface of angular (1).

**Character 77 (Turner, 2015: char. 77, modified from Clark, 1994: char. 77)**: Splenial involvement in mandibular symphysis in ventral view: not involved (0); involved slightly in symphysis (1); extensively involved (2)

**Character 78 (Turner & Sertich, 2010: char. 78, modified from Clark, 1994: char. 78)**: Posterior premaxillary teeth: similar in size to anterior teeth (0); longer but does not form an enlarged caniniform tooth (1); much longer forming one large premaxillary caniniform tooth (2); much longer forming two large premaxillary caniniform teeth (3)

**Character 79 (Turner, 2015: char. 79, modified from Clark, 1994: char. 79)**: Maxillary tooth waves: absent, no tooth size variation (0); one wave of teeth enlarged (1); enlarged maxillary teeth occur in two waves (festooned) (2)

**Character 80 (Clark, 1994: char. 80)**: Anterior dentary teeth opposite premaxilla-maxilla contact: no more than twice the length of other dentary teeth (0); more than twice the length (1).

**Character 81 (Turner, 2015: char. 81, modified from Clark, 1994: char. 81)**: Dentary teeth posterior to tooth opposite premaxilla-maxilla contact: equal in size (0); enlarged dentary teeth opposite to smaller teeth in maxillary toothrow (1).

**Character 82 (Ortega et al., 2000: char. 120, modified from Clark, 1994: char. 82)**: Anterior and posterior scapular edges: symmetrical in lateral view (0); anterior edge more strongly concave than posterior edge (1); dorsally narrow with straight edges (2).

**Character 83 (Ortega et al., 2000: char. 121, modified from Clark, 1994: char. 83)**: Coracoid length: up to two-thirds of scapular length (0); subequal in length to scapula (1).

**Character 84 (Clark, 1994: char. 84)**: Anterior process of ilium: similar in length to posterior process (0); one-quarter or less length of posterior process (1).

**Character 85 (Clark, 1994: char. 85)**: Pubis: rod-like without expanded distal end (0); with expanded distal end (1).

**Character 86 (Clark, 1994: char. 86)**: Pubis: forms anterior half of ventral edge of acetabulum (0); contacting ilium but partially excluded from acetabulum by anterior process of ischium (1); completely excluded from acetabulum by anterior process of ischium (2)

**Character 87 (Clark, 1994: char. 87)**: Distal end of femur: with large lateral facet for fibula (0); with very small facet (1).

**Character 88 (Clark, 1994: char. 88)**: Fifth pedal digit: with phalanges (0); without phalanges (1).

**Character 89 (Clark, 1994: char. 89)**: Atlas intercentrum: broader than long (0); as long as broad (1).

**Character 90 (Turner, 2015: char. 90, modified from Clark, 1994: char. 90)**: Cervical neural spines: all anteroposteriorly large (0); only posterior ones rod-like (1); all spines rod-like (2)

**Character 91 (Buscalioni & Sanz, 1988: char. 37 and Brochu, 1997: char. 7, modified from Clark, 1994: char. 91)**: Hypapophyses in cervicodorsal vertebrae: absent (0); present only in cervical vertebrae (1); present in cervical and first two dorsal vertebrae (2); present up to the third dorsal vertebra (3); present up to the fourth dorsal vertebrae (4)

**Character 92 (Clark, 1994: char. 92)**: Cervical vertebrae: amphicoelous or amphyplatyan (0); procoelous (1).

**Character 93 (Clark, 1994: char. 93)**: Trunk vertebrae: amphicoelous or amphyplatyan (0); procoelous (1).

**Character 94 (Clark, 1994: char. 94)**: All caudal vertebrae: amphicoelous or amphyplatyan (0); first caudal vertebra biconvex with other procoelous (1); procoelous (2).

**Character 95 (Turner, 2015: char. 95, modified from Clark, 1994: char. 95)**: Dorsal osteoderms, shape: rounded or ovate (0); rectangular, broader than long (1); square (2); rectangular, longer than broad (3).

**Character 96 (Turner, 2015: char. 96, modified from Clark, 1994: char. 96 and Brochu, 1997: char. 40)**: Dorsal osteoderms: without articular anterior process (0); with a discrete convexity on anterior margin (1); with a well-developed process located anterolaterally in dorsal parasagittal osteoderms (2)

**Character 97 (Ortega et al., 2000: char. 107 & 108, modified from Clark, 1994: char. 97)**: Rows of dorsal primary osteoderms (sensu Frey, 1988): two parallel rows (0); more than two rows (1); more than four rows (2)

**Character 98 (Clark, 1994: char. 98)**: Osteoderms: some or all imbricated (0); sutured to one another (1).

**Character 99 (Clark, 1994: char. 99)**: Tail osteoderms: dorsal only (0); surrounding tail (1).

**Character 100 (Clark, 1994: char. 100)**: Trunk osteoderms: absent from ventral part of the trunk (0); present on ventral part (1).

**Character 101 (Clark, 1994: char. 101)**: Osteoderms: with longitudinal keels on dorsal surfaces (0); without longitudinal keels (1).

**Character 102 (Wu & Sues, 1996: char. 14)**: Jugal: participating in margin of antorbital fossa (0); separated from it (1).

**Character 103 (Turner, 2015: char. 103, modified from Wu & Sues, 1996: char. 17)**: Mandibular symphysis in lateral view: shallow and tapering anteriorly (0); deep and tapering anteriorly (1); deep and anteriorly convex (2); shallow and anteriorly convex (3).

**Character 104 (Turner, 2015: char. 104, modified from Wu & Sues, 1996: char. 23)**: Articular facet for quadrate condyle: equal in length to quadrate condyles (0); slightly longer (1); close to three times length of quadrate condyles (2)

**Character 105 (Turner, 2015: char. 105, modified from Wu & Sues, 1996: char. 24 and Wu et al., 1997: char. 124)**: Jaw joint: placed level with occipital condyle (0); below occipital condyle but above level of lower toothrow (1); below level of toothrow (2)

**Character 106 (Turner, 2015: char. 106, modified from Wu & Sues, 1996: char. 27 and Ortega et al., 2000: char. 133)**: Premaxillary tooth number: six (0); five (1); four (2); three (3); two (4)

**Character 107 (Turner, 2015: char. 107, modified from Wu & Sues, 1996: char. 29)**: Unsculptured region along alveolar margin on lateral surface of maxilla: absent (0); present (1).

**Character 108 (Wu & Sues, 1996: char. 30)**: Maxillary tooth number: eight or more (0); seven (1); six (2); five (3); four (4)

**Character 109 (Wu & Sues, 1996: char. 33)**: Coracoid, posteromedial or ventromedial process absent (0); elongate posteromedial process present (1); distally expanded ventromedial process present (2).

**Character 110 (Wu & Sues, 1996: char. 40)**: Radiale and ulnare, size: short and massive (0); elongate (1).

**Character 111 (Turner, 2015: char. 111, modified from Gomani, 1997: char. 4)**: Prefrontals anterior to orbits: elongate, oriented parallel to anteroposterior axis of skull (0); short and broad, oriented posteromedially-anterolaterally (1).

**Character 112 (Turner, 2015: char. 112, modified from Gomani, 1997: char. 32)**: Basioccipital and ventral part of otoccipital, orientation: facing posteriorly (0); posteroventrally (1).

**Character 113 (Buscalioni & Sanz, 1988: char. 35)**: Vertebral centra, shape: cylindrical (0); spool-shaped (1).

**Character 114 (Turner, 2015: char. 114, modified from Buscalioni & Sanz, 1988: char. 39)**: Transverse process of posterior dorsal vertebrae, shape: dorsoventrally low and laminar (0); dorsoventrally high (1).

**Character 115 (Buscalioni & Sanz, 1988: char. 44)**: Number of sacral vertebrae: two (0); more than two (1).

**Character 116 (Buscalioni & Sanz, 1988: char. 49)**: Supraacetabular crest: present (0); absent (1).

**Character 117 (Buscalioni & Sanz, 1988: char. 54)**: Proximal end of radiale, shape: expanded symmetrically, similarly to distal end (0); more expanded proximolaterally than proximomedially (1).

**Character 118 (Ortega et al., 1996: char. 5):** Lateral surface of anterior region of surangular and posterior region of dentary: without longitudinal depression (0); with longitudinal depression (1).

**Character 119 (Ortega et al., 1996: char. 9)**: Ventral exposure of splenials: absent (0); present (1).

**Character 120 (Turner, 2015: char. 120, modified from Andrade & Bertini, 2008: char. 120; Ortega et al., 1996: char. 11 and Ortega et al., 2000: char. 100)**: Tooth margin carinae: without carinae or with smooth or crenulated carinae (0); with homogeneous denticulate carinae (denticles are small and symmetrical in form as in ziphodont teeth) (1); with heterogeneous carinae possessing rounded tubercle-like denticles, developed preferentially along posterior margin (2).

**Character 121 (Turner, 2015: char. 121, modified from Pol, 1999a: char. 133 and Ortega et al., 2000: char. 145)**: Lateral surface of anterior process of jugal: flat or convex (0); with broad shelf below the orbit with triangular depression underneath it (1).

**Character 122 (Pol, 1999a: char. 134)**: Jugal, extension below the orbit: does not exceed the anterior margin of orbit (0); exceeds margin of orbit (1).

**Character 123 (Pol, 1999a: char. 135)**: Notch in premaxilla on lateral edge of external nares: absent (0); present on the dorsal half of the external nares lateral margin (1).

**Character 124 (Pol, 1999a: char. 136)**: Dorsal border of external nares: formed mostly by the nasals (0); formed by both the nasals and premaxilla (1).

**Character 125 (Pol, 1999a: char. 138)**: Posterodorsal process of premaxilla: absent (0); present extending posteriorly and wedging between maxilla and nasal (1).

**Character 126 (Pol, 1999a: char. 139 and Ortega et al., 2000: char. 9)**: Premaxilla-maxilla suture in palatal view, medial to alveolar region, orientation of suture: anteromedially directed (0); sinusoidal, posteromedially directed on its lateral half and anteromedially directed along its medial region (1); posteromedially directed (2); straight (3); posteromedially curved (U-shaped) (4).

**Character 127 (Pol, 1999a: char. 140)**: Nasal lateral border posterior to external nares: laterally concave (0); straight (1).

**Character 128 (Pol, 1999a: char. 141)**: Nasal lateral edges: nearly parallel (0); oblique to each other, converging anteriorly (1); oblique to each other, diverging anteriorly (2).

**Character 129 (Pol, 1999a: char. 143)**: Palatine anteromedial margin: exceeding anterior margin of suborbital fenestrae: extending anteriorly between maxillae (0); not exceeding anterior margin of suborbital fenestrae (1).

**Character 130 (Pol, 1999a: char. 144)**: Dorsoventral height of jugal antorbital region with respect to infraorbital region equal or lower (0); antorbital region more expanded than infraorbital region (1).

**Character 131 (Pol, 1999a: char. 145)**: Maxilla-lacrimal contact in antorbital fossa: partially included (0); completely included (1).

**Character 132 (Pol, 1999a: char. 146)**: Lateral eustachian tube openings, location: located posteriorly to medial opening (0); aligned anteroposteriorly and dorsoventrally (1).

**Character 133 (Pol, 1999a: char. 147)**: Anterior process of ectopterygoid: developed (0); reduced or absent (1).

**Character 134 (Pol, 1999a: char. 148)**: Posterior process of ectopterygoid: developed (0); reduced or absent (1).

**Character 135 (Pol, 1999a: char. 149 and Ortega et al., 2000: char. 13)**: Small foramen located in the lateral surface of premaxilla-maxilla suture in lateral surface (not for big mandibular teeth): absent (0); present (1).

**Character 136** **(Pol, 1999a: char. 150)**: Jugal posterior process, extent of process: exceeding posteriorly the infratemporal fenestrae (0); does not exceed infratemporal fenestrae (1).

**Character 137 (Pol, 1999a: char. 151)**: Compressed crown of maxillary teeth, orientation: oriented parallel to the longitudinal axis of skull (0); obliquely disposed (1).

**Character 138 (Pol, 1999a: char. 152)**: Large and aligned neurovascular foramina on lateral maxillary surface: absent (0); present (1).

**Character 139 (Turner, 2015: char. 139, modified from Pol, 1999a: char. 153)**: External surface of maxilla and premaxilla, general shape: with single plane facing laterally (0); with ventral region facing laterally and dorsal region facing dorsolaterally (1).

**Character 140 (Turner, 2015: char. 140, modified from Pol, 1999a: char. 154; Ortega et al., 2000: char. 104 and Andrade & Bertini, 2008: char. 135)**: Maxillary teeth, lateral compression: absent (0); present, compression asymmetrically occurring only along distal margin giving teeth a teardrop shape (1); present, lateral compression symmetrically developed (2).

**Character 141 (Pol, 1999a: char. 155)**: Posteroventral corner of quadratojugal: reaching quadrate condyles (0); does not reach quadrate condyles (1).

**Character 142 (Turner, 2015: char. 142, modified from Pol, 1999a: char. 156)**: Base of postorbital process of jugal, orientation: directed posterodorsally (0); directed dorsally (1); directed anterodorsally (2)

**Character 143 (Pol, 1999a: char. 157)**: Postorbital process of jugal, location on jugal: anteriorly placed (0); in middle (1); posteriorly positioned (2)

**Character 144 (Pol, 1999a: char. 158 and Ortega et al., 2000: char. 36)**: Postorbital-ectopterygoid contact: present (0); absent (1).

**Character 145 (Pol, 1999a: char. 161)**: Quadratojugal, ornamentation: absent (0); ornamented at the base (1).

**Character 146 (Pol, 1999a: char. 162)**: Prefrontal-maxilla contact in the inner anteromedial region of orbit absent (0); present (1).

**Character 147 (Pol, 1999a: char. 163)**: Basisphenoid, exposure on braincase: without lateral exposure (0); with lateral exposure (1).

**Character 148 (Pol, 1999a: char. 165)**: Quadrate process of pterygoids: well developed (0); poorly developed (1).

**Character 149 (Turner, 2015: char. 149, modified from Pol, 1999a: char. 166 and Ortega et al., 2000: char. 44)**: Quadrate major axis, direction of orientation: posteroventrally (0); ventrally (1); anteroventrally (2)

**Character 150 (Pol, 1999a: char. 167)**: Quadrate distal end: with only one plane facing posteriorly (0); with two distinct faces in posterior view, a posterior one and a medial one bearing foramen aereum (1).

**Character 151 (Pol, 1999a: char. 168)**: Anteroposterior development of neural spine in axis: well developed, covering all of neural arch length (0); poorly developed, located over posterior half of neural arch (1).

**Character 152 (Pol, 1999a: char. 169)**: Prezygapophyses of axis, development relative to neural arch: not exceeding edge of neural arch (0); exceeding the anterior margin of neural arch (1).

**Character 153 (Pol, 1999a: char. 170)**: Postzygapophyses of axis: well developed, curved laterally (0); poorly developed (1).

**Character 154 (Turner, 2015: char. 154, modified from Pol, 1999b: char. 212)**: Shape of dentary symphysis in ventral view: tapering anteriorly forming an angle (0); U-shaped, smoothly curving anteriorly (1); lateral edges longitudinally oriented, convex anterolateral corner and extensive transversally oriented anterior edge (2)

**Character 155 (Pol, 1999b: char. 213)**: Unsculpted region in the dentary below the tooth row absent (0); present (1).

**Character 156 (Buckley & Brochu, 1999: char. 102)**: Surangular, contribution to the glenoid fossa: forms only the lateral wall of glenoid (0); forms approximately one-third of the glenoid (1).

**Character 157 (Turner, 2015: char. 157, modified from Buckley & Brochu, 1999: char. 102)**: Femur, anterior margin: linear (0); bears flange for PIFI 1 musculature (1).

**Character 158 (Turner, 2015: char. 158, modified from Buckley & Brochu, 1999: char. 105)**: Dentary, lateral surface: smooth lateral to seventh alveolus (0); with lateral concavity for the reception of the enlarged maxillary tooth (1).

**Character 159 (Turner, 2015: char. 159, modified from Ortega et al., 1996: char. 1 and Buckley & Brochu, 1999: char. 107)**: Dorsal edge of dentary: slightly concave or straight and subparallel to the longitudinal axis of skull (0); straight with an abrupt dorsal expansion, being straight posteriorly (1); with a single dorsal expansion and concave posterior to this (2); sinusoidal, with two concave waves (3)

**Character 160 (Turner, 2015: char. 160, modified from Ortega et al., 1996: char. 2 and Buckley & Brochu, 1999: char. 108)**: Dentary compression and ventrolateral surface anterior to mandibular fenestra: compressed and vertical (0); not compressed and convex (1).

**Character 161 (Ortega et al., 1996: char. 7 and Buckley & Brochu, 1999: char. 110)**: Splenial posterior to symphysis: thin (0); robust dorsally (1).

**Character 162 (Ortega et al., 1996: char. 13 and Buckley et al., 2000: char. 117)**: Cheek teeth: not constricted at base of crown (0); constricted (1).

**Character 163 (Ortega et al., 2000: char. 10)**: Ventral edge of premaxilla, location relative to maxilla: at the same height as the ventral edge of maxilla (0); located deeper, with the dorsal contour of anterior part of dentary strongly concave (1).

**Character 164 (Turner, 2015: char. 164, modified from Ortega et al., 2000: char. 19)**: Maxillary dental implantation: teeth in isolated alveoli (0); located in dental groove (1).

**Character 165 (Ortega et al., 2000: char. 24)**: Caudal tip of nasals: converge at sagittal plane (0); separated by anterior sagittal projection of frontals (1).

**Character 166 (Ortega et al., 2000: char. 33)**: Relative length between squamosal and postorbital: squamosal is longer (0); postorbital longer (1).

**Character 167 (Turner, 2015: char. 167, modified from Ortega et al., 2000: char. 34)**: Jugal portion of postorbital bar, relative to lateral surface of jugal: flush with lateral surface (0); anteriorly continuous but posteriorly inset (1); medially displaced and a ridge separate postorbital bar from lateral surface of jugal (2)

**Character 168 (Ortega et al., 2000: char. 42)**: Outer surface of squamosal laterodorsally oriented: extensive (0); reduced and sculpted (1); reduced and unsculpted (2).

**Character 169 (Ortega et al., 2000: char. 47)**: Quadratojugal spine at posterior margin of infratemporal fenestra: absent (0); present (1).

**Character 170 (Turner, 2015: char. 170, modified from Ortega et al., 2000: char. 53)**: Quadrate condyles: poorly developed intercondylar groove (0); medial condyle expands ventrally, being separated from lateral condyle by a deep intercondylar groove (1).

**Character 171 (Ortega et al., 2000: char. 62)**: Exposure of supraoccipital in skull roof: absent (0); present (1).

**Character 172 (Ortega et al., 2000: char. 70)**: Nasal participation in antorbital fenestra: present (0); absent (1).

**Character 173 (Ortega et al., 2000: char. 75)**: Anterior opening of temporo-orbital canal in dorsal view: exposed (0); hidden and overlapped by squamosal rim of supratemporal fossa (1).

**Character 174 (Ortega et al., 2000: char. 90)**: Foramen intramandibularis oralis: small or absent (0); big and slotlike (1).

**Character 175 (Turner, 2015: char. 175, modified from Ortega et al., 2000: char. 98)**: Coronoid size: short and located below the dorsal edge of the mandibular ramus (0); anteriorly extended with posterior region elevated at dorsal margin of mandibular ramus (1).

**Character 176 (Ortega et al., 2000: char. 101)**: Width of root of teeth with respect to crown: narrower or equal (0); wider (1).

**Character 177 (Ortega et al., 2000: char. 109)**: Gap in cervico-thoracic dorsal armor: absent (0); present (1).

**Character 178 (Ortega et al., 2000: char. 130)**: Lateral contour of snout in dorsal view: straight (0); sinusoidal (1).

**Character 179 (Ortega et al., 2000: char. 138)**: Pterygoidean flanges: laminar and expanded (0); bar-like and elongate (1); bar-like and poorly developed (2).

**Character 180 (Ortega et al., 2000: char. 146)**: Ectopterygoid medial process, shape: single (0); forked (1).

**Character 181 (Turner, 2015: char. 181, modified from Ortega et al., 2000: char. 157)**: Skull roof, shape in dorsal view: rectangular (0); trapezoidal (1).

**Character 182 (Ortega et al., 2000: char. 30)**: Prefrontal pillars when integrated in palate: pillars transversely expanded (0); transversely expanded in their dorsal part and columnar ventrally (1); longitudinally expanded in their dorsal part and columnar ventrally (2).

**Character 183 (Ortega et al., 2000: char. 21)**: Ventral edge of maxilla in lateral view: straight or convex (0); sinusoidal (1).

**Character 184 (Turner, 2015: char. 184, modified from Ortega et al., 2000: char. 156)**: Position of first enlarged maxillary teeth: second or third alveolus (0); fourth or fifth (1).

**Character 185 (Pol & Apesteguía, 2005: char. 180)**: Splenial-dentary suture at symphysis on ventral surface: v-shaped (0); transversal (1).

**Character 186 (Pol & Apesteguía, 2005: char. 181)**: Posterior peg at symphysis: absent (0); present (1).

**Character 187 (Pol & Apesteguía, 2005: char. 182)**: Posterior ridge on glenoid fossa of articular: present (0); absent (1).

**Character 188 (Turner, 2015: char. 188, modified from Pol et al., 2009: char. 188; Gomani, 1997: char. 46; Buckley et al., 2000: char. 113 & Andrade & Bertini, 2008: char. 149)**: Cusps of teeth, number and conformation: one unique cusp (0); one main cusp with smaller cusps arranged in one row (1); one main cusp with smaller cusps arranged in more than one row (2); several cusps of equal size arranged in more than one row (3)

**Character 189 (Pol & Apesteguía, 2005: char. 184)**: Dorsal surface of mandibular symphysis: flat or slightly concave (0); strongly concave and narrow, trough-shaped (1).

**Character 190 (Pol & Apesteguía, 2005: char. 185)**: Medial surface of splenials posterior to symphysis: flat or slightly convex (0); markedly concave (1).

**Character 191 (Pol & Apesteguía, 2005: char. 186)**: Choanal septum shape: narrow vertical bony sheet (0); T-shaped bar expanded ventrally (1); massive and blocky (2).

**Character 192 (Pol & Norell, 2004a: char. 164)**: Cross-section of distal end of quadrate: mediolaterally wide and anteroposteriorly thin (0); subquadrangular (1).

**Character 193 (Pol & Apesteguía, 2005: char. 188)**: Lateral surface of dentaries below alveolar margin, at mid to posterior region of tooth row: vertically oriented, continuous with rest of lateral surface of dentaries (0); flat surface exposed dorsolaterally, divided by ridge from rest of lateral surface of dentaries (1).

**Character 194 (Pol & Norell, 2004a: char. 165)**: Palatine-pterygoid contact on palate: palatines overlie pterygoids (0); palatines firmly sutured to pterygoids (1).

**Character 195 (Pol et al., 2004: char. 164)**: Ectopterygoid main axis orientation: laterally or slightly anterolaterally (0); anteriorly, subparallel to the skull longitudinal axis (1).

**Character 196 (Wu et al., 1997: char. 103)**: Squamosal descending process: absent (0); present (1).

**Character 197 (Turner, 2015: char. 197, modified from Wu et al., 1997: char. 105)**: Development of distal quadrate body ventral to otoccipital-quadrate contact: distinct (0); incipiently distinct (1); indistinct (2)

**Character 198 (Wu et al., 1997: char. 106)**: Pterygoid flanges: thin and laminar (0); dorsoventrally thick, with pneumatic spaces (1).

**Character 199 (Wu et al., 1997: char. 108)**: Postorbital participation in infratemporal fenestra: almost or entirely excluded (0); bordering infratemporal fenestra (1).

**Character 200 (Wu et al., 1997: char. 109)**: Palatines, contribution to suborbital fenestra: form margin of suborbital fenestra (0); excluded from margin of suborbital fenestra (1).

**Character 201 (Wu et al., 1997: char. 110)**: Angular posterior to mandibular fenestra, location on mandible: widely exposed on lateral surface of mandible (0); shifted to ventral surface of mandible (1).

**Character 202 (Wu et al., 1997: char. 112)**: Posteroventral edge of mandibular ramus, shape: straight or convex (0); markedly deflected (1).

**Character 203 (Pol et al., 2009: char. 203, modified from Wu et al., 1997: char. 128)**: Quadrate ramus of pterygoid, width in ventral view: narrow (0); broad (1).

**Character 204 (Wu et al., 1997: char. 121)**: Pterygoids, contact on palate: not in contact anterior to basisphenoid on palate (0); pterygoids in contact (1).

**Character 205 (Turner, 2015: char. 205, modified from Wu et al., 1997: char. 122)**: Olecranon: well developed (0); absent (1).

**Character 206 (Wu et al., 1997: char. 123)**: Cranial table width with respect to ventral portion of skull: as wide as ventral portion (0); narrower than the ventral portion of the skull (1).

**Character 207 (Wu et al., 1997: char. 127)**: Depression on posterolateral surface of maxilla: absent (0); present (1).

**Character 208 (Pol et al., 2009: char. 208, modified from Wu et al., 1997: char. 128)**: Paired anterior palatal fenestrae: absent (0); present (1).

**Character 209 (Pol & Norell, 2004a: char. 179)**: Paired ridges located medially on ventral surface of basisphenoid: absent (0); present (1).

**Character 210 (Pol et al., 2004: char. 179)**: Ventral margin of infratemporal bar of jugal: straight (0); dorsally arched (1).

**Character 211 (Pol & Norell, 2004a: char. 180)**: Posterolateral end of quadratojugal, shape and relationship with quadrate: acute or rounded, tightly overlapping the quadrate (0); with sinusoidal ventral edge and wide and rounded posterior edge slightly overhanging the lateral surface of the quadrate (1).

**Character 212 (Pol & Norell, 2004a: char. 181)**: Quadrate body distal to otoccipital-quadrate, orientation of contact in posterior view: ventrally oriented (0); ventrolaterally oriented (1).

**Character 213 (Gasparini et al., 1993: char. 3)**: Wedge-like process of maxilla in lateral surface of premaxilla-maxilla suture: absent (0); present (1).

**Character 214 (Pol & Norell, 2004b: char. 181)**: Palpebrals: separated from (or weakly sutured to) lateral edge of frontals (0); extensively sutured to each other and to lateral margin of frontals (1).

**Character 215 (Pol & Norell, 2004b: char. 182)**: External surface of ascending process of jugal: exposed laterally (0); exposed posterolaterally (1).

**Character 216 (Turner & Sertich, 2010: char. 216, modified from Pol & Norell, 2004b: char. 183)**: Longitudinal ridge on lateral surface of jugal below infratemporal fenestra: absent (0); present, running entire length of posterior process of jugal (1); present, running entire length of jugal (2)

**Character 217 (Pol & Norell, 2004b: char. 184)**: Dorsal surface of posterolateral region of squamosal: without ridges (0); with three curved ridges oriented longitudinally (1).

**Character 218 (Pol & Norell, 2004b: char. 185)**: Ridge along dorsal section of quadrate-quadratojugal contact: absent (0); present (1).

**Character 219 (Pol et al., 2009: char. 219, modified from Pol & Norell, 2004b: char. 186)**: Sharp ridge on the surface of the angular: absent (0); present on the ventral-most margin (1); present along the lateral surface (2).

**Character 220 (Pol & Norell, 2004b: char. 187)**: Longitudinal ridge along the dorsolateral surface of surangular: absent (0); present (1).

**Character 221 (Pol & Norell, 2004b: char. 188)**: Dorsal surface of osteoderms ornamented with anterolaterally and anteromedially directed ridges (fleur de lys pattern of Osmólska et al., 1997): absent (0); or present (1).

**Character 222 (Pol & Norell, 2004b: char. 189)**: Cervical region surrounded by lateral and ventral osteoderms sutured to dorsal elements: absent (0); present (1).

**Character 223 (Pol & Norell, 2004b: char. 190)**: Appendicular osteoderms: absent (0); present (1).

**Character 224 (Ortega et al., 2000: char. 72)**: Supratemporal fenestra: present (0); absent (1).

**Character 225 (Pol et al., 2009: char. 225, modified from Pol & Apesteguía, 2005: char. 221)**: Flat ventral surface of internal nares septum: parallel sided (0); tapering anteriorly (1); expanding anteriorly (2).

**Character 226 (Turner, 2015: char. 226, modified from Pol & Apesteguía, 2005: char. 221)**: Perinarial fossa: restricted extension (0); extensive, with distinctly concave surface facing anteriorly (1); large concave surface facing anteriorly, projecting anteroventrally from external nares and opening toward alveolar margin (2); extremely large and well-developed, occupying nearly entire surface of premaxilla ventral to external naris (3)

**Character 227 (Turner, 2015: char. 227, modified from Sereno et al., 2001: char. 67)**: Premaxillary palate, circular paramedian depressions: absent (0); present, located anteriorly on premaxilla (1); present, located at premaxilla-maxilla suture (2).

**Character 228 (Pol & Apesteguía, 2005: char. 223)**: Nasals, shape of posterolateral region: flat surface facing dorsally (0); lateral region deflected ventrally, forming part of the lateral surface of snout (1).

**Character 229 (Zaher et al., 2006: char. 193)**: Lacrimal, posterior extent and relationship with jugal: lacrimal: extending posteroventrally, widely contacting the jugal (0); tapers ventroposteriorly, does not contact or contacts jugal only slightly (1).

**Character 230 (Zaher et al., 2006: char. 194)**: Jugal, large foramen on the lateral surface, near its anterior margin: absent (0); present (1).

**Character 231 (Turner, 2015: char. 231, modified from Zaher et al., 2006: char. 195)**: Procumbent premaxillary alveoli: absent (0); present (1).

**Character 232 (Zaher et al., 2006: char. 196 & Turner, 2004: char. 119, modified from Martinelli, 2003: char. 36)**: Palatines, orientation: run parasagittally along midline (0); diverge laterally becoming rod-like caudally forming palatine bars (1).

**Character 233 (Zaher et al., 2006: char. 197)**: Ectopterygoid, participation in the palatine bar: absent (0); present (1).

**Character 234 (Pol & Norell, 2004a: char. 183)**: Choanal opening: opened posteriorly and continuous with pterygoid surface (0); closed posteriorly by an elevated wall formed by the pterygoids (1).

**Character 235 (Zaher et al., 2006: char. 198)**: Ectopterygoid, extent of medial projection on the ventral surface of pterygoid flanges: barely extended (0); widely extended, covering approximately the lateral half of the ventral surface of the pterygoid flanges (1).

**Character 236 (Gasparini et al., 2006: char. 236)**: Evaginated maxillary alveolar edges: absent (0), present as continuous sheet (1); present as discrete evaginations at each alveolus (2).

**Character 237 (Gasparini et al., 2006: char. 237)**: Premaxilla, foramen in perinarial depression: absent (0); present (1).

**Character 238 (Sereno et al., 2001: char. 27)**: Frontal, anterior ramus with respect to anterior tip of the prefrontal: ending posteriorly (0); ending anteriorly (1).

**Character 239 (Turner, 2015: char. 239, modified from Sereno et al., 2001: char. 68)**: Premaxilla, anterior alveolar margin orientation: vertical (0); inturned (1).

**Character 240 (Turner, 2015: char. 240, modified from Sereno et al., 2001: char. 69)**: Premaxillary tooth row orientation: arched posteriorly from midline (0); angled posterolaterally, at 120◦ angle (1); transverse (2)

**Character 241 (Sereno et al., 2001: char. 70)**: Last premaxillary tooth position relative to tooth row: anterior (0); anterolateral (1).

**Character 242 (Gasparini et al., 2006: char. 242)**: Posterior teeth with rings of undulated enamel: absent (0); present (1).

**Character 243 (Gasparini et al., 2006: char. 243, modified from Brochu, 1999: char. 108)**: Maxilla-palatine suture, shape of palatines: palatine anteriorly rounded (0); palatine anteriorly pointed (1); palatine invaginated (2).

**Character 244 (Gasparini et al., 2006: char. 244)**: Postorbital bar, lateral surface formed by: postorbital and jugal (0); only by postorbital (1).

**Character 245 (Gasparini et al., 2006: char. 245)**: Surangular groove, enlarged foramen at anterior end: absent (0); present (1).

**Character 246 (Gasparini et al., 2006: char. 246)**: Shape of antorbital fossa: subcircular or subtriangular (0); elongate, low, and oriented obliquely (1).

**Character 247 (Gasparini et al., 2006: char. 247)**: Prefrontal lateral development: reduced (0); enlarged, extending laterally over the orbit (1).

**Character 248 (Gasparini et al., 2006: char. 248)**: Foramen for the internal carotid artery: reduced, similar in size to openings for cranial nerves IX–XI (0); extremely enlarged (1).

**Character 249 (Gasparini et al., 2006: char. 249)**: Squamosal posterolateral region, lateral to paroccipital process: narrow (0); bearing subrounded flat surface (1).

**Character 250 (Gasparini et al., 2006: char. 250)**: Posteromedial branch of squamosal, orientation: transversely oriented (0); posterolaterally oriented (1).

**Character 251 (Gasparini et al., 2006: char. 251)**: Squamosal, dorsal margin of occipital flange: straight (0); dorsally concave (1).

**Character 252 (Gasparini et al., 2006: char. 252)**: Sculpture in external surface of rostrum: absent (0); present (1).

**Character 253 (Gasparini et al., 2006: char. 253)**: Longitudinal depressions on palatal surface of maxillae and palatines: absent (0); present (1).

**Character 254 (Gasparini et al., 2006: char. 254)**: Angle between medial and anterior margins of supratemporal fossa: approximately 90 degrees (0); approximately 45 degrees (1).

**Character 255 (Gasparini et al., 2006: char. 255)**: Sacral vertebrate, direction of transverse process: laterally (0); markedly deflected ventrally (1).

**Character 256 (Gasparini et al., 2006: char. 256)**: Prefrontal and lacrimal around orbits: forming flat rims (0); evaginated, forming elevated rims (1).

**Character 257 (Gasparini et al., 2006: char. 257)**: Nasal bones: paired (0); partially or completely fused (1).

**Character 258 (Brochu, 1997: char. 3)**: Axial neural spines, width of posterior half: wide (0); narrow (1).

**Character 259 (Brochu, 1997: char. 19)**: Axis hypapophysis, deep fork: present (0); absent (1).

**Character 260 (Brochu, 1997: char. 27)**: Ulna, width of olecranon process: narrow and subangular (0); wide and rounded (1).

**Character 261 (Brochu, 1997: char. 29)**: M. teres major and M. dorsalis scapulae: insert separately on humerus, scars can be distinguished dorsal to deltopectoral crest (0); insert with common tendon, single insertion scar (1).

**Character 262 (Turner, 2015: char. 262, modified from Brochu, 1997: char. 53)**: Dentary, projection of anterior alveoli: project anterodorsally (0); weakly procumbent (1); strongly procumbent (2)

**Character 263 (Brochu, 1997: char. 84)**: Squamosal, dorsal and ventral rims of squamosal groove for external ear valve musculature: parallel (0); squamosal groove flares anteriorly (1).

**Character 264 (Brochu, 1997: char. 91)**: Ectopterygoid, contact with maxilla near toothrow: ectopterygoid abuts maxillary toothrow (0); maxilla broadly separates ectopterygoid from maxillary toothrow (1).

**Character 265 (Brochu, 1997: char. 92)**: Shallow fossa at anteromedial corner of supratemporal fenestra: present (0); absent, anteromedial corner of supratemporal fenestra smooth (1).

**Character 266 (Pol et al., 2009 : char. 266, modified from Brochu, 1997: char. 103)**: Lateral margins of frontal, relative to the skull surface: flush with skull surface (0); elevated, forming ridged orbital margins (1).

**Character 267 (Brochu, 1997: char. 130)**: Laterosphenoid, orientation of capitate process: laterally oriented (0); anteroposteriorly oriented toward midline (1).

**Character 268 (Pol et al., 2009 : char. 268, modified from Brochu, 1997: char. 141)**: Otoccipital, development of boss and paroccipital process: process lateral to cranioquadrate opening short (0); boss small or absent on paroccipital process, process lateral to cranioquadrate opening long (1).

**Character 269 (Brochu, 1997: char. 149, modified from Norell, 1988: char. 32)**: Ectopterygoid, extent along lateral pterygoid flange, at maturity: extends to posterior tip of lateral pterygoid flange (0); does not extend to posterior tip of lateral pterygoid (1).

**Character 270 (Turner, 2015: char. 270, modified from Brochu, 1997: char. 153)**: Incisive foramen, location relative to premaxillary toothrow: foramen situated far from premaxillary toothrow, at the level of the second or third alveolus (0); abuts premaxillary toothrow (1); projects between first premaxillary teeth (2).

**Character 271 (Pritchard et al., 2013: char. 271, modified from Turner, 2006: char. 126)**: Ventral surface of choanal septum: smooth to slightly depressed (0); marked by an acute groove (1); vomeral septum divided into bilateral laminae (2).

**Character 272 (Pol et al., 2009: char. 272, modified from Turner, 2006: char. 128)**: Proximal-most portion of fibular head: straight sided to weakly developed posteriorly (0); very sharply projecting posteriorly, forming distinct extension (1).

**Character 273 (Turner, 2015: char. 273, modified from Turner, 2006: char. 129)**: Cervical rib shaft, posterior process, posterodorsally projecting spine at the junction with tubercular process: absent (0); present (1).

**Character 274 (Turner, 2015: char. 274, modified from Pol et al., 2009: char. 274)**: Longitudinal keels on dorsal surfaces of osteoderms: restricted to the posterior edge of osteoderm (0); not restricted to the posterior edge (1).

**Character 275 (Pol et al., 2009: char. 275)**: Jugal, anteriorly on lateral surface below orbits: lacks a depression (0); possesses a depression (1).

**Character 276 (Schwarz et al., 2017: char. 276, modified from Pol et al., 2009: char. 276)**: Transverse ridge crossing the frontal anteromedial to the orbits: absent (0); present as a ridge (1); present as prominent anteriorly-curved shelf (transverse interorbital crest sensu Andrade and Hornung 2011) (2)

**Character 277 (Pol et al., 2009: char. 277)**: Shallow hemispherical depression on the lacrimal and/or prefrontal anterior to orbital margin (not articulation facet for palpebral): absent (0); present (1).

**Character 278 (Pol et al., 2009: char. 278)**: Anterior half of interfenestral bar between suborbital fenestrae: lateral margins are parallel to subparallel (0); flared anteriorly (1).

**Character 279 (Pol et al., 2009: char. 279)**: Posterior half of interfenestral bar between suborbital fenestrae: lateral margins are parallel to subparallel (0); flared posteriorly (1).

**Character 280 (Pol et al., 2009: char. 280)**: Angular, shape of posteroventral margin: straight or gently arched dorsally (0); strongly arched dorsally (1).

**Character 281 (Pol et al., 2009: char. 281)**: Squamosal, lateral margin of dorsal surface: squared off with continuous ear valve groove (0); bears a prominent depressed area just anterior to posterior lobe of squamosal, groove for ear valve discontinuous (1).

**Character 282 (Pol et al., 2009: char. 282)**: Fibula, shaft distal to iliofibularis trochanter: straight (0); bowed posteriorly (1).

**Character 283 (Buckley & Brochu, 1999: char. 106)**: Scapular blade width: no more than twice the length of the scapulocoracoid articulation (0); scapular blade very broad and greater than twice the length of scapulocoracoid articulation (1).

**Character 284 (Pritchard et al., 2013: char. 284, modified from Buckley et al., 2000: char. 115)**: Vomer, exposure on palate: vomer contributes flattened plate to secondary palate (0); vomer forms no part of secondary palate (1).

**Character 285 (Turner & Buckley, 2008: char. 285)**: Supraoccipital, when present on dorsal skull roof: with narrow exposure, parietal forms portion of occipital surface (0); with broad exposure, parietal does not form portion of occipital surface (1).

**Character 286 (Turner & Buckley, 2008: char. 286)**: Jugal, anterior and posterior processes: inline dorsoventrally (0); anterior and posterior processes at a sharp angle to one another, both processes slope ventrally to form a strongly arched jugal (1).

**Character 287 (Turner & Buckley, 2008: char. 287)**: Lateral expansion of posterodorsal edge of surangular anterior to glenoid fossa: absent (0); present (1).

**Character 288 (Turner & Buckley, 2008: char. 288)**: In lateral view, anterior process of the squamosal extending to the orbital margin, overlapping the postorbital: absent (0); present (1).

**Character 289 (Turner & Buckley, 2008: char. 289)**: In lateral view, surangular and dentary suture: simple, with little or no interdigitation (0); suture complex, with interlocking prongs from both surangular and dentary, three posterior prongs from dentary and two from surangular (1).

**Character 290 (Turner & Buckley, 2008: char. 290)**: Prominent depression on palate near alveolar margin at level of sixth or seventh alveolus: absent (0); present (1).

**Character 291 (Andrade & Bertini, 2008: char. 103)**: Pterygoid, ventral surface of pterygoid flanges, parachoanal fossae: absent (0); present (1).

**Character 292 (Turner & Sertich, 2010: char. 292)**: Pterygoid, in ventral view, participation in suborbital fenestra: pterygoid forms margin of suborbital fenestra (0); excluded from suborbital fenestra by ectopterygoid-palatine contact (1).

**Character 293 (Turner & Sertich, 2010: char. 293)**: Maxilla, lateral surface along alveolar margin, conformation of the neurovascular foramina: foramina absent or form a single continuous row (0); gap in foramina between an anterior series and a posterior series (1).

**Character 294 (Turner & Sertich, 2010: char. 294)**: Surface of tooth enamel: smooth or slightly crenulated (0); with ridges at base of crown (often extending apically) (1).

**Character 295 (Turner & Sertich, 2010: char. 295)**: Posterior (molariform) teeth, wear facets: absent (0); present (1).

**Character 296 (Turner & Sertich, 2010: char. 296)**: Tooth (with transitional morphology) present at premaxilla-maxilla contact: absent (0); present (1).

**Character 297 (Turner & Sertich, 2010: char. 297)**: Basioccipital, midline crest on basioccipital plate below occipital condyle: absent (0); present (1).

**Character 298 (Turner & Sertich, 2010: char. 299)**: Dorsal osteoderms, accessory ranges of osteoderms (sensu Frey, 1988): absent (0); present (1).

**Character 299 (Andrade & Bertini, 2008: char. 131)**: Maxillary tooth, size relative to maxillary palatal surface in palatal view: proportionally small teeth, occupying only marginal portion of ventral surface of maxilla (0); proportionally well-developed teeth, occupying large area of maxillary palatal surface (1).

**Character 300 (Jouve, 2004: char. 68 & Jouve, 2009: char. 75)**: Ventral lamina of jugal: extends far anterior to ectopterygoid (0); ends at the level of the ectopterygoid (1).

**Character 301 (Sereno & Larsson, 2009: char. 199, modified from Norell, 1988: char. 42 & Brochu, 1997: char. 51)**: Surangular extension toward posterior end of retroarticular process: along entire length (0); pinched off anterior to posterior tip (1).

**Character 302 (Turner, 2015: char. 302)**: Muscle attachment scars on ventral surface of quadrate ramus: form modest crests (0); prominent knobs (1).

**Character 303 (Turner, 2015: char. 303)**: Pterygoid flange shape: mediolaterally broad, reaching laterally beyond medial margin of quadrate condyles (0); relatively narrow, does not reach laterally to medial margin of quadrate condyles (1).

**Character 304 (Turner, 2015: char. 304)**: In ventral view, posterior process of maxilla relative to ITF excluded from ITF (0); forms part of ITF (1).

**Character 305 (Turner, 2015: char. 305)**: Highly modified ectopterygoid, mediolaterally broad and flattened with greatly expanded: absent (0); present, robust anterior process larger than posterior process (1); present, anterior and posterior process roughly equal in size (2).

**Character 306 (Turner, 2015: char. 306)**: In ventral view, palate medial to toothrow: forms a single continuous surface (0); ridge running on the palate medial to toothrow formed by maxilla and ectopterygoid (1).

**Character 307 (Turner, 2015: char. 307)**: Maxillary tooth row, penultimate and ultimate maxillary teeth enlarged and highly modified crushing tooth: absent (0); present (1).

**Character 308 (Turner, 2015: char. 308)**: Prefrontals: do not meet at midline (0); meet at midline (1).

**Character 309 (Turner, 2015: char. 309)**: Pear shaped external naris: absent (0); present (1).

**Character 310 (Turner, 2015: char. 310)**: Skull, dorsal surface at parietal-squamosal contact surface: continuous across suture (0); suture marked by groove or sulcus (1).

**Character 311 (Turner, 2015: char. 311)**: Maxilla, lateral surface, continuous groove or sulcus extending from orbital margin towards narial opening: absent (0); present (1).

**Character 312 (Turner, 2015: char. 312)**: Maxilla, posteromedial process curving posteriorly onto palatine formed nasopharyngeal passage: absent (0); present (1).

**Character 313 (Turner, 2015: char. 313)**: Squamosal, posterior half, dorsal and ventral rims of groove for external ear valve musculature: thin or parallel sided (0); flared posteriorly (1).

**Character 314 (Turner, 2015: char. 314)**: Lacrimal, in dorsal view, anterior extent on rostrum relative to prefrontal: prefrontal extends farther anteriorly (0); lacrimal extends farther anteriorly (1); lacrimal and prefrontal subequal in anterior extent (2).

**Character 315 (Turner, 2015: char. 315)**: Lacrimal, in dorsal view, mediolateral width relative to prefrontal: equal to or less than width of prefrontal (0); wider than prefrontal (1).

**Character 316 (Turner, 2015: char. 316)**: Premaxillae, degree of contact posterior to the incisive foramen: extensive contact (0); narrow contact (1).

**Character 317 (Turner, 2015: char. 317)**: Posterior margin of the palatines where they form the floor of the nasopharyngeal passage, shape: V- or U-shaped (0); straight (1).

**Character 318 (Turner, 2015: char. 318)**: Posterior margin of the choanal groove, location: anteriorly on the pterygoids (0); posteriorly on the pterygoids near the posterior margin of pterygoids (1).

**Character 319 (Turner, 2015: char. 319)**: Pterygoid-palatine contact, ventral aspect of palate, shape of the suture: transverse, or nearly so (0); prong of pterygoid projects anteriorly (1).

**Character 320 (Turner, 2015: char. 320)**: On palate, foramen located on premaxilla/maxilla suture near the alveolar border: absent (0); present (1).

**Character 321 (Turner, 2015: char. 321)**: Ectopterygoid/pterygoid contact in ventral view: complex, anterior part of ectopterygoid forming suture whereas posterior part of ectopterygoid overlaps the pterygoid (0); sutured along entire contact, no part of ectopterygoid overlapping pterygoid (1).

**3. Character and state descriptions for the matrix adapted from Tennant et al. (2016) (Supplementary Dataset S3)**

Cranial Characters

1. Dorsal cranial bones (skull roof, cranial table), external surface: smooth (0); ornamented (1) (Clark, 1994).
2. Dorsal cranial bones (skull roof, cranial table), external surface: slightly grooved (0); heavily ornamented with deep pits and/or grooves (1); with shallow pits (2) (Clark, 1994) (character state 2 added here; added ‘/or’ to state 1).
3. Snout, external surface, sculpting: absent (0); present but to a lesser degree than cranial table (1); present, as prominent as on cranial table (2) (Gasparini *et al*., 2006) (character state 1 added here; added ‘as prominent as on cranial table’ to character state 2) [**ordered**].
4. Rostrum, dorsal projection posterior to the external nares, relative to remainder of rostrum: absent, rostrum straight or low (0); rostrum upturned (1) (Andrade *et al* 2011) (added ‘posterior to the external nares, relative to remainder of rostrum’).
5. Skull, lateral expansion at orbits relative to rostrum: gradual (0); abrupt (1) (Clark, 1994) (added ‘lateral’ and ‘relative to rostrum’).
6. Snout, lateral contour, in dorsal view: straight (0); sinusoidal (‘festooned’) (1) (Ortega *et al*., 2000).
7. Snout, overall proportions: narrow oreinirostral (tall and domed) (0); nearly tubular (1); platyrostral (broad and flat) (2) (Clark, 1994; sensu Rayfield and Milner, 2008) [**ordered**].
8. Snout, profile of dorsal edge in lateral view (anterior to cranial table): concave (0); convex (1); approximately straight (2) (Sweetman *et al*., 2015).
9. External nares, orientation: facing anterolaterally (0); facing dorsally (1); facing anterodorsally (2) (Tennant et al., 2016, modified from Clark, 1994)
10. External nares: completely divided by a septum (0); partially divided posteriorly (1); confluent, no indication of a septum (2) (Clark, 1994) [**ordered**].
11. Orbit, orientation: facing anterolaterally (0); facing fully laterally (1) (Wilkinson *et al*., 2008).
12. Orbit, anterolateral border: continuous margin (0); develops as a small groove into pre-orbital elements (1) (Tennant et al., 2016).
13. External antorbital fenestra: large, greater than 0.5 times the size of the orbit (0) small, less than or equal to 0.5 times the size of the orbit (1); absent (2) (Tennant et al., 2016, modified from Andrade *et al*., 2011) [**ordered**].
14. Antorbital fenestra, shape: rounded or dorsoventrally high (0); dorsoventrally low and anteroposteriorly elongate, slit-like (1) (2) (Gasparini *et al*., 2006).
15. External supratemporal fenestra: present (0); absent (1) (Ortega *et al*., 2006).
16. External supratemporal fenestra: perforated (0); imperforated (1) (Tennant et al., 2016, adapted from Joffe, 1967).
17. External supratemporal fenestra, shape: square to sub-rectangular (0); circular to subcircular (1); mediolaterally narrow and slit-like (2) (Tennant et al., 2016, modified from Andrade *et al*., 2011)
18. External supratemporal fenestra, maximum anteroposterior length: equal to or shorter than orbits (0); longer than orbits (1) (Tennant et al., 2016, modified from Clark, 1994).
19. Intertemporal mediolateral width (minimum between supratemporal fenestrae), relative to interorbital mediolateral width (minimum between orbits): intertemporal region broader (0); intertemporal region equal or narrower (1) (Tennant et al., 2016)
20. Lateral temporal fenestra in lateral view, size proportional to orbit in dorsal view: small to absent, no more than 20% of the area of the orbit (0); more than 20 to less than 50% of the area of the orbit (1); area is larger than 50% of the area of the orbit (2) (Tennant et al., 2016, modified from Andrade *et al*., 2011) [**ordered**].
21. Lateral temporal fenestra, orientation: faces laterally (0); faces dorsolaterally (1) (Andrade *et al*., 2011).
22. Lateral temporal fenestra, shape: triangular (0); elliptical to subpolygonal (1) (Ortega *et al*., 2000).
23. Suborbital fenestra: small, less than 50% of orbital area (0); between 50% and the same size as the orbit (1); larger than the orbit (2) (Tennant et al., 2016, modified from Andrade *et al*., 2011) [**ordered**].
24. Intermandibular angle (degrees): lower than 40° (0); 40-45 (1); 46-50° (2); greater than 50° (3) (Tennant et al., 2016) [**ordered**].
25. Skull length: skull width, ratio: less than 2.0 (0); 2.0 to less than 2.5 (1); 2.5 or greater (2) (Tennant et al., 2016) [**ordered**].
26. Skull length: snout length, ratio: less than 2.0 (0); 2.0 to less than 2.5 (1); 2.5 to less than 3.0 (2); 3.0 or greater (3) (Tennant et al., 2016) [**ordered**].
27. Skull length: orbit length, ratio: less than 3.0 (0); 3.0 to less than 4.0 (1) 4.0 to less than 5.0 (2); 5.0 or greater (3) (Tennant et al., 2016) [**ordered**].
28. Skull width: orbit width, ratio: less than 2.5 (0); 2.5 to less than 3.5 (1); 3.5 or greater (2) (Tennant et al., 2016) [**ordered**].
29. Skull length: supratemporal fenestra length, ratio: less than 6.0 (0); 6.0 to less than 7.0 (1); 7.0 to less than 8.0 (2) 8.0 or greater (3) (Tennant et al., 2016) [**ordered**].
30. Skull width: supratemporal fenestra width, ratio: less than 3.0 (0); 3.0 to less than 5.0 (1); 5.0 to less than 6.0 (2); 6.0 or greater (3) (Tennant et al., 2016) [**ordered**].
31. Premaxilla, maximum mediolateral width of paired premaxillae relative to that of the rostrum at the level of alveoli 4 or 5: premaxillae equal or narrower (0); rostrum narrower (1) (Jouve, 2009).
32. Premaxilla, anterior to nares: narrower than, or equal to, twice the anterior nasal mediolateral width (0); broader than twice the anterior nasal width (1) (Tennant et al., 2016, modified from Clark, 1994).
33. Premaxilla-maxilla, distance between the anterior tip of the snout and the anteriormost position of the premaxilla-maxilla suture in dorsal view, relative to the distance between the anteriormost position of the premaxilla-maxilla suture and the posterodorsal extremity of the premaxilla in dorsal view: distance between the tip of the snout and the anteriormost position of the premaxilla-maxilla suture larger (0); distances approximately equal (1); distance between the anteriormost position of the premaxilla-maxilla suture and the posterodorsal extremity of the premaxilla larger (2) (Jouve, 2004) [**ordered**].
34. Premaxilla-maxilla suture, small foramen in lateral surface (not for large mandibular teeth): absent (0); present (1) (Pol, 1999a).
35. Premaxilla, projection of the internarial bar relative to the main body of premaxilla and narial opening: does not project anterior to the main body of the premaxilla (0); strongly projected anteriorly from narial opening, extending anterior to main body of maxilla (1) (Andrade *et al*., 2011).
36. Premaxilla, participation in internarial bar: forming at least the anterior half (0); with little participation (1) (Tennant et al., 2016, modified from Clark, 1994)
37. Premaxilla, ventral edge relative to maxilla: lower than ventral edge of maxilla, with dorsal contour of anterior part of dentary strongly concave to accommodate (0); at same height as ventral edge of maxilla (1); premaxilla ventral edge dorsal to maxilla (2) (Tennant et al., 2016, modified from Ortega *et al*., 2000) [**ordered**].
38. Premaxilla, perinarial crests: absent (0); present as well-defined and distinct ridges, cornering the lateral to posterior borders of the naris (1) (Andrade *et al*., 2011).
39. Premaxilla, notch on lateral edge of external nares: absent (0); present on the dorsal half of the lateral edge of the external nares (1) (Pol, 1999a).
40. Premaxilla, perinarial fossa: absent (0); present (1) (Pol and Apesteguía, 2005)
41. Premaxilla, postnarial fossa: absent (0); present (1) (Andrade *et al*., 2011).
42. Premaxilla-maxilla, suture: confluent ventrally (0); opened contact on ventral edge of rostrum (1) (Clark, 1994).
43. Premaxilla-maxilla contact, orientation in dorsal view, whether or not posterodorsal process is present: anteromedially directed (0); posteromedially directed (1) (Tennant et al., 2016, modified from Schwarz and Salisbury, 2005)
44. Premaxilla, posterodorsal process: absent (0); present, extending posteriorly and wedging between maxillae and nasals (1) (Pol, 1999a).
45. Premaxilla, orientation of anterior alveolar margin: vertical (0); out-turned (1) (Tennant et al., 2016, modified from Sereno *et al*., 2001)
46. Maxillae, posterior palatal branches anterior to palatines: do not meet (0); meet (1) (Clark, 1994).
47. Maxilla-premaxilla, suture in palatal view medial to alveolar region: sinusoidal, posteromedially directed on lateral half and anteromedially directed along medial region (0); posteromedially-directed (1) (Tennant et al., 2016, modified from Pol, 1999a).
48. Maxilla-premaxilla, lateral fossa excavating alveolus of last premaxillary tooth: absent (0); present (1) (Larsson and Sues, 2007).
49. Maxilla, depression on posterolateral surface, laterally positioned: absent (0); present (1) (Tennant et al., 2016, modified from Wu *et al*., 1997).
50. Maxilla, depression on anterolateral surface, medially positioned: absent (0); present (1) (Tennant et al., 2016).
51. Maxilla, lateral surface of jugal process (posterior portion): heavily striated (0); ornamented, like rest of rostrum (1); smooth (2) (Tennant et al., 2016).
52. Maxilla, ventral edge in lateral view: straight or convex (0); sinusoidal (1) (Ortega *et al*., 2000).
53. Maxilla and premaxilla, general shape of external surface: single plane facing laterally (0); with ventral region facing laterally and dorsal region facing dorsolaterally (1) (Pol, 1999a).
54. Maxilla, presence of occlusal pit for reception of enlarged dentary tooth anterior to dental arcade (or M2): present (0); absent (1) (Tennant et al., 2016, adapted from Martin *et al*., 2014).
55. Maxilla, evaginated alveolar edges: absent (0); present (1) (Gasparini *et al*., 2006).
56. Maxilla, lateral surface, unsculpted region along alveolar margin: absent (0); present (1) (Wu and Sues, 1996).
57. Maxilla, sculpturing of palatal surface: absent, palatal surface smooth (0); present anteriorly, absent posteriorly (1); present throughout, palatal surface ornamented with ridges (2) (Ortega *et al*., 2000) [**ordered**].
58. Maxilla, foramen on palatal surface, dorso-medial to enlarged 5th tooth: absent (0); present (1); develops elongate groove (2) (Tennant et al., 2016) [**ordered**].
59. Maxillary teeth, dental implantation, middle teeth: confluent, located in dental groove (0); in isolated alveoli (1) (Tennant et al., 2016).
60. Maxillary teeth, dental implantation, posterior teeth: confluent, located in dental groove (0); in isolated alveoli (1) (Tennant et al., 2016).
61. Maxilla, palatine process: absent (0); present, next to the anterior border of suborbital fenestrae (1) (Andrade and Bertini, 2008).
62. Maxilla-nasal, suture, orientation with respect to snout lateral margins: parallel (0); oblique (1) (Tennant et al., 2016).
63. Nasal, participation in dorsal margin of external nares: present (0); absent (1) (Tennant et al., 2016).
64. Nasal participation in margins of external nares: present posteriorly (0); present posteriorly and medially (1) (Clark, 1994).
65. Nasals: paired and unfused (0); partially or completely fused (1) (Tennant et al., 2016, modified from Gasparini *et al*., 2006).
66. Nasal, lateral border posterior to external nares: concave (0); straight (1); convex (2); sinusoidal (3) (Tennant et al., 2016, modified from Pol, 1999a).
67. Nasal, lateral edges: sub-parallel (0); oblique to one another, converging anteriorly (1) (Pol, 1999a).
68. Nasal, participation in antorbital fenestra: present (0); absent (1) (Ortega *et al*., 2000).
69. Nasal, posterior portion of the dorsal surface: anteroposteriorly crenulated (0); smooth or sculpted as rest of rostrum (1) (Tennant et al., 2016).
70. Nasals, posterior mediolateral widening adjacent to the maxilla (anterior to contact with peri-orbital elements): abrupt (0); gradual (constant) (1); (Tennant et al., 2016, adapted from Lauprasert *et al*., 2011).
71. Nasal-lacrimal contact: present (0); absent (1) (Clark, 1994).
72. Nasal-lacrimal contact: along medial surface of lacrimal (0); lacrimal forms a point contact with nasal (1) (Tennant et al., 2016, modified from Clark, 1994)
73. Nasal, posterior tips of nasals: converge along the sagittal plane (0); separated by anterior projection of frontals (1) (Ortega *et al*., 2000).
74. Nasal, posterior portion and anterior portion of frontal, midline anteroposterior depression: absent (0); present (1) (Montefeltro *et al*., 2011).
75. Lacrimal, total anteroposterior length relative to anteroposterior length of prefrontal: longer (0); shorter or equal to (1) (Tennant et al., 2016, modified from Brochu, 1999)
76. Lacrimal, shape: anteroposteriorly longer than mediolaterally broad (0); as anteroposteriorly long as mediolaterally broad (1) (Sereno and Larsson, 2009).
77. Lacrimal, posterior extent and relationship with jugal: extending posteroventrally, widely contacting jugal (0); tapering posteroventrally, does not contact jugal or only point contact with jugal (1) (Zaher *et al*., 2006).
78. Lacrimal and jugal, anterior margins: confluent, with no notch at the anterior contact (0); jugal edge convex, producing an anterior notch at contact (filled with maxilla) (1) (Larsson and Sues, 2007).
79. Jugal, anterior extension below orbit, in dorsolateral view: does not extend beyond anterior margin of orbit (0); extends beyond anterior margin (1) (Pol, 1999a).
80. Jugal, dorsoventral depth of orbital portion in relation to infratemporal portion: almost the same to less than twice the depth (0); orbital portion twice the depth of the infratemporal portion (1) (Tennant et al., 2016, modified from Clark, 1994).
81. Jugal, foramen on the lateral surface near the anterior margin: absent (0); present (1) (Zaher *et al*., 2006).
82. Jugal, anterior process length relative to infratemporal fenestrae anteroposterior length: 1.0 or less times the length (0); longer than 1.0 times the length (1) (Tennant et al., 2016, modified from Larsson and Sues, 2007).
83. Jugal, orientation of base of postorbital process: directed posterodorsally (0); directed dorsally (1) (Pol, 1999a).
84. Jugal, location of postorbital process relative to main jugal body: anteriorly placed (0); in the middle (1); posteriorly positioned (2) (Pol, 1999a) [**ordered**].
85. Jugal portion of postorbital bar, relative to lateral surface of jugal: flush with lateral surface of jugal (0); inset (1) (Ortega *et al*., 2000).
86. Jugal, anterior fossa: bordered by ornamented ridge (0); continuous with lateral surface (1) (Tennant et al., 2016).
87. Jugal, anteroposterior ridge on lateral surface below infratemporal fenestrae: absent (0); present (1) (Pol and Norell, 2004b).
88. Jugal-ectopterygoid, suture ridge: absent (0); present (1) (Montefeltro *et al*., 2011).
89. Preorbital elements, anterior palpebral bone: no notable depression or projection (0); marked depression, developing into an incipient lateral projection (1); marked depression forming a prominent lateral projection for the support of the anterior palpebral (2) (Tennant et al., 2016, modified from Sereno and Larsson, 2009 [**ordered**].
90. Prefrontal, lateral development: reduced, no notable lateral projection (0); enlarged, extending laterally or posterolaterally over orbit (1) (Tennant et al., 2016, modified from Gasparini *et al*., 2006).
91. Prefrontal, anterior morphology: tapers anteriorly to a point (0); anteriorly broad (1)
92. Prefrontal-frontal sutures, form paired dorsal crests: absent (0); present (1) (Pol and Powell, 2011).
93. Prefrontal-lacrimal suture, crest: absent (0); present, situated anterior to orbit (1) (Tennant et al., 2016, modified from Andrade *et al*., 2011).
94. Prefrontal and lacrimal around orbits: forming flat rims (0); evaginated, forming elevated rims from the dorsal surface of the skull (1) (Gasparini *et al*., 2006).
95. Prefrontal pillars (ventral process): not contacting palate (0); contacting palate (1) (Clark, 1994).
96. Frontals: unfused (0); fused (1) (Tennant et al., 2016, modified from Clark, 1994)
97. Frontal, mediolateral width of paired frontals between orbits: broader than nasals (0) equal or narrower than nasals (1) (Tennant et al., 2016, modified from Clark, 1994)
98. Frontals, mediolateral width between orbits: narrower than posterior end (posterior end flares laterally posterior to orbits) (0); equal width or broader than posterior end (1) (Tennant et al., 2016, adapted from Sweetman *et al*., 2015).
99. Frontal, morphology of anteriormost border of anterior process: truncated (0); wedge-like (1); bifurcated (2) (Tennant et al., 2016, modified from Andrade *et al*., 2011) [**ordered**].
100. Frontals, dorsal surface: flat (0); with anteroposterior ridge along midline suture (1) (Tennant et al., 2016, modified from Clark, 1994) [**ordered**] (note: see Schwarz and Salisbury [2005] for discussion of how this character relates to ontogeny).
101. Frontal, dorsal anteroposterior ridge(s): restricted to the posterior portion (0); restricted to median portion (1); restricted to anterior portion (2); occupy entire length of frontal (3) (Tennant et al., 2016, modified from Montefeltro *et al*., 2011)
102. Frontal, anterior extension of anterior margin: level with, or anterior to, the orbits (0); does not reach the anterior margin of the orbits (1) (Andrade *et al*., 2011).
103. Frontal, participation in orbit border: forming great part of posterior, medial and anterior (or anteromedial) regions (0); restricted to posterior and posteromedial region (1); restricted to medial margin (2) (Tennant et al., 2016, modified from Montefeltro *et al*., 2013) [**ordered**].
104. Frontal, transverse ridge crossing anteromedial to the orbits: absent (0); present (1) (Pol *et al*., 2009).
105. Frontal, participation in supratemporal fenestra: absent (0); present (1) (Tennant et al., 2016).
106. Frontal, participation in supratemporal fenestra: anteromedially (0), anteriorly only (1) (Tennant et al., 2016).
107. Frontal, anterior ramus with respect to the anterior tip of the prefrontal: does not extend past the anterior tip (0); extends beyond the anterior tip of the prefrontal (1) (Tennant et al., 2016, modified from Sereno *et al*., 2001)
108. Frontal, lateral margin relative to the skull surface: flush (0); elevated, forming ridged orbital margins (1) (Brochu, 1999).
109. Frontal, anterior process constriction with respect to main body of frontal, excluding sagittal projection into nasals anterior to orbits: absent, lateral edges parallel to sub-parallel (0); present, anterior portion mediolaterally constricted, with convergent lateral margins (1) (Tennant et al., 2016, modified from Montefeltro *et al*., 2013).
110. Postorbital, anterolateral process: absent or poorly-developed (0); well-developed, long and distally acute (1) (Clark, 1994).
111. Postorbital-jugal contact, configuration: postorbital medial to jugal (0); postorbital dorsal to jugal (1) (Clark, 1994).
112. Parietal-postorbital suture: absent from the dorsal surface of the skull roof (0); present on the dorsal surface of the skull roof (1) (character broken down into characters 107-109 by Tennant et al., 2016) (Clark, 1994).
113. Parietal-postorbital suture: absent from the supratemporal fossa (0); present within the supratemporal fossa (1) (Tennant et al., 2016).
114. Parietal-postorbital suture within the supratemporal fossa: present within the ventral region (0); broadly present (1) (Tennant et al., 2016).
115. Parietal, dorsal surface: projects dorsally, relative to the skull roof (0); same level as squamosal (1); depressed relative to the squamosal (2) (Tennant et al., 2016, modified from Andrade *et al*., 2011) [**ordered**].
116. Parietal, posterior region dorsal surface: smooth (0); presenting a anteroposterior dorsal ridge (1); marked ventral deflection (‘bevelled’) in posterior portion (2); sculpted as with the rest of the skull table (3) (Tennant et al., 2016, modified from Montefeltro *et al*., 2013).
117. Parietal-squamosal emargination (anterior concavity at suture contact), posterior margin in dorsal view: absent (0); present (1) (Wilkinson *et al*., 2008).
118. Supratemporal roof, dorsal surface: postorbital and squamosal with flat shelves extending laterally beyond quadrate contact (0); lateral edge terminating medial or immediately dorsal to medial-most point of contact with quadrate (1) (Tennant et al., 2016, modified from Clark, 1994).
119. Supratemporal fenestra, medial border: flat, sculpted region (or unsculpted if rest of cranial table unsculpted) (0); forming a low sagittal crest (rims) posteriorly (1); forming a low sagittal crest (rims) along full length (2) (Tennant et al., 2016, modified from Clark, 1994) [**ordered**].
120. Supratemporal fenestra, dermal bone overhang: absent (0); present (1) (Norell, 1988).
121. Supratemporal fenestra, dermal bone overhang: present only medially and posteriorly (0); present about the entire edge (1) (Norell, 1988).
122. Supratemporal fenestra, shallow fossa at anteromedial corner: present (0); absent, corner smooth (1) (Brochu, 1999).
123. Supratemporal medial rims: continuous with orbital rims (0); separated from orbital rims by the postorbital bar (1) (Tennant et al., 2016).
124. Supratemporal medial rims, extend posteriorly to contact posterior skull margin: present (0); absent (1) (Tennant et al., 2016).
125. Supratemporal fenestra, relative contribution of frontal and parietal to medial margin: parietal with equal or greater contribution (0); frontal excluded from margin (1) (Tennant et al., 2016).
126. Supratemporal fenestrae, minimum width between fenestrae, with respect to maximum width of cranial table: one third or less of total width (0); more than one third of total width (1) (Tennant et al., 2016, adapted from Schwarz and Salisbury, 2005).
127. Orbitotemporal channel, size of the dorsal aperture: area of foramen less than or equal to 30% of that of the internal supratemporal fenestra (0); larger than 30% of the internal supratemporal fenestrae area (1) (Montefeltro *et al*., 2013).
128. Postorbital bar between orbit and supratemporal fossa, shape: broad and solid, as broad as dorsal surface of the cranial table lateral to the supratemporal fenestra (0); much narrower (1); much narrower and connected to orbit via a thin, superficial furrow in postorbital (2) (Tennant et al., 2016, modified from Clark, 1994) [**ordered**].
129. Postorbital bar between orbit and supratemporal fenestra, external texture: sculpted (0); unsculpted (1) (Clark, 1994).
130. Postorbital bar, lateral surface formed by: postorbital and jugal (0); only by postorbital (1) (Gasparini *et al*., 2006).
131. Postorbital bar, shape: transversely flattened (0); transversely broad, with elliptical cross-section (1); slender and cylindrical (2) (Clark, 1994).
132. Postorbital bar, shape of dorsal end nearing skull table: continuous with dorsal part of postorbital (0); dorsal part of postorbital bar constricted, distinct from the dorsal part of the postorbital (1) (Clark, 1994).
133. Cranial table (skull roof), width with respect to ventral portion of skull: as wide as ventral portion of skull (quadrates covered by squamosal) (0); narrower, but still covering more than half of the mediolateral region of quadrates (1); narrower, exposing more than half of mediolateral region of quadrate (2) (Tennant et al., 2016, modified from Wu *et al*., 1997) [**ordered**].
134. Squamosal and postorbital, lateral margins, dorsal view excluding posterolateral process: parallel (0); diverging posteriorly (1); medially concave (2); converging posteriorly (3) (Tennant et al., 2016, modified from Ortega *et al*., 2000).
135. Squamosal, lateral surface, longitudinal groove for attachment of the upper ear lid: absent (0); present (1) (Clark and Sues, 2002).
136. Squamosal groove for upper ear lid: ventral edge is laterally displaced relative to dorsal edge (0); ventral edge is directly beneath dorsal edge (1) (Clark and Sues, 2002).
137. Squamosal, dorsal edge of groove for dorsal ear lid: parallel to ventral edge (0); dorsal margin with a medial curvature (1) (Montefeltro *et al*., 2013).
138. Squamosal, dorsolateral edge: straight and parallel to skull roof (0); bevelled ventrally, with anterolateral notch (1) (Tennant et al., 2016, adapted from Schwarz and Salisbury, 2005).
139. Squamosal, posterolateral process: present (0); absent (1) (Tennant et al., 2016, adapted from Buscalioni and Sanz, 1990).
140. Squamosal, posterolateral process: level with skull table (0); depressed from skull table (1) (Sereno and Larsson, 2009).
141. Squamosal, posterolateral process projection: ventrally directed, confluent with ventral rim of groove for the earflap (0); posteriorly directed and parallel to skull roof (1) (Tennant et al., 2016, modified from Ortega *et al*., 2000).
142. Squamosal, posterolateral process, ornamentation: absent (0); present (1) (Larsson and Sues, 2007).
143. Squamosal, posterolateral process, distal end: tapered and pointed (0); broad and rounded (1) (Larsson and Sues, 2007).
144. Squamosal, anterior process extending anteriorly to the orbital margin, overlapping the postorbital, in lateral view: absent (0); present (1) (Turner and Buckley, 2008).
145. Squamosal, obliquely-oriented ridge on dorsal surface: present (0); absent (1) (Tennant et al., 2016).
146. Squamosal, oblique ridge on dorsal surface, position with respect to the supratemporal fenestra: posterior to supratemporal fenestra (0); posterolateral or lateral to supratemporal fenestra (1) (Tennant et al., 2016).
147. Squamosal-parietal suture: flat, not elevated from the skull table (0); forms a well-developed anteroposterior groove (often bounded by elevated ridges) (1) (Tennant et al., 2016, adapted from Buffetaut, 1983).
148. Squamosal, anterior portion nearing orbital edge: sculpted or unsculpted, consistent with the rest of the skull table (0); sculpting pattern changes (1) (Tennant et al., 2016).
149. Quadratojugal, ornamentation at base (dorsolateral surface): absent (0); present (1) (Pol, 1999a).
150. Quadratojugal, length of anterior process relative to the lower temporal bar: absent or less than one third of lower temporal bar (0); one third to one half the length of the lower temporal bar (1); long, greater than half of the lower temporal bar (2) (Tennant et al., 2016, modified from Larsson and Sues, 2007) [**ordered**].
151. Quadratojugal, shape of posterolateral end and relationship with quadrate: acute or rounded, tightly overlapping quadrate (0); sinusoidal ventral edge, and wide and rounded posterior edge slightly overhanging lateral surface of quadrate (1) (Pol and Norell, 2004a).
152. Quadratojugal, contribution to the lateral temporal fenestra, in dorsal view: extensive contact with the ventral and posterior margins (0); contributes to the posterior and dorsal margin (1); only contributes to the posterior margin (2) (Tennant et al., 2016).
153. Quadratojugal-postorbital contact, in lateral view: not in contact (0); small, point contact (1); broad contact between the quadratojugal and the posterior portion of the postorbital descending flange (2) (Tennant et al., 2016, modified from Clark, 1994) [**ordered**].
154. Infratemporal fenestra, posterior margin, dorsal view: straight (0); with an anterior projection, forming an acute angle (1) (Tennant et al., 2016, modified from Ortega *et al*., 2000).
155. Quadrate, posterior edge: broad medial to tympanum, gently concave (0); narrow dorsal to otoccipital contact, strongly concave (1) (Clark, 1994).
156. Quadrate, dorsal surface fenestration: absent (0); present (1) (Clark 1994).
157. Otic aperture (not including additional quadrate fenestrae): open posteriorly (0); closed posteriorly by quadrate and otoccipital (1) (Clark, 1994).
158. Quadrate, distal body: anterior margin oriented at a right angle in relation to quadratojugal (0); anterior margin gently slopes relative to quadratojugal (1) (Montefeltro *et al*., 2011).
159. Quadrate, pterygoid ramus: with flat ventral edge (0); with deep groove on ventral surface (1); rod-like (2) (Tennant et al., 2016, modified from Clark, 1994).
160. Quadrate, anterodorsal ramus in ventral view: developed, forming more than or equal to 50% of the lateral edge of the internal supratemporal fenestra (0); restricted, forming less than 50% of the lateral edge of the supratemporal fenestra (1) (Montefeltro *et al*., 2011).
161. Quadrate, ventral surface: smooth, with simple muscle scars (0); with multiple developed ridges (1) (Ősi *et al*., 2007).
162. Quadrate, condyles: with poorly developed intercondylar groove (0); medial condyle expands ventrally, being separated from the lateral condyle by a deep intercondylar groove (1) (Ortega *et al*., 2000).
163. Quadrate, development of distal body ventral to otoccipital-quadrate contact: distinct, developing posteroventrally to contact (0); indistinct, not surpassing contact (1) (Tennant et al., 2016, modified from Wu *et al*., 1997).
164. Quadrate, dorsoventral height of the proximal region: less than or equal to 50% of the skull roof total width (0); more than 50% of the skull roof total width (1) (Tennant et al., 2016, modified from Montefeltro *et al*., 2013).
165. Cranio-quadrate canal: opened laterally (0); closed laterally (1) (Clark, 1994).
166. Ectopterygoid-maxilla, contact: present (0); absent (1) (Tennant et al., 2016).
167. Ectopterygoid, contribution to postorbital bar: absent (0); present (1) (Sereno and Larsson, 2009).
168. Ectopterygoid, main axis orientation: mediolaterally or slightly anterolaterally (0); anteroposteriorly, subparallel to anteroposterior axis of skull (1) (Tennant et al., 2016, modified from Pol *et al*., 2004).
169. Ectopterygoid, extent of medial projection on the ventral surface of pterygoid flanges: barely extended (0); widely extended, covering approximately the lateral half of the ventral surface of the pterygoid flanges (1) (Zaher *et al*., 2006).
170. Ectopterygoid, anterior process: developed (0); reduced or absent (1) (Pol, 1999a).
171. Ectopterygoid, posterior process: developed (0); reduced or absent (1) (Pol, 1999a).
172. Palatines, palatal processes: do not meet on palate below narial passage (0); meet ventral to narial passage, forming part of secondary palate (1) (Clark, 1994).
173. Palatine-maxilla, suture when fused at midline: palatine anteriorly rounded (0); palatine anteriorly pointed (1); suture transverse to midline axis (2) (Tennant et al., 2016, modified from Brochu, 1999).
174. Interfenestral bar, anterior half between suborbital fenestrae, lateral margins: parallel to subparallel (0); flared anteriorly (1) (Tennant et al., 2016, modified from Pol *et al*., 2009).
175. Interfenestral bar, posterior half between suborbital fenestrae, lateral margins: flared posteriorly (0); parallel to subparallel (1); converge posteriorly (2) (Tennant et al., 2016, modified from Pol *et al*., 2009) [**ordered**].
176. Palatines, anteroposterior axis: run parasagittally (0); diverge laterally, becoming rod-like and forming palatine bars posteriorly (1) (Martinelli, 2003).
177. Palatine-pterygoid, contact on palate: run parasagittally (0); palatines firmly sutured to pterygoids (1) (Pol and Norell, 2004a).
178. Pterygoids, contact with one another on palate: not in contact anterior to basisphenoid on the palate (0); pterygoids in contact (1) (Wu *et al*., 1997).
179. Pterygoid, role of primary palate in forming choanal opening: forms posterior half of choanal opening (0); forms posterior, lateral, and part of anterior margin of choana (1); completely encloses choana (2) (Clark, 1994) [**ordered**].
180. Pterygoid, participation in the suborbital fenestra, ventral view: forms margin of suborbital fenestra (0); excluded from suborbital fenestra by ectopterygoid-palatine contact (1) (Turner and Sertich, 2010).
181. Choanae, anterior edge, location: situated between suborbital fenestrae (or anteriorly) (0); near posterior edge of suborbital fenestrae (1) (Clark, 1994).
182. Choanal opening, conformation in palate: continuous with pterygoid ventral surface except for anterior and anterolateral borders (0); opens into palate through deep choanal groove (1) (Clark, 1994).
183. Choanal groove: undivided (0); partially septated (1); completely septated (2) (Clark, 1994) [**ordered**].
184. Pterygoid, quadrate process: well-developed, extending posterolaterally beyond anterior margin of basioccipital (0); poorly developed, only present as an incipient projection (1) (Tennant et al., 2016, modified from Pol, 1999a).
185. Pterygoid, quadrate ramus, in ventral view: narrow and bar-like (0); broad and laminar (1) (Tennant et al., 2016, modified from Wu *et al*., 1997).
186. Pterygoid, palatal surface: smooth (0); sculpted (1) (Clark, 1994).
187. Pterygoid flanges: mediolaterally expanded, laterally surpassing the quadrate medial condyle (0); relatively short, and do not reach laterally to the level of the quadrate medial condyle (1) (Ősi *et al*., 2007).
188. Basisphenoid, ventral exposure on braincase: exposed on ventral surface of braincase (0); virtually excluded from ventral surface by pterygoid and basioccipital (1) (Clark, 1994).
189. Basisphenoid, lateral exposure on braincase: absent (0); present (1) (Pol, 1999a).
190. Basisphenoid: ventral surface continuous with surrounding bones (0); body ventrally developed and separated from the remaining elements by a posteroventral step formed by a sulcus separating it from the main occipital plane, forming a postchoanal pterygoid-basisphenoid tuberosity (1) (Montefeltro *et al*., 2011).
191. Basisphenoid, ventral surface, mediolateral size relative to basioccipital: shorter than basioccipital (0); equal or longer than basioccipital (1) (Tennant et al., 2016, modified from Clark, 1994)
192. Basioccipital: without well-developed bilateral tuberosities (0); with large, pendulous tubera (1) (Clark, 1994).
193. Basioccipital, midline crest on basioccipital plate below occipital condyle: absent (0); present (1) (Turner and Sertich, 2010).
194. Basioccipital and ventral part of otoccipital, orientation (when skull held horizontally): posteriorly (0); posteroventrally (1) (Gomani, 1997).
195. Otoccipital, ventrolateral contact with quadrate: very narrow, otoccipital only abuts quadrate (0); broad, ventrolateral margin of otoccipital extensively contacts ventromedial portion of quadrate (1) (Tennant et al., 2016, modified from Clark, 1994).
196. Supraoccipital, posterodorsal exposure in skull roof: absent (0); present (1) (Ortega *et al*., 2000).
197. Supraoccipital, posterodorsal exposure: exposed in midline portion of posterior region of skull table (0); restricted to a thin surface attached to posteriormost portion of parietal and squamosal (1) (Montefeltro *et al*., 2011).
198. Supraoccipital, relationship with foramen magnum: forms dorsal edge (0); otoccipitals meet dorsally, separating the foramen magnum from the supraoccipital (1) (Clark, 1994).
199. Cranial nerves IX-XI, passage through braincase: all pass through common large foramen vagi in otoccipital (0); cranial nerve IX passes medial to nerves X and XI in separate passage (1) (Clark, 1994).
200. Mastoid antrum, location: does not extend into supraoccipital (0); extends through transverse canal in supraoccipital to connect middle ear regions (1) (Clark, 1994).

**Mandibular Characters**

201. Mandible, outer surface sculpture, lateral surface: absent (0); present (1) (Montefeltro *et al*., 2011).
202. Mandible, outer surface sculpture, ventral surface: present on dentary (0); present on dentary and splenial (1) (Montefeltro *et al*., 2011).
203. Mandibular symphysis, anteroposterior length relative to mediolateral width: short, length and width subequal or shorter than wide (0); proportionally long, longer than wide (1); extremely long, length at least five times its width (2) (Andrade *et al*., 2011) [**ordered**].
204. Mandibular symphysis, posterior extension, terminating medial to the dentary alveoli: short, up to the D5 (0); to the D5-D6 (1); to the D7 or greater in length (Tennant et al., 2016) [**ordered**].
205. Mandibular symphysis, lateral view: shallow and tapering dorsoventrally anteriorly (0); deep and tapering dorsoventrally anteriorly (1); shallow and anterior margin convex (2) (Tennant et al., 2016, modified from Wu and Sues, 1996).
206. Mandibular symphysis, shape, in ventral view: tapering mediolaterally anteriorly, forming an angle (0); U-shaped, smoothly curving anteriorly (1); lateral edges anteroposteriorly oriented with convex anterolateral cornier and extensive, transversely oriented anterior edge (2) (Pol, 1999a).
207. External mandibular fenestra: absent (0); present (1) (Clark, 1994).
208. External mandibular fenestra, size: present as a diminutive passage, less than 50% of the total size of the lateral temporal fenestra (0); present as an evident fenestra, 50% or greater than the total size of the lateral temporal fenestra (1) (Tennant et al., 2016, modified from Clark, 1994).
209. External mandibular fenestra, orientation of main axis: horizontal to sub-horizontal (0); inclined, directed anteroventrally-posterodorsally (Tennant et al., 2016, modified from Andrade *et al*., 2011).
210. External mandibular fenestra, shape: subcircular to elliptical (0); triangular (1) (Tennant et al., 2016, modified from Andrade *et al*., 2011).
211. Jaw joint, location of dorsal edge: level with or dorsolateral to occipital condyle (0); ventrolateral occipital condyle (1) (Wu and Sues, 1996).
212. Dentary, lateral surface adjacent to seventh alveolus: smooth (0); with lateral concavity for reception of enlarged maxillary tooth (1) (Buckley and Brochu, 1999).
213. Dentary, lateral surface below alveolar margin, at middle to posterior region of tooth row: vertically oriented, continuous with rest of lateral surface of the dentaries (0); flat surface exposed dorsolaterally, divided by ridge from the rest of the lateral surface of the dentary (1); flat, unsculpted surface confluent with rest of the lateral surface (2) (Tennant et al., 2016, modified from Pol and Apesteguía, 2005).
214. Dentary, relative to external mandibular fenestra: extends posteriorly beneath mandibular fenestra, posteriorly exceeding anterior margin (0); does not extend beneath fenestra, either terminating anteriorly to fenestra or only forming a point contact (1) (Clark, 1994).
215. Dentary, mediolateral compression and ventrolateral surface anterior to mandibular fenestra (or of anterior portion posterior to symphysis if fenestra is absent): compressed and flat (0); uncompressed and convex (1) (Tennant et al., 2016, modified from Ortega *et al*., 1996).
216. Dentary, sculpted below the tooth row: lacking sculpting (0); present (1) (Pol, 1999a).
217. Dentary alveoli: all independent of one another (0); some confluent (1); all confluent, within continuous alveolar groove (2) (Tennant et al., 2016) [**ordered**].
218. Dental alveoli, transitional shape morphology from circular to sub-circular or oval: absent (0); present (1) (Tennant et al., 2016).
219. Dentary alveoli, transitional shape morphology: posteriorly from D4 (0); posteriorly from D5 (1) (Tennant et al., 2016).
220. Dentary, distinct foramina on occlusal surface, lingual to dental arcade: absent (0); present (1) (Tennant et al., 2016).
221. Dentary, distinct foramina on occlusal surface, lingual to dental arcade: at D2-D3 (0); at D4 or positioned more posteriorly (1) (Tennant et al., 2016).
222. Dentary, external alveolar margins, dorsal edge: vertically festooned, forming raised rims about each alveolus (0); flat (1) (Tennant et al., 2016).
223. Dentary, internal alveolar margins: forming raised rims (0); flat and confluent with dentary occlusal surface (1) (Tennant et al., 2016).
224. Dentary, anterior portion, lateral margin shape in dorsoventral view: straight (0); distinctly spatulate, with abrupt lateral expansion (1); laterally convex (2) (Tennant et al., 2016).
225. Dentary, diastema (gap) between D7 and D8: present (0); absent (1) (Tennant et al., 2016).
226. Dentary, pitted ornamentation of external surface: absent (0); present (1) (Tennant et al., 2016).
227. Dentary, grooved ornamentation of external surface: absent (0); present (1) (Tennant et al., 2016).
228. Dentary, inter-alveolar septae within anterior dental arcade (D4-D8): present (0); absent (1) (Tennant et al., 2016).
229. Dentary, symphysis and dentary arcade lateral to symphysis, in dorsoventral view: parallel (0); oblique (1) (Tennant et al., 2016).
230. Dentary, occlusal surface: smooth (0); anteroposteriorly crenulated (1) (Tennant et al., 2016).
231. Dentary, obliquely inclined crenulations posterodorsal to D8-D9: present (0); absent (1) (Tennant et al., 2016).
232. Dentary, dorsolateral edge: slightly concave or straight and subparallel to anteroposterior axis of skull (0); sinusoidal, with two concave waves (1) (Tennant et al., 2016, modified from Ortega *et al*., 1996).
233. Splenial, involvement in symphysis, in ventral view: not involved (0); involved (1) (Clark, 1994)
234. Splenial, contact with dentary, in ventral view: confluent (0); dorsally inset (1) (Tennant et al., 2016).
235. Splenial, posterior to symphysis: approximately constant mediolateral thickness throughout element (0); more robust posterodorsally (1) (Tennant et al., 2016, modified from Ortega *et al*., 1996).
236. Angular and posterior surangular, strong pitted pattern: absent (0); present (1); lateral surface with rugose pattern instead of pits (2) (Tennant et al., 2016, modified from Andrade *et al*., 2011).
237. Surangular, dorsal edge in lateral view: mostly straight (0); arched dorsally, excluding articular projection (1) (Clark, 1994).
238. Surangular, anteroposterior ridge along the dorsolateral surface: absent (0); present (1) (Pol and Norell, 2004b).
239. Surangular, extension toward posterior end of retroarticular process: along entire length (0); pinched off anterior to posterior tip (Norell, 1988).
240. Articular, posterior ridge on glenoid fossa: posterior margin well-developed, evidently high (0); posterior margin poorly delimited, crest absent (1) (Pol and Apesteguía, 2005).
241. Articular, medial process articulating with otoccipital and basisphenoid: absent (0); present (1) (Clark, 1994).
242. Retroarticular process: absent or extremely reduced (0); posteroventrally projecting and paddle-shaped (1); pointed, projects posteriorly and ventrally recurved (2); projects posteriorly and dorsally recurved (3) (Tennant et al., 2016, modified from Clark, 1994).

**Dental Characters**

243. Premaxillary teeth, number: five or more (0); four or fewer (1) (Tennant et al., 2016, modified from Wu and Sues, 1996).
244. Posterior premaxillary teeth, apicobasal length: less than 1.5 times the size of the anterior teeth (0); 1.5 times or greater than anterior teeth (1) (Tennant et al., 2016, modified from Clark, 1994).
245. Maxillary teeth, mesiodistal margin carinae: absent or with smooth and crenulated carinae (0); with denticulate carinae (ziphodont condition) (1) (Tennant et al., 2016, modified from Ortega *et al*., 1996).
246. Maxillary tooth rows, middle to posterior elements: crowns not mesiodistally compressed, subcircular in cross-section (0); crowns slightly compressed mesiodistally (mesiodistal to labiolingual diameter ratio more than 0.5 at mid-height) (1); crowns highly compressed mesiodistally (mesiodistal to labiolingual diameter ratio less than or equal to 0.5 at mid-height) (2) (Pol, 1999a) [**ordered**].
247. Maxillary tooth rows, mesiodistal compression of middle to posterior elements: absent, or symmetrical compression (0); asymmetrical compression, occurring only along the distal margin giving teeth a teardrop shape (1) (Tennant et al., 2016, modified from Andrade and Bertini, 2008).
248. Maxillary teeth, middle to posterior elements, ridged ornamentation on enamel surface: absent (0); present (1) (Andrade *et al*., 2011).
249. Maxillary teeth, enamel surface: smooth or slightly crenulated (0); with ridges at base of crown (often extending apically) (1) (Turner and Sertich, 2010).
250. Maxillary teeth, striations on labial and lingual faces: present (0); absent (1) (Tennant et al., 2016, adapted from Martin *et al*., 2014).
251. Cheek teeth, base (i.e., immediately apical to root), with respect to remainder of tooth crown: not constricted (0); constricted (1) (Tennant et al., 2016, adapted from Martin *et al*., 2014)
252. Maxillary teeth, width of root with respect to crown: narrower (0); wider in anterior teeth and equal in posterior teeth (1) (Tennant et al., 2016, modified from Ortega *et al*., 2000).
253. Maxillary teeth, posterior teeth, mediolaterally compressed lanceolate-shaped morphotype (sometimes called ‘leaf-shaped’), visible in labial or lingual view, with wide crown tapering apically to a sharp point (note that the point can often be abraded): present (0); absent (1) (Tennant et al., 2016, adapted from Schwarz and Salisbury, 2005).
254. Maxillary teeth, low-crowned and strongly labiolingually compressed morphotypes, forming a crown that is mesiodistally broader than it is apicobasally tall: present, apical margins oriented at less than 45° from horizontal (0); absent (1) (Tennant et al., 2016, adapted from Schwarz and Salisbury, 2005).
255. Tooth, present at premaxilla-maxilla contact with transitional size-based morphology: absent (0); present (1) (Tennant et al., 2016, modified from Turner and Sertich, 2010).
256. Maxillary teeth, size variation waves: absent, no tooth size variation (0); one wave of enlarged teeth (1); enlarged maxillary teeth occur in two waves (festooned) (2) (Clark, 1994).
257. Enlarged maxillary teeth (at least 1.5 times the apicobasal size of remaining teeth): present at M2 and/or M3 (0); present at M4 and/or M5 (1) (Martin *et al*., 2014).
258. Maxillary tooth 5, apicobasal size relative to adjacent maxillary teeth: subequal, or less than 4.0 times the size of adjacent teeth (0); hypertrophied, at least 4.0 times the size of adjacent teeth (1) (Tennant et al., 2016).
259. Maxillary tooth 5, hypertrophied: directed posteroventrally (0); directed ventrally (1) (Tennant et al., 2016).
260. Maxillary teeth 6 and 7: continuous with tooth row (0); dorsally inset (1) (Tennant et al., 2016).
261. Maxillary teeth, bulbous tooth morphotype (tribodont): present (0); absent (1) (Sweetman *et al*., 2015).
262. Dentary teeth, anterior teeth (opposite premaxilla-maxilla contact) apicobasal length, relative to rest of dentary teeth: no more than twice the length (0); more than twice the length (1) (Clark, 1994).
263. Dentary teeth, posterior teeth: occlude medial to opposing maxillary teeth (0); occlude lateral to, or interlock with, opposing maxillary teeth (1) (Tennant et al., 2016, adapted from Sweetman et al 2015)

**Axial Characters**

264. Vertebrae, centra shape along axial column: cylindrical throughout (0); grade continuously from cylindrical to elongated spool-shaped (1); spool-shaped throughout (2) (Tennant et al., 2016, modified from Buscalioni and Sanz, 1988) [**ordered**].
265. Cervical vertebrae: amphicoelous or amphiplatyan (0); procoelous, and posterior centrum face (condyle) with a central depression (‘semi-procoely’) (1); fully procoelous (2) (Tennant et al., 2016, modified from Clark, 1994) [**ordered**].
266. Cervical vertebrae, number: 6 or fewer (0); 7 (1); 8 or more (2) (Tennant et al., 2016) [**ordered**].
267. Atlas, intercentrum size: mediolaterally wider than anteroposteriorly long (0); subequal diameters or anteroposteriorly longer (Clark, 1994).
268. Cervical vertebrae, neural spine: absent, or extremely reduced (0); present, distinct from centrum body (1) (Tennant et al., 2016).
269. Cervical vertebrae, neural spines: rod-like and elongate (0); short and transversely flattened (1) (Tennant et al., 2016).
270. Cervical and dorsal vertebrae, hypapophyses or anterior keels: absent (0); present (1) (Tennant et al., 2016, modified from Buscalioni and Sanz, 1988).
271. Dorsal vertebrae: amphicoelous or amphiplatyan (0); procoelous (1) (Tennant et al., 2016, modified from Clark, 1994).
272. Dorsal vertebrae, number: 14 or fewer (0); 15-16 (1); 17 or more (2) (Tennant et al., 2016) [**ordered**].
273. Posterior dorsal vertebrae, transverse process shape: dorsoventrally low and laminar (0); dorsoventrally high (1) (Buscalioni and Sanz, 1988).
274. Sacral vertebrae, number: two (0); three or more (1) (Buscalioni and Sanz, 1988).
275. Sacral vertebrae, orientation of transverse processes: project laterally (horizontally) (0); deflected markedly ventrally (1) (Gasparini *et al*., 2006).
276. Caudal vertebrae, number: less than 50 (0); 50 or more (1) (Tennant et al., 2016).
277. Caudal vertebrae: amphicoelous or amphiplatyan (0); procoelous (1) (Tennant et al., 2016, adapted from Salisbury and Frey, 2005).
278. Caudal vertebrae, first: same morphology as rest of caudal series (0); biconvex (1) (Tennant et al., 2016, adapted from Salisbury and Frey, 2001).
279. Caudal vertebrae, anteroposterior ridge/lamina separating centrum and neural arch: present (0); absent (1) (Tennant et al., 2016; note that this could be an ontogenetic feature).

**Appendicular Characters**

280. Scapula, proximodorsal edge in lateral view: flat and confluent with scapular shaft (0); forms a distinct crest (1) (Tennant et al., 2016).
281. Coracoid, medial process: elongate posteromedial process (0); distally expanded ventromedial process (1) (Wu and Sues, 1996).
282. Coracoid, distal expansion: equal to or larger than the proximal expansion (0); less expanded than the proximal region (1) (Pol *et al*., 2012).
283. Humerus, circular depression on the posterior surface of the proximal end, for the insertion of the *M. scapulohumeralis caudalis*: absent (0); present (1) (Pol *et al*., 2012).
284. Humerus, lateral and medial surfaces of distal end: flat and anteroposteriorly broad, similar in anteroposterior length to the transverse width of the distal end of the humerus (0); convex and reduced in comparison with the transverse width of the distal humerus (1) (Pol *et al*., 2012).
285. Forelimb:hindlimb length, ratio: less than 0.7 (0); 0.7 to less than 0.8 (1); 0.8 or greater (2) (Tennant et al., 2016) [**ordered**].
286. Humerus:femur length, ratio: less than 0.75 (0); 0.75 to less than 1.0 (1); 1.0 or greater (2) (Tennant et al., 2016) [**ordered**].
287. Ulna, morphology of olecranon process: narrow and subangular (0); wide and rounded (1) (Brochu, 1999).
288. Radius:humerus length, ratio: less than 0.6 (0); 0.6 to less than 0.75 (1); 0.75 or greater (2) (Tennant et al., 2016) [**ordered**].
289. Radius:tibia length, ratio: less than 0.6 (0); 0.6 to less than 0.7 (1); 0.7 or greater (2) (Tennant et al., 2016) [**ordered**].
290. Radiale, proximal end, shape: expanded symmetrically, similar to distal end (0); more expanded laterally than medially (“hatchet shaped”) (1) (Buscalioni and Sanz, 1988).
291. Ilium, anterior (preacetabular) process, length relative to posterior (postacetabular) process: greater than 75% the length of the posterior process (0); 75% or less the length of the posterior process (1); completely absent (2) (Tennant et al., 2016, modified from Clark, 1994) [**ordered**].
292. Ilium, development of the posterior (postacetabular) process: well-developed as a distinct process that extends anteroposteriorly for 60% or more of the acetabular length (0); extremely reduced or absent, extending anteroposteriorly less than 60% of the acetabular length (1) (Tennant et al., 2016, modified from Pol *et al.*, 2012).
293. Ilium, posterior end of the postacetabular process: tapering posteriorly to an acute tip (0); subrectangular with a vertically oriented posterior margin (1) (Tennant et al., 2016, modified from Pol *et al*., 2012).
294. Pubis, shape: rod-like without expanded distal end (0); with anterodorsally-posteroventrally expanded distal end (1) (Tennant et al., 2016, modified from Clark, 1994).
295. Pubis, anterior process: absent (0); present (1) (Clark, 1994).
296. Femur, proximal development of greater trochanter: prominent, ridge-like lateral border that separates the lateral surface of the proximal femur from a flat posterior surface reaching down to the level of the fourth trochanter (0); proximodistally short trochanteric surface lacking a distinct ridge, terminating well above the fourth trochanter (1) (Pol *et al*., 2012).
297. Femur, femoral head: mediolaterally flattened (0); hemispherical (1) (Tennant et al., 2016).
298. Tibia, distal projection of articular surfaces: medial region of distal articular surface extends further distally than the lateral region, forming a strongly oblique distal margin of the tibia (0); medial and lateral regions sub-equally extended, with distal margin sub-horizontally oriented (1) (Pol *et al*., 2012).
299. Tibia, posterior surface of shaft: flattened and confluent with fibula (0); twists posteriorly, leaving a void between the tibia and fibula (1) (Tennant et al., 2016).
300. Tibia:femur length, ratio: less than 0.9 (0); 0.9 to less than 1.0 (1); 1.0 or greater (2) (Tennant et al., 2016) [**ordered**].
301. Astragalus, anterior margin of the tibial facet: forming a well-defined ridge that reaches medially the ball-shaped region for the articulation of metatarsal I-II and closes the proximomedial corner of the anterior hollow of the astragalus (0); forming a low ridge that is medially separated by a notch from the ball-shaped region for the articulation of the metatarsals I-II, failing to close the proximomedial corner of the anterior hollow (1) (Pol *et al*., 2012).
302. Distal tarsals, digits 2-4, dorsal surface: longitudinally grooved (0); smooth and flat (1) (Tennant et al., 2016).
303. Metatarsals I-IV: equidimensional (0); metatarsal I shorter than metatarsals II-IV (1) (Tennant et al., 2016).

**Osteoderm Characters**

304. Osteoderms, dorsal surface: entirely sculpted (0); partially or completely unsculpted (1) (Tennant et al., 2016).
305. Presacral armour: cervical and dorsal trunk shields undifferentiated, morphology grading continuously (0); cervical shields clearly differentiated from dorsa trunk shields by size and general morphology (regardless of contact between nuchal and trunk series) (1); anteriormost cervical osteoderms developed into distinct shield (2) (Andrade *et al*., 2011) (character state 2 added) [**ordered**].
306. Nuchal osteoderms: consistent morphology along series (0); vary substantially in size in a random fashion (1); systematically increase in size posteriorly (2) (Tennant et al., 2016).
307. Nuchal osteoderms, with size variation: nuchals no less than 50% the size of dorsal osteoderms (0); some smaller than one half of the size of the dorsal osteoderms (1) (Tennant et al., 2016).
308. Dorsal osteoderms, shape: rounded or ovate (0); subrectangular (mediolaterally wider than anteroposteriorly long) (1); subtriangular (2); square (3) (Tennant et al., 2016, modified from Clark, 1994).
309. Dorsal osteoderms, articular anterior process: absent (0); present (1) (Clark, 1994).
310. Dorsal osteoderms, articular anterior process: as discrete convexity on anterior margin (0); well-developed process located anterolaterally (‘peg and socket’ articulation) (1) (Clark, 1994).
311. Dorsal osteoderms, anteroposterior keel on anterior part of dorsal surface: absent (0); present (1) (Tennant et al., 2016).
312. Dorsal osteoderms, anteroposterior keel on posterior part of dorsal surface: absent (0); present (1) (Tennant et al., 2016).
313. Dorsal and cervical osteoderms: some or all imbricated (0); not in contact (1) (Tennant et al., 2016).
314. Dorsal osteoderms, sutured anterior and posterior contacts: present (0); absent (1) (Tennant et al., 2016).
315. Dorsal primary osteoderms (sensu Frey, 1988), rows: two parallel rows (0); four rows or more (1) (Tennant et al., 2016, modified from Clark, 1994).
316. Dorsal osteoderms, accessory osteoderms (sensu Frey, 1988; i.e., osteoderms not forming part of the dorsal shield): absent (0); present (1) (Turner and Sertich, 2010).
317. Dorsal osteoderms, dorsal keel: same morphology in anteriormost dorsal osteoderms as remainder of dorsal series (0); keel shifts laterally in more posterior dorsal osteoderms (1) (Tennant et al., 2016).
318. Dorsal osteoderms, anterior edge of dorsal surface (i.e., articular surface, if present): sculpted, undifferentiated from main osteoderm body (0); unsculpted (1) (Tennant et al., 2016).
319. Dorsal osteoderms, outline in dorsal aspect (excluding peg articulation): symmetrical about anteroposterior axis (0); asymmetrical (1) (Tennant et al., 2016).
320. Dorsal osteoderms, mediolateral contacts: contact but not sutured (0); sutured (1) (Tennant et al., 2016).
321. Dorsal osteoderms, ventral to dorsal vertebrae beneath trunk: absent (0); present (1) (Clark, 1994).
322. Caudal osteoderms: absent (0) present on dorsal surface only (1); completely surrounding tail (2) (Clark, 1994) [**ordered**].
323. Caudal osteoderms: ovate (0); subcircular (1); subrectangular (2) (Tennant et al., 2016).
324. Caudal osteoderms, bearing anteroposterior ridge: present (0); absent (1) (Tennant et al., 2016).
325. Caudal osteoderms, anteroposterior ridge: present medially (0); forms a distinct lateral step in posterior-most elements (1) (Tennant et al., 2016)
326. Caudal osteoderms, geometry: continuous from short to elongate oval (0); continuous from subrectangular (rounded corners) to suboval (1); isometric (equal geometry along series) (2) (Tennant et al., 2016).
327. Caudal osteoderms, medial and lateral edges: serrated (0); smooth (1) (Tennant et al., 2016).
328. Caudal osteoderms, secondary osteoderms: present (0); absent (1) (Tennant et al., 2016).
329. Caudal osteoderms, anteroposterior ridges: same morphology along series (0); becoming more pronounced posteriorly, coincident with a decrease in osteoderm size (1) (Tennant et al., 2016).
