## Supplementary figures and images for "Durophagous adaptations in a new crocodyliform from the Santonian of Hungary and a systematic reassessment of European Cretaceous atoposaurids"

### Figure_1.jpg

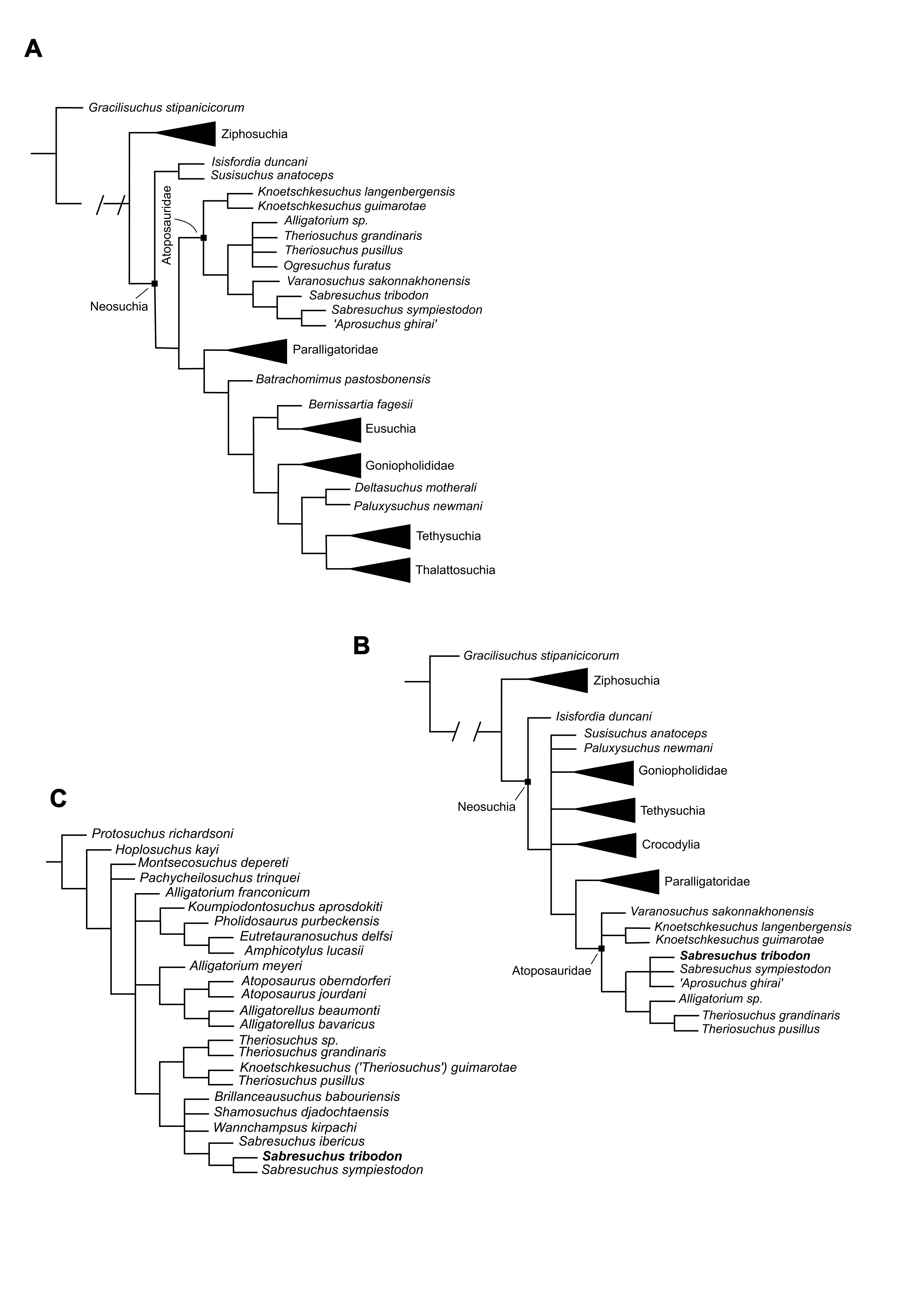

### Figure_2.jpg

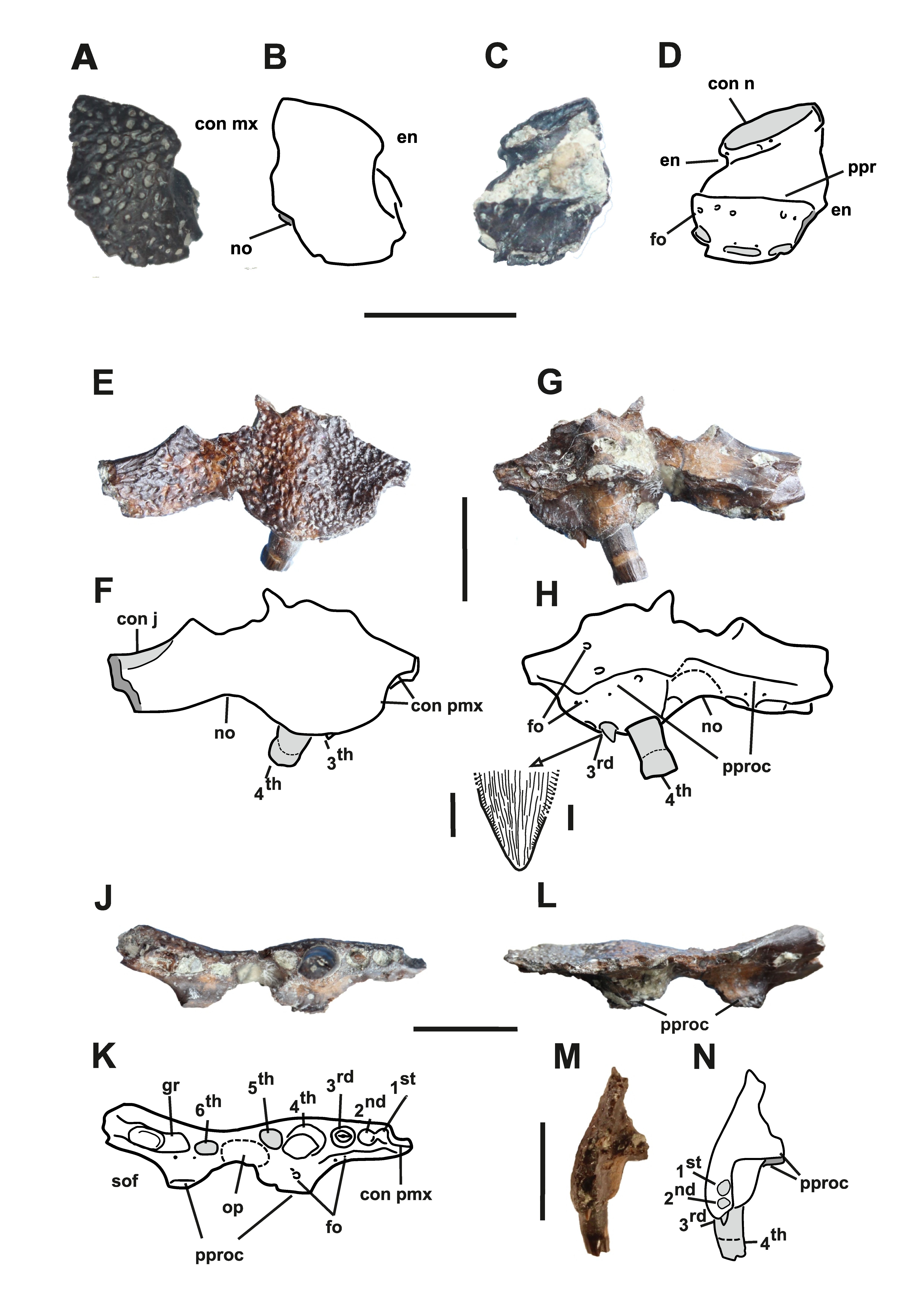

### Figure_3.jpg

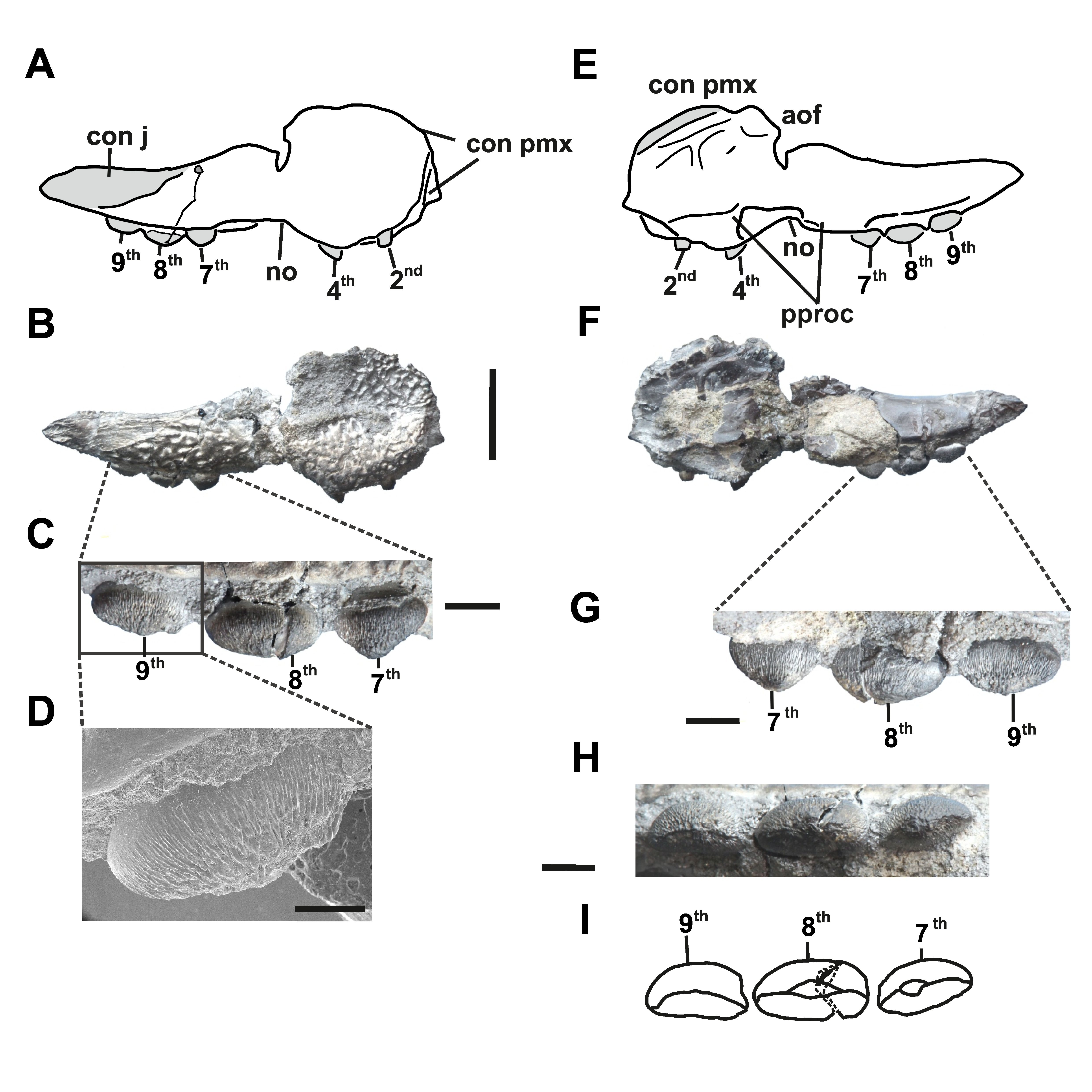

### Figure_4.jpg

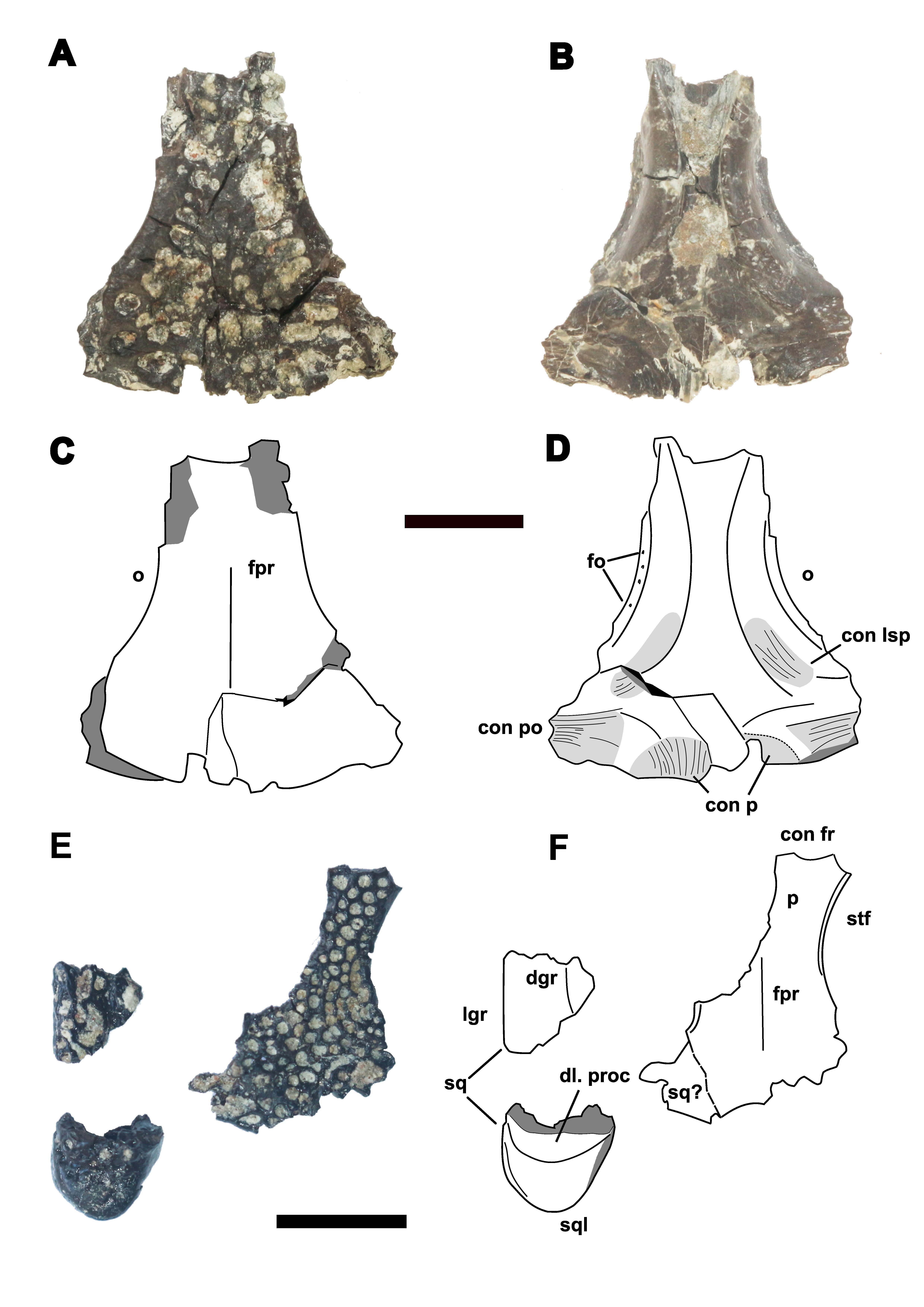

### Figure_5.jpg

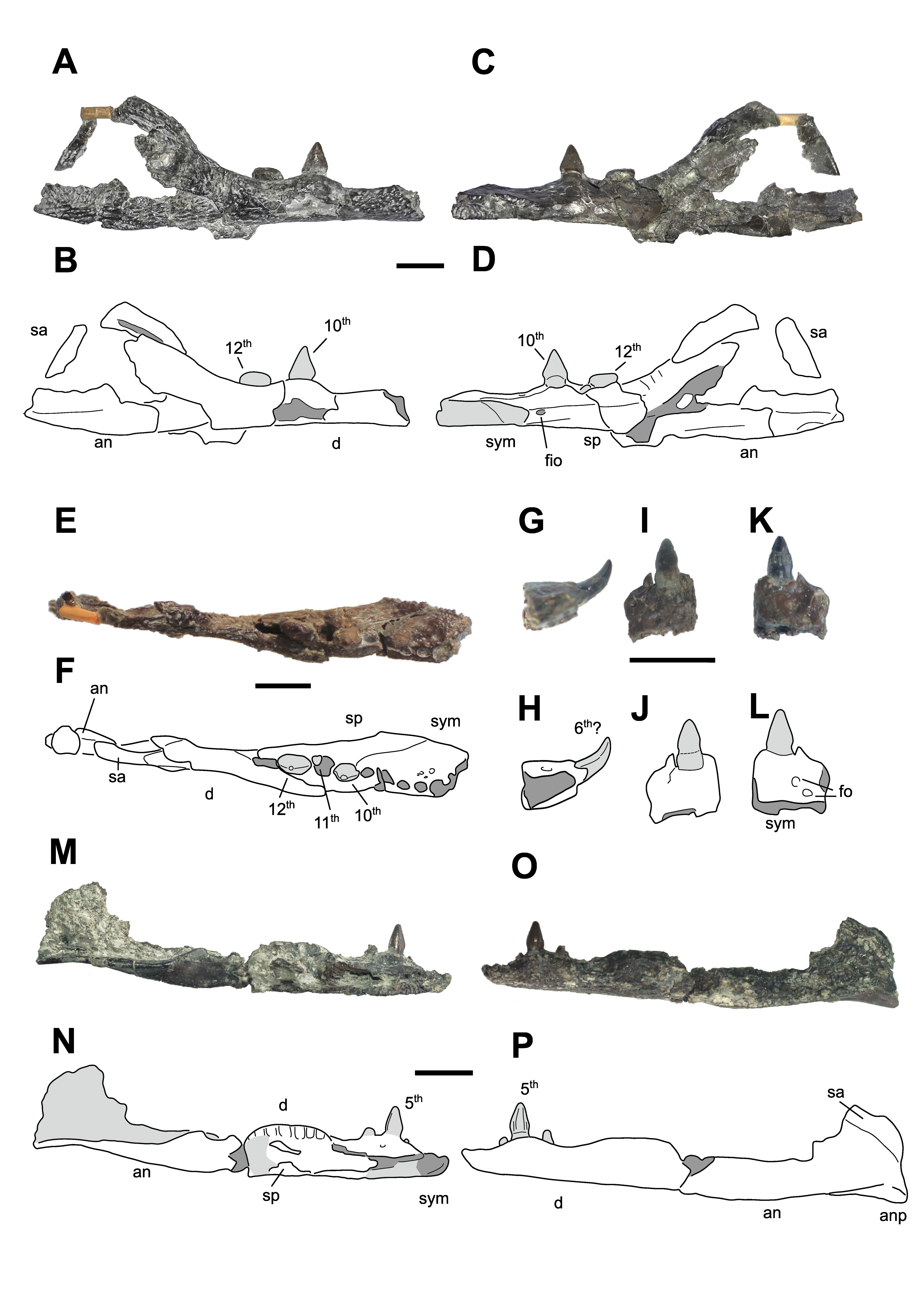

### Figure_6.jpg

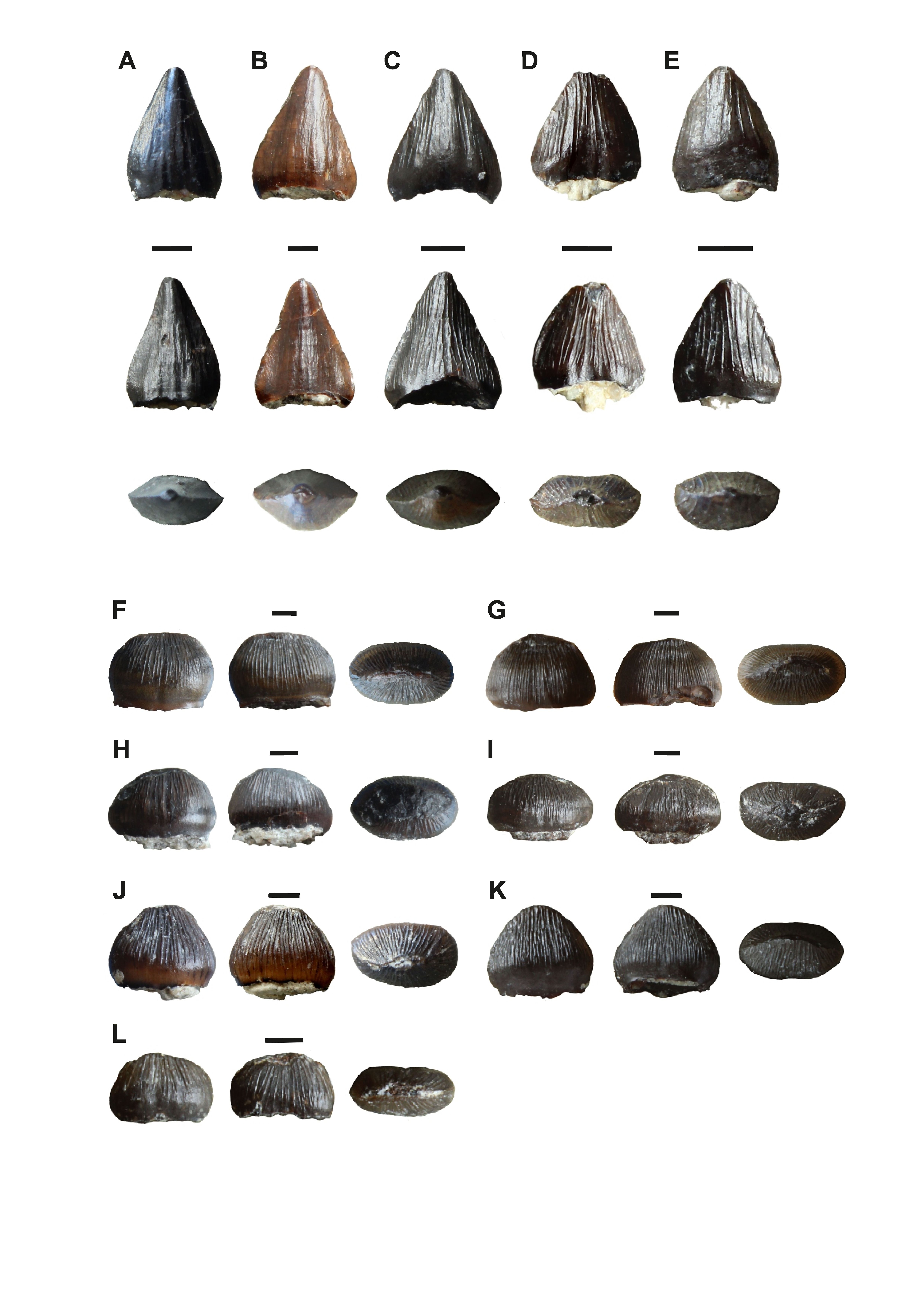

### Figure_7.jpg

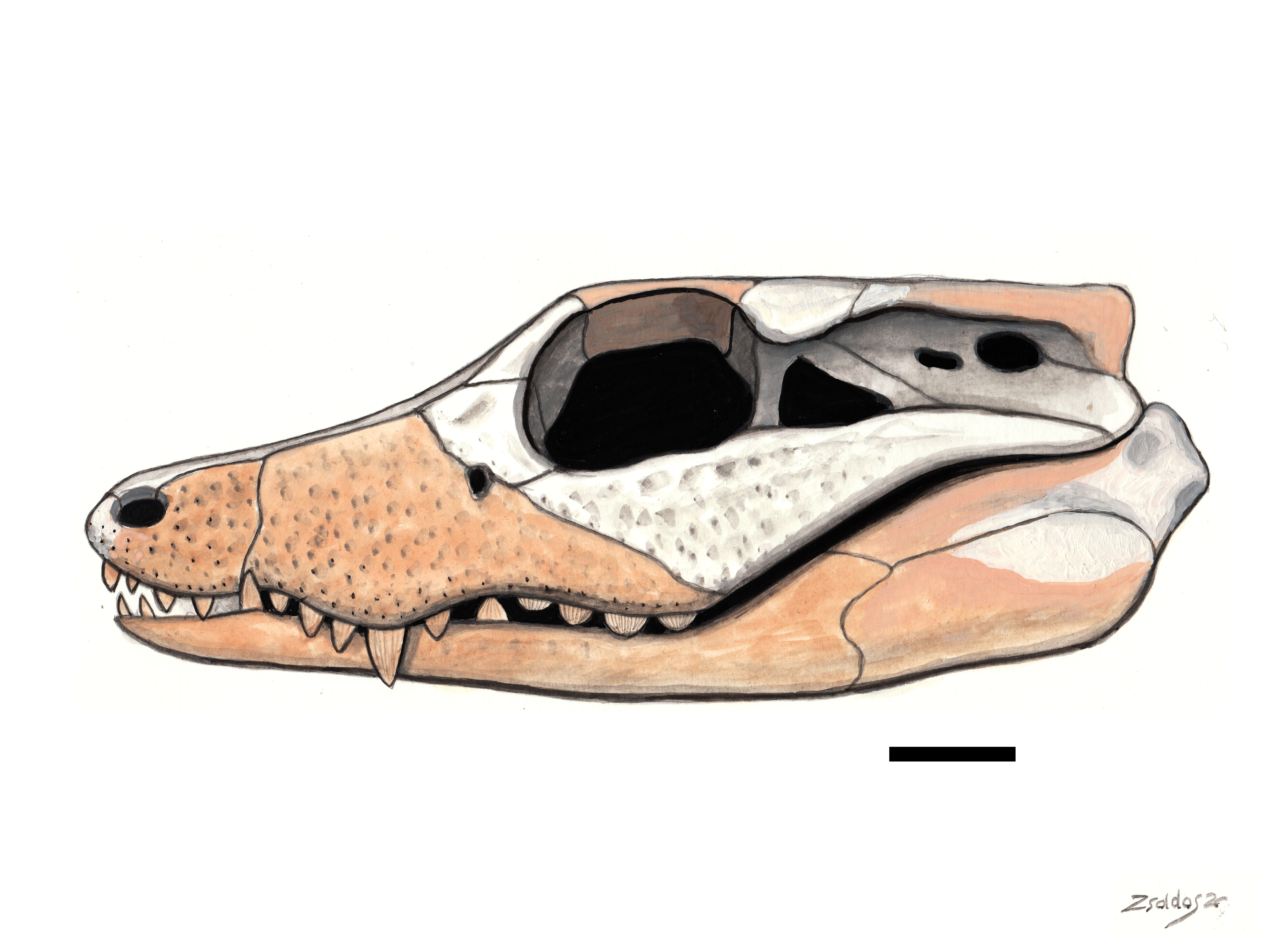

### Figure_8.jpg

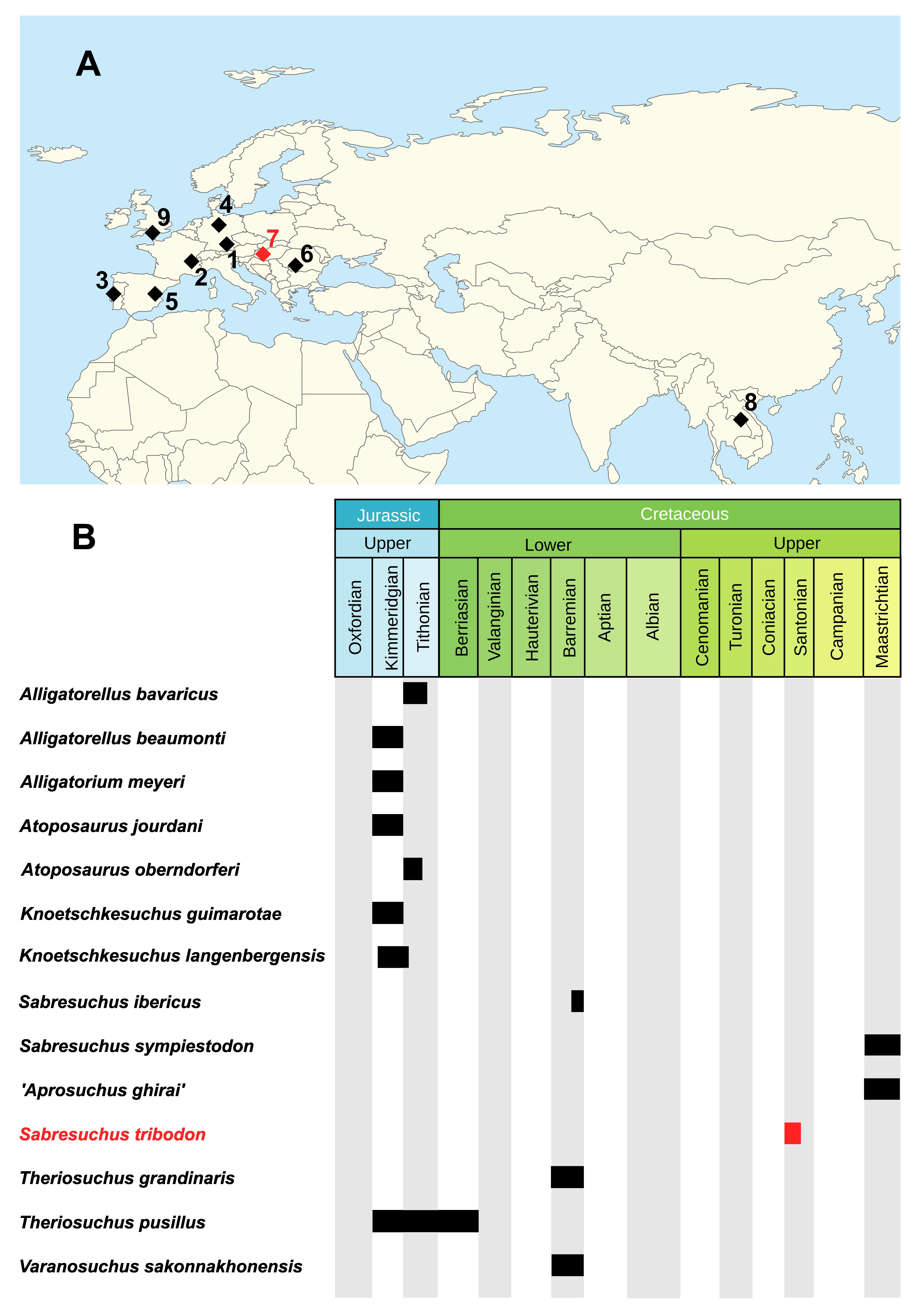
